# Bifunctional probes reveal the rules of intracellular ether lipid transport

**DOI:** 10.1101/2024.07.26.605283

**Authors:** Kristin Böhlig, Juan M. Iglesias-Artola, H. Mathilda Lennartz, Anna C. Link, Björn Drobot, André Nadler

## Abstract

Ether glycerophospholipids bear a long chain alcohol attached via an alkyl or vinyl ether bond at the *sn1* position of the glycerol backbone. Emerging evidence suggests that ether lipids play a significant role in physiology and human health but their precise cellular functions remain largely unknown. Here, we introduce bifunctional ether lipid probes bearing diazirine and alkyne groups to study ether lipid biology. To interrogate the kinetics of intracellular ether lipid transport in mammalian cells we used a combination of fluorescence imaging, machine learning-assisted image analysis and mathematical modelling. We find that alkyl-linked ether lipids are transported up to twofold faster than vinyl-linked plasmalogens, suggesting that the lipid transport machinery can distinguish between linkage types differing by as little as two hydrogen atoms. We find that ether lipid transport predominantly occurs via non-vesicular pathways, with varying contributions from vesicular mechanisms between cell types. Altogether, our results suggest that differential recognition of alkyl- and vinyl ether lipids by lipid transfer proteins contributes to their distinct biological functions. In the future, the probes reported here will enable studying ether lipid biology in much greater detail through identification of interacting proteins and in-depth characterization of intracellular ether lipid dynamics.

## Main

Eukaryotic ether lipids are primarily phosphatidylcholines (PCs) and phosphatidylethanolamines (PEs) featuring a long-chain alcohol attached via alkyl (plamanyl) or vinyl (plasmenyl) ether bonds at the *sn1* position of the glycerol scaffold.[1] Between 10-20% of all glycerophospholipids in humans are ether lipids, with increased levels in brain, heart, kidney and skeletal muscle tissues.[1] Ether lipids are thought to have specific physiological roles despite the rather small structural deviation from the corresponding diacyl ester lipids. For example, disruption of peroxisomal ether lipid biosynthesis causes severe hereditary diseases, which are characterized by skeletal, renal and cerebral abnormalities.[2] Corresponding mice models show abnormal eye development, male infertility and irregular behavior.[3–6] Many studies also indicate a correlation between disturbed ether lipid metabolism and neurodegenerative diseases, psychiatric disorders and cancer but the underlying molecular mechanisms are unclear.[1,7–13]

Our knowledge of the cell biology of ether lipids is derived primarily from indirect evidence. Ether lipids have been proposed to play a role in membrane trafficking[14], the general architecture of the endomembrane system, lipid sorting in the cell, neurotransmitter release[15] and ferroptosis.[16–19] Intriguingly, elevated levels of ether lipids were recently shown to be an adaptation of deep sea organisms to the high pressure environment.[20] Perhaps the clearest evidence points to a role of ether lipids in modulating the export of GPI-anchored proteins via a direct interaction with the transmembrane domain of mammalian p24 proteins, being metabolically co-regulated with sphingolipids.[21] Recent progress in ether lipid biology was mostly made by combining genetic or other perturbations such as oxidative stress with lipidomic approaches that now allow to distinguish between plasmenyl and plasmanyl lipids.[22] So far, very few ether lipid functions have been studied in mechanistic detail in biological assays. Thus, the possibility remains that the wide range of observed phenotypes are partially second-order effects of genetic perturbations. The difficulties associated with mechanistic investigations into cellular functions of ether lipids can be explained by an almost complete lack of suitable chemical tools for studying ether lipids in situ. Apart from ether lipid analogues labelled with bulky fluorescent moieties[23,24] which are of limited use in cellular assays, the most promising approach has so far been the use of polyene lyso-ether lipids, which are intrinsically fluorescent due to conjugated trans double bonds.[25,26] Despite promising early results, their use has been limited, likely due to the relative complexity of the synthetic routes and issues with rapid photobleaching during microscopy. Further development of strategies for directly monitoring intracellular ether lipid localization, trafficking and interaction partners are needed.

Among the most versatile tools for studying lipid biology are bifunctional probes equipped with photoactivatable (diazirine) and clickable (alkyne) moieties. After UV-induced crosslinking to proximal proteins, reporter groups can be attached to the alkyne group by copper-mediated click chemistry. Bifunctional lipids have been used to study lipid-protein interactions of fatty acyls, regular (diacyl) glycerophospholipids, sphingolipids and sterols by utilizing affinity tags (e.g. biotin) and subsequent pulldown of lipid-protein conjugates generated by crosslinking.[27–34] In a limited number of examples, these probes have also been used to report on lipid localization and transport by attaching fluorophores after crosslinking and cell fixation and subsequent analysis by fluorescence microscopy.[29,34,35] Recently, we expanded upon this methodology by employing bifunctional phospholipids to quantify both lipid transport and metabolism using a combination of ultra-high resolution mass spectrometry and fluorescence imaging.[36] We reasoned that bifunctional ether lipid probes would be ideal tools to study ether lipid biology.

As the primary probe of this study, we synthesized a PC ether lipid (1) featuring a bifunctional, alkyl-ether-linked *sn1* chain in a 15-step synthesis featuring a stereoselective Sc(OTf)_3_-mediated epoxide opening. A complementary set of bifunctional phosphatidylcholines (2-5) that only differ in the *sn1* linkage type via ester, alkyl ether or a vinyl ether bond was generated to enable comparative studies of lipid transport in cells. We performed pulse-chase experiments in two different cell lines (HCT-116 and U2OS) and found that the plasmalogen probe was consistently transported slower compared to both alkyl-ether and ester lipid probes. Ether lipid transport was found to be primarily non-vesicular in both investigated cell lines, but the fraction of material transported through the vesicular route was higher in U2OS vs HCT-116 cells. Our data suggest that lipid transfer proteins distinguish plasmanyl and plasmenyl ether lipids, suggesting a plausible mechanism for the enrichment of distinct ether lipids in cellular membranes.

## Results

### Synthesis of bifunctional ether lipid probes

To study the influence of the linkage type on lipid transport, we synthesized bifunctional lipids with identical head groups and closely related side chain compositions. We focused on PC species as they are among the most abundant ether lipids in mammalian cells.[37] As plasmalogens feature one unsaturation in the *sn1* chain we decided to use the monounsaturated oleic acid for the corresponding ester and alkyl ether version. Probes **1** and **2** were designed to assess the influence of alkyl ether vs ester linkage at the *sn1* position (Fig. 1a), as they are otherwise identical molecules. A complementary set of probes (**3**-**5**) was designed to perform a three-way comparison of alkyl-ether, vinyl-ether and ester linkages (Fig. 1a). Together, the complete set of probes was designed to study the effects of the following structural elements that can plausibly affect transport mechanisms of ether lipids in cells: (i) side chain linkage types (alkyl ether, vinyl ether, ester) and (ii) positioning of a single double bond (*sn1*/*sn2* chain, Δ9/Δ1).

The synthesis of the bifunctional lipids required two different strategies. For molecules 2-5, we first synthesized bifunctional fatty acids according to our recently published protocol (see details in SI)[36]. We then coupled the bifunctional fatty acids to the respective lyso-lipids via EDC/DMAP-mediated esterification (see SI for details). Probe 1, which bears the diazirine and alkyne functionalities in the ether chain at the *sn1* position, required a 15-step synthesis (Fig. 1b, see SI for synthetic details and analytical data). We chose a synthetic route via the 1-Alkyl-2-acylglycerol ether 17, as this intermediate offers straightforward access to a range of other ether lipid classes via phosphoramidite chemistry.

**Figure 1.**
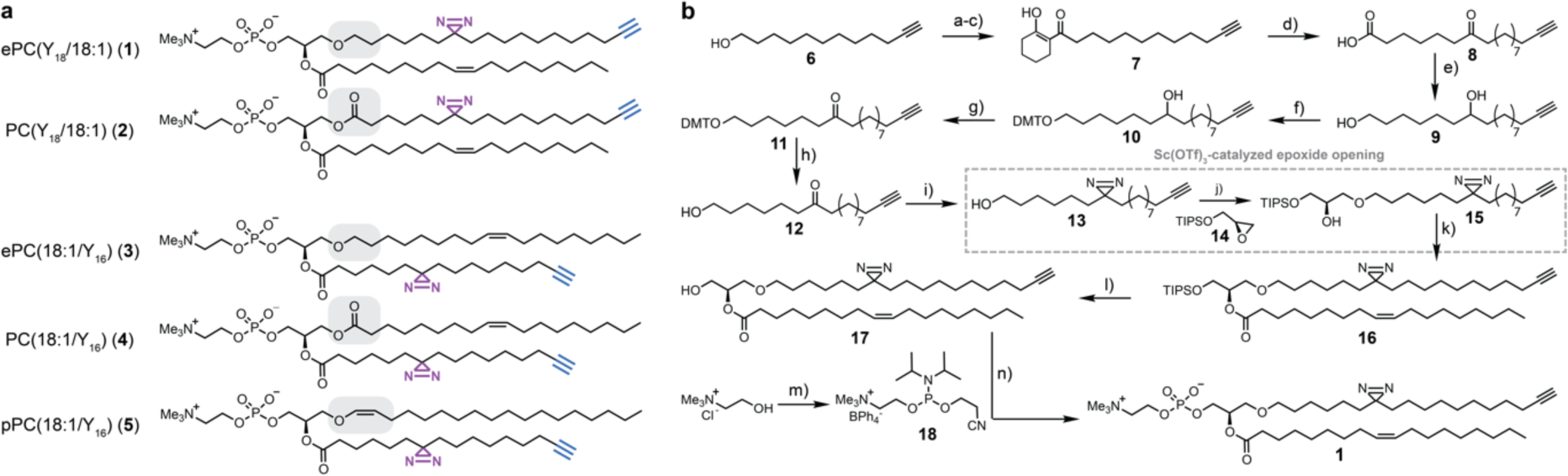
Overview of bifunctional ether lipid probes and synthesis of ePC(Y_18_/18:1) bearing a bifunctional ether linked side chain. **a**, Structures of synthesized bifunctional ether and ester PCs 1-5 differing in the linkage type at the *sn1* position. **b,** Synthesis of the bifunctional ether lipid 1. The bifunctional alcohol 13 was synthesized in 8 steps starting from 11-Dodecynol (6) and coupled to the TIPS-protected (S)-glycidol 14. After oleic acid coupling and deprotection, phosphorylation according to Xu et. al.[38] gave the final molecule 1. Reaction conditions: a) CrO_3_, H_2_SO_4_, H_2_O, 0 °C-RT, 1h, 85%; b) (COCl)_2_, DMF, DCM, 1 h, 0 °C-RT; c) 1-morpholinocyclohexene, NEt_3_, DCM, O.N., RT, 76% over 2 steps; d) aq. KOH, 15 min, 100 °C, 94%; e) LiAlH_4_, THF, O.N., 0 °C -RT, 83%; f) DMTCl, pyridine, 3.5 h, RT, 95%; g) DMP, pyridine, DCM, O.N., RT, 82%; h) FeCl_3_, MeOH, CHCl_3_, O.N., RT, 90%; i) 1. ammonia (solution in MeOH), MeOH, 0 °C-RT, 5.5 h. 2. H_2_NOSO_3_H, MeOH, O.N., 0 °C-RT. 3. NEt_3_, iodine, MeOH, 8 h, 16%; j) 14, 5 mol% Sc(OTf)_3_, DCM, 2 d, RT, 59%; k) oleic acid, DMAP, EDC×HCl, DCM, O.N., 0 °C-RT, 57%; l) TBAF, acetic acid, THF, O.N., 0 °-RT, 32%; m) 1. NaBPh_4_, water. 2. 2-cyanoethyl-N,N,N’,N’-tetraisopropylphosphoro-amidite, 1H-tetrazole, MeCN, O.N., RT; n) 1. 18, 1H-tetrazole, MeCN, 3.5 h, RT. 2. ^t^BuOOH, MeCN, 1.5 h, 0 °C-RT. 3. NEt_3_, DCM, 20 h, RT, 47% over 3 steps.

The key step of the synthesis was the regioselective epoxide opening reaction used to attach the bifunctional long-chain alcohol at the *sn1* position of the glycerol scaffold. Similar transformations have been reported previously[39–41], but the utilized reagents gave unsatisfactory yields in our hands, presumably due to the longer chain length and the presence of the diazirine and alkyne groups.

The installation of the two functional groups at the ether-linked *sn1* chain required the synthesis of a long-chain alcohol 13 with an internal ketone and a terminal alkyne. Test reactions indicated that the yields of a Grignard reaction with ε-caprolactone[42] as well as a Rh-catalyzed C-C-coupling reaction[43] were not sufficient (see SI for details). Thus, we first synthesized a long-chain fatty acid 8 bearing the internal ketone and a terminal alkyne. To generate the alcohol 13, we initially attempted to selectively reduce the carboxylic acid in the presence of the ketone. The use of mild reducing reagents, such as NBu_4_BH_4_, after activation of the carboxylic group via an acyl chloride only gave moderate yields (see SI for details). The best overall yield was achieved with a complete reduction of carboxylate and ketone to the respective hydroxy groups, followed by a selective protection of the primary hydroxy group with DMTCl. After oxidation of the secondary hydroxy group with Dess-Martin-Periodinane and deprotection via FeCl3, the long-chain alcohol 12 was isolated in a yield of 58% over four steps. We note, that the alternative route via ketone acetalization, reduction of the carboxylate and deprotection was also feasible but gave a lower overall yield. After introducing the diazirine group, the resulting bifunctional alcohol 13 was coupled to a glycerol backbone via an epoxide opening reaction to afford intermediate 15. Many different reagents, both bases and Lewis-acid catalysts, have been described in the literature to afford high yields in similar transformations.[40,41,44–52] In our hands, the epoxide opening worked best with catalytic amounts of Sc(OTf)_3_ (5 mol%) in dry DCM,[52] whereas most published protocols gave either very low yields or no product at all (see SI for details). The bifunctional 1-alkyl-glycerol ether 15 was isolated in 59% yield under optimized reaction conditions (3.5 equivalents of epoxide 14 added, longer reaction time and addition of a second batch of catalyst after 24 h). After subsequent esterification with oleic acid using EDC/DMAP, the silyl protection group at the *sn3* hydroxy group was removed using TBAF. We note that the addition of acetic acid was necessary to minimize acyl chain migration from the *sn2* to the *sn3*-position. The synthesis was concluded by attaching the phosphocholine headgroup to intermediate 17 using phosphoramidite 18.[34] After tetrazole-mediated coupling, the phosphor atom was oxidized with ^t^BuOOH and the cyanoethyl protecting group was removed with NEt_3_ to obtain the bifunctional alkyl ether PC 1 in an overall yield of 2.9%.

### Ether lipid transport depends on the linkage type and position of the bifunctional chain

With the set of bifunctional ether lipids in-hand, we first asked how the linkage type at the *sn1* position influences the intracellular transport behaviour of PCs. Bifunctional ether lipid probes were incorporated into the outer leaflet of the plasma membrane of living cells by incubation with probe-containing liposomes and α-methylcyclodextrin.[36] After the 4 min loading pulse, the loading mix was replaced with cell growth medium and the cells were placed at 37 °C to enable retrograde lipid transport. UV-crosslinking (10 s at 310 nm) after the respective chase times was followed by cell fixation and detergent treatment, achieving both cell permeabilization and the removal of non-crosslinked lipid probes. The generated covalent lipid-protein conjugates were subsequently fluorescently labelled via Cu-mediated click chemistry using an AF594 picolyl-azide dye and lipid localization was assessed by fluorescence microscopy (Fig. 2a).

We first confirmed lipid incorporation into the plasma membrane by analysing the colocalization of lipid signal with a plasma membrane marker in HCT-116 cells after the 4 min loading pulse (Fig. 2b). Control experiments without lipid loading and/or without UV irradiation demonstrate that the observed fluorescence signal is specific to the covalent lipid-protein conjugates, as very low background signal was observed. (Fig. 2b lower panels, Fig. 2c, see supporting information for other timepoints). We observed that both bifunctional lipids modified in the *sn1* position (probes **1**+**2**) exhibited a significant lower overall fluorescence intensity in contrast to their *sn2*-functionalized counterparts (probes **3**-**5**) (Fig. 2c). This could be due to different crosslinking probabilities caused by the chain length difference or a higher likelihood of proteins to interact with the *sn2* sidechain.

In order to analyze lipid transport processes, we performed four-color fluorescence microscopy experiments with iterative immunofluorescent co-staining for the plasma membrane, the endoplasmic reticulum, the Golgi and endosomes. We quantified the lipid signal in each organelle using our recently reported image analysis pipeline based on machine learning-assisted image segmentation.[36] Briefly, self-trained Ilastik Models were used to obtain probability masks to assign the fluorescence signal in the lipid channel to individual organelles and quantify lipid distribution in the cell (Fig. 2d, see SI for details of image background correction and image analysis).

**Figure 2.**
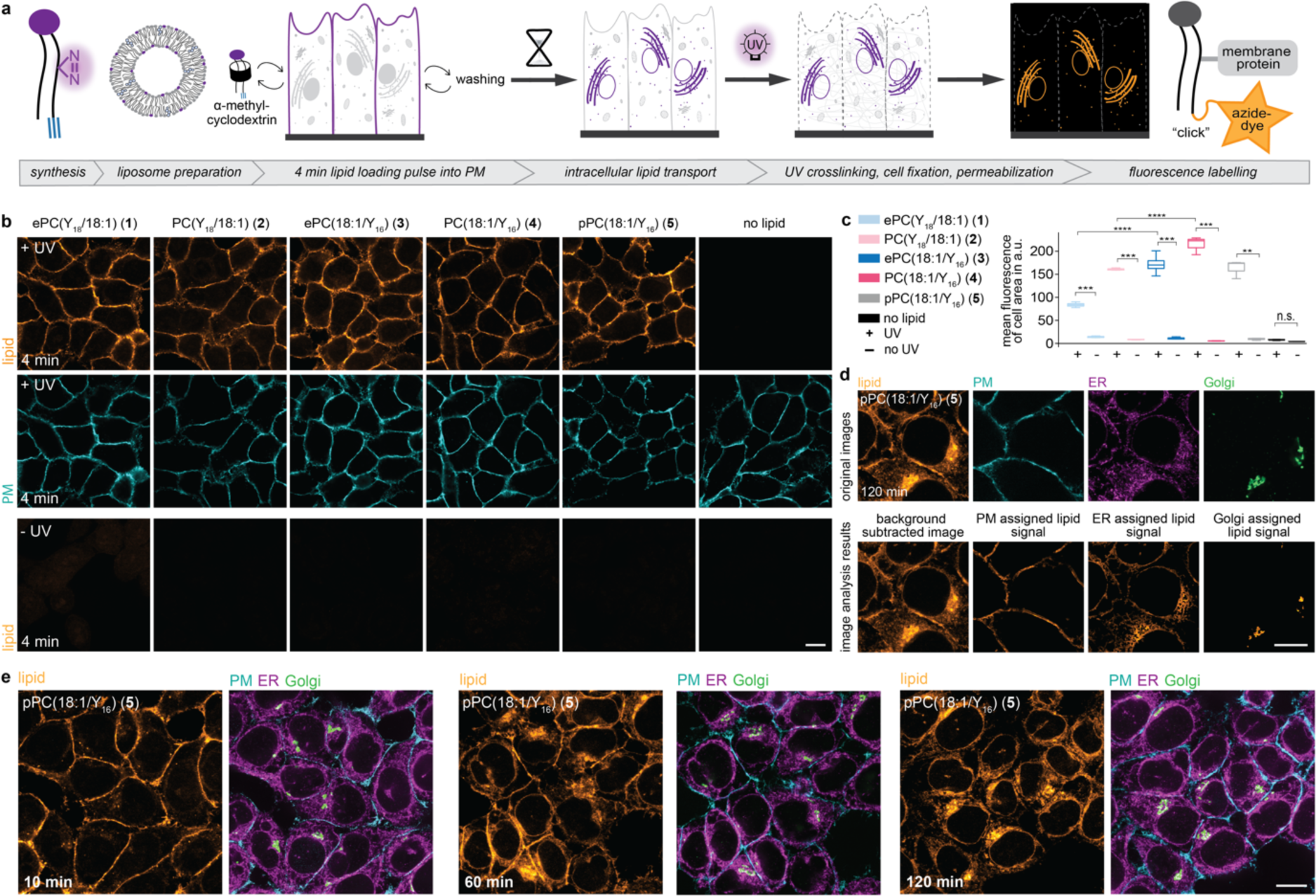
Fluorescence signal derived from bifunctional probes is specific and allows to analyze ether lipid localization after intracellular transport. **a,** General workflow for pulse-chase experiments to monitor ether lipid transport with fluorescence microscopy. **b,** Cellular localization and signal intensity of bifunctional lipids 1-5 in HCT-116 cells after a 4 min pulse with and without UV irradiation. Lipid-protein conjugates were labeled with an Azide-AF594 dye via copper-catalyzed click chemistry. The plasma membrane was labeled with NHS-PEG4-Biotin. The +UV and -UV images for the same probe are shown using identical settings, +-UV image pairs are brightness-contrast adjusted for better comparability as the signal intensity between lipid probes varied. Scale bar is 10 μm. **c,** Quantification of lipid signal intensity at 4 min for all probes and control conditions. The mean fluorescence intensity of the cell-covered area is shown; images were processed using a Gaussian blur (sigma=4) filter and background corrected. The error bars indicate the three-quartile values of the distribution. A permutation test from the Python library Mlxtend (method=approximate, seed=0, number of rounds=1000) was used to test the significance. **d,** Colocalization of lipid signal with plasma membrane, ER and Golgi markers allows to assign the lipid signal to the respective organelles. The plasma membrane was labelled with MemBrite®Fix 405/430. The ER and Golgi were labeled using immunofluorescence. Upper panels: Raw data of all 4 channels. Lower panels: Background corrected lipid signal and lipid signal assigned to the plasma membrane, the ER and the Golgi, respectively. Scale bar is 10 μm. **e,** Cellular localization of pPC(18:1/C_16_) at 10 min, 60 min and 120 min after lipid loading with co-staining of the plasma membrane, ER and Golgi. The plasma membrane was labelled with MemBrite®Fix 405/430. The ER and Golgi were labeled using immunofluorescence. Scale bar is 10 μm.

To quantify changes of lipid distribution over time, we performed pulse-chase experiments with all lipid probes (**1**-**5**) covering a time period of 120 min in HCT-116 cells (Fig. 2e). Lipid signal intensity in the plasma membrane decreased over time for all lipids, indicating probe internalization via retrograde trafficking (Fig. 3a,b, SI Fig.S2-S8 for -UV controls). A comparison between lipid probes revealed that the plasmalogen probe pPC(18:1/Y_16_) (**5**) was retained at the plasma membrane to a larger extent compared to both alkyl ether and ester lipid probes (Fig. 3a,b). Upon internalization most of the signal for all probes was detected within the ER (Fig. 3c). Apart from a brief spike at 20 min, very little fluorescence signal was assigned to endosomal structures (Fig. 3d). Interestingly, all probes bearing the bifunctional fatty acid at the *sn2* position (probes **3**-**5**) exhibited more Golgi localization after 30 min (Fig. 3a,e, SI Fig.S9). This suggested that intracellular lipid transport routes are specific to the presence of the double bond within the *sn1* chain. At 120 min the plasmalogen was more enriched in the Golgi (Fig. 3e) implying a differential steady state distribution between the probes.

To obtain a quantitative description of the lipid transport kinetics and to test the contribution of vesicular and non-vesicular transport routes, we fitted a kinetic model to the obtained time-course data (Fig. 3f,g see SI for model details). The model distinguishes non-vesicular transport from the plasma membrane to the ER from retrograde vesicular transport (plasma membrane via endosomes and the Golgi to the ER). Anterograde lipid transport is described by a summary rate constant from the ER to the plasma membrane capturing both vesicular and non-vesicular transport modes. We found that transport via the non-vesicular route in the retrograde direction (captured by the rate constant k_PM_ER_) was much faster than vesicular membrane trafficking (k_PM_Endo_) for all lipids (Fig. 3h). The alkyl ether and ester lipid probes bearing the bifunctional moieties at the *sn1* chain (probes 1+2) were transported faster than the corresponding *sn2*-modified probes (probes 3-5), which is likely due to both chain length and positioning. However, the linkage type via alkyl ether or ester had only negligible effects on transport kinetics. In contrast, the vinyl ether linkage of the plasmalogen 5 led to overall much slower transport. The steady-state distribution of the plasmalogen probe pPC(18:1/Y_16_) (5) was shifted towards the plasma membrane compared to the corresponding alkyl ether (ePC(18:1/Y_16_) (3) and ester lipid (PC(18:1/Y_16_) (4) probes (quasi-equilibrium constants k_PM_ER_/k_ER_PM_, Fig. 3i). In general, we observed that the effect of plasmalogen-vinyl ether linkage supersedes the effect caused by the placement of the double bond in the *sn1* or *sn2* acyl chain whereas the latter appears to be stronger than the difference between alkyl ether and ester linkage types.

**Figure 3.**
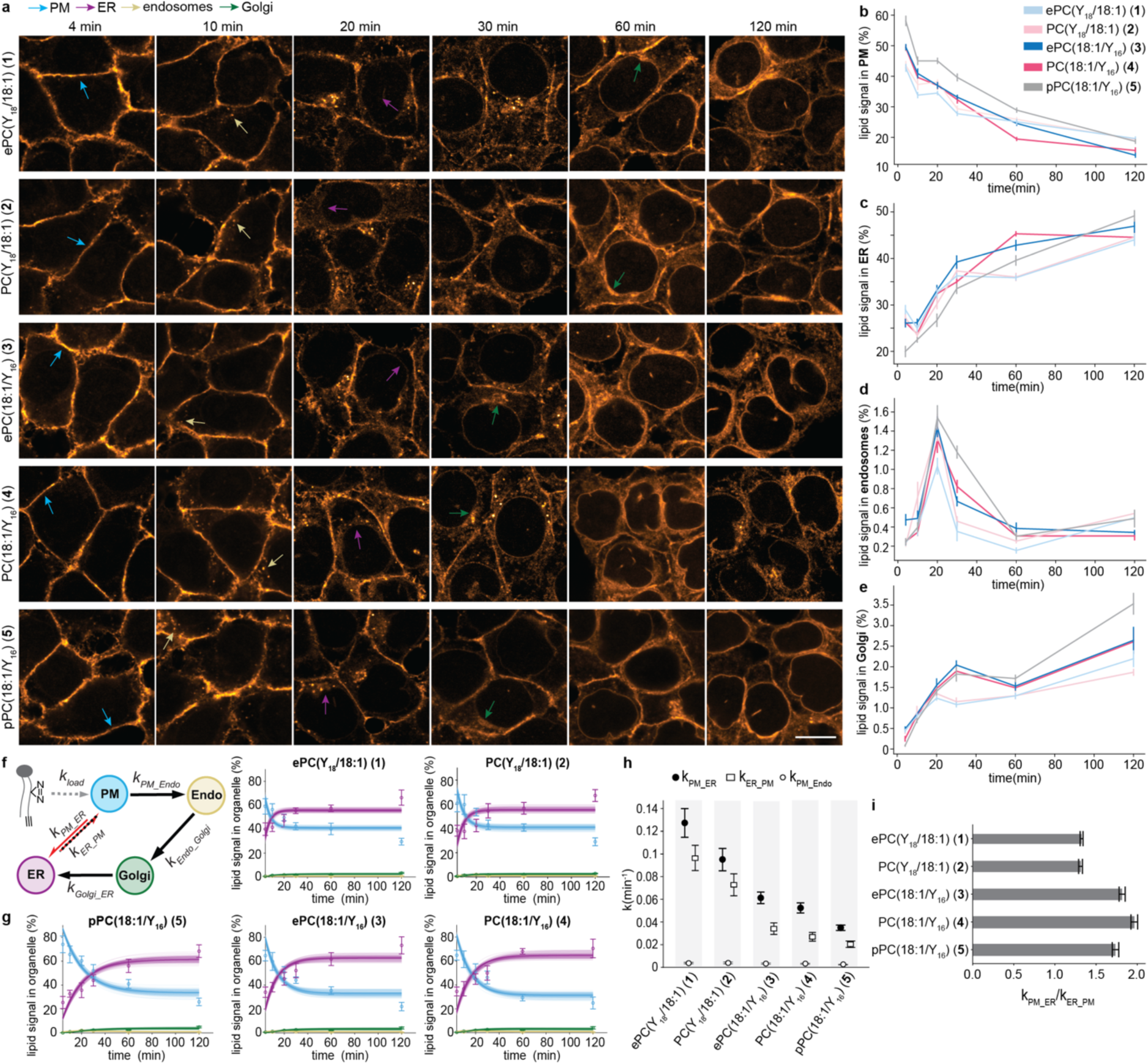
Plasmalogens exhibit slower retrograde transport kinetics than the corresponding alkyl-ether and ester lipid probes in HCT-116 cells. **a,** Cellular localization of bifunctional lipids 1-5 in HCT-116 cells at 4, 10, 20, 30, 60 and 120 min after lipid loading. The lipid was labeled with an Azide-AF594 dye via copper-catalyzed click chemistry. Scale bar is 10 μm. **b-e)**, Quantification of the relative lipid signal in the plasma membrane, the ER, endosomes and the Golgi apparatus. Error bars: SE. **f,** Kinetic model for quantifying lipid transport from fluorescence microscopy data. Black arrows indicate vesicular transport, red arrows non-vesicular transport. **g,** The corresponding model fits of the lipid transport kinetics for all lipids derived from 100 model runs. **h,** Comparison of rate constants describing retrograde vesicular transport from the plasma membrane to endosomes, retrograde non-vesicular transport from the PM to the ER and total transport in the anterograde direction from the ER to the PM. Error bars: SD. **i,** Ratio of the rate constants for non-vesicular lipid transport between the plasma membrane and the ER in the retrograde and anterograde direction. Error bars: SD.

### Ether lipid transport is similar in HCT-116 and U2OS cells

To assess whether the trends observed for ether lipid transport in HCT-116 cells are more generally conserved, we conducted lipid transport experiments in U2OS cells. The two cell lines differ significantly in origin (bone marrow and colon) and morphology. U2OS cells are large, flat cells resembling mesenchymal cells, whereas HCT-116 cells are smaller and retain most structural features of their epithelial origin. We used the *sn2*-bifunctional lipid probes in pulse-chase experiments to assess potential differences in lipid transport. Endpoint quantification at 30 min showed that the plasmalogen was retained at the plasma membrane to a larger extent than both the corresponding ester and alkyl ether probes (Fig. 4 a,b). This effect was also reflected by lower plasmalogen lipid signal in the ER at all timepoints (Fig. 4 a,c). Overall, the relative differences between individual lipids were more pronounced for U2OS cells compared to HCT-116 cells.

One notable difference between the cell lines were bright vesicular structures in the lipid image observed in U2OS but not in HCT-116 cells. These structures were identified as endosomes with immunofluorescence (Fig. 4 d). The quantification of the lipid signal fraction in endosomes confirmed that a larger amount of lipid material was routed through the endocytic route in U2OS cells (Fig. 4 e). This observation suggests that the non-vesicular transport route is even more dominant in HCT-116 cells compared to U2OS cells which is also reflected in the difference between rate constants of PC(18:1/Y16) (**4**) describing endocytosis and non-vesicular transport from the plasma membrane to the ER in the retrograde direction (Fig. 4 f).[36] A closer inspection of lipid incorporation in endosomes in U2OS cells revealed a slower uptake of the plasmalogen probe, which is also reflected in a slower appearance of lipid signal in the Golgi (Fig. 4 g,h and SI Fig. S10). This effect is not caused by a change in the bulk endocytic rate induced by probe loading, as the number and size of endosomes were found to be constant for all probes (SI Fig. S11). Taken together, while some cell line-specific differences were apparent, the major trends for ether lipid transport in both cell lines were similar, in particular with regard to slower internalization kinetics of the plasmalogen probe and predominant non-vesicular transport.

**Figure 4.**
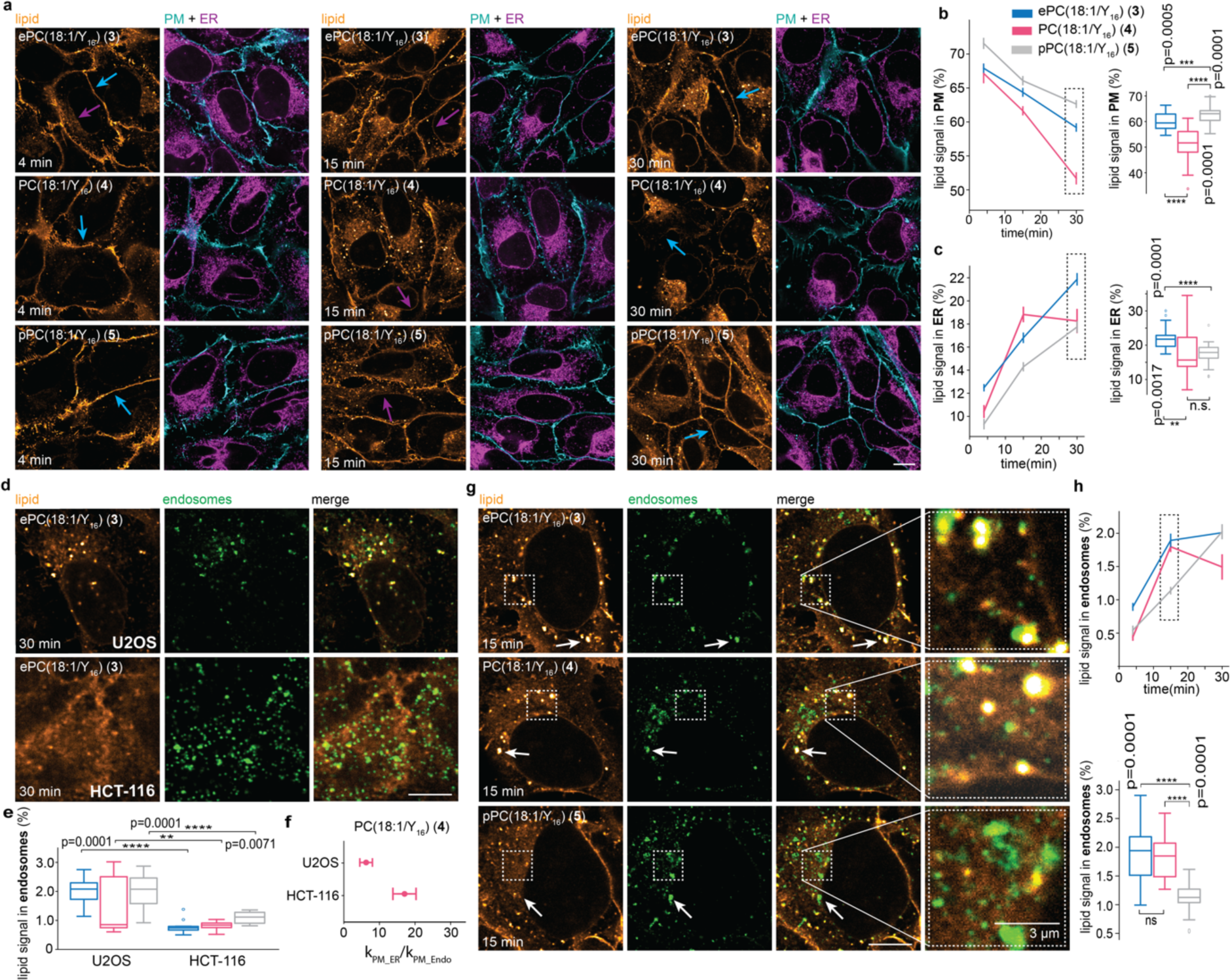
Retrograde vesicular transport of ether lipids is cell-and linkage type specific. **a,** Cellular localization of *sn2*-bifunctional lipid probes at 4 min, 15 min and 30 min in U2OS cells with co-staining of the plasma membrane and the ER. Scale bar is 10 μm. **b-c,** Quantification of the relative lipid intensity in the plasma membrane and the ER. A permutation test from the Python library Mlxtend (method=approximate, seed=0, number of rounds=1000) was used to test the significance at the 30 min timepoint. **d,** Cellular localization of ePC(18:1/Y_16_) in U2OS and HCT-116 cells at the bottom plane after 30 min with co-staining of the Rab5 and Rab7 proteins. Scale bar is 10 μm. **e,** Quantification of the relative lipid signal in endosomes of the *sn2*-bifunctional lipid probes in U2OS and HCT-116 cells after 30 min. **f,** Comparison of ratios between the kinetic rates for non-vesicular (k_PM_ER_) and vesicular (k_PM_Endo_) transport of PC(18:1/Y_16_) in HCT-116 (this work) and U2OS cells[36]. Error bars: SD. **g,** Cellular localization of the sn2-bifunctional lipid probes in U2OS cells at 15 min with co-staining of the Rab5 and Rab7 proteins. Scale bar is 10 μm. **h,** Quantification of the relative lipid signal in endosomes. A permutation test from the Python library Mlxtend (method=approximate, seed=0, number of rounds=1000) was used to test the significance at the 15 min timepoint.

## Discussion

We report a new set of bifunctional photoaffinity probes for studying ether lipid biology. We find notable differences in transport speed for plasmalogen and alkyl ether lipids, suggesting that the lipid transport machinery indeed exhibits specificity for distinct ether lipid types. While very little is known about the cellular roles of individual ether lipid species, our findings imply distinct functions of plasmalogen vs alkyl ether and ester lipids. The effect sizes found here are smaller compared to more structurally diverse lipids with different headgroups and saturation degrees, but nonetheless significant.[36] These data imply that small structural differences are recognized by the cellular lipid handling machinery, providing further evidence for the biological importance of lipid diversity.[53]

The utilized modular synthesis employing the long chain bifunctional alcohol **13**, stereoselective Sc(OTf)_3_ catalyzed epoxide opening, and phosphor amidite headgroup coupling enables straightforward synthetic access to further bifunctional ether lipid classes. This opens the way to study the biology of ether lipids both with regard to detailed analyses of individual lipid species as well as to expand the scope of investigations to the broader ether lipidome. Furthermore, while this study is focused on analyzing ether lipid transport by fluorescence imaging, bifunctional lipids can also be employed for tracing metabolism and identifying lipid-protein interactions.[29,31,35,36] Taken together, the bifunctional ether lipid probes reported here represent a versatile toolkit for studying ether lipid biology in mechanistic detail. This capability will have significant benefits for understanding the functions of ether lipids in fundamental cell biology and their role in human diseases.

## Acknowledgements

AN gratefully acknowledges financial support by the European Research Council (ERC) under the European Union’s Horizon 2020 research and innovation program (grant agreements no GA 758334 ASYMMEM and AURORA). This research was supported by an Allen Distinguished Investigator Award, a Paul G. Allen Frontiers Group advised grant of the Paul G. Allen Family Foundation to AN. AN and KB acknowledge financial support by the VolkswagenStiftung within the Life? initiative. We thank the following services and facilities at MPI-CBG Dresden for their support: Light Microscopy Facility, Computer Department, in particular Oscar Gonzalez for expert advice with regard to high performance computing, and Mass Spectrometry Facility.

