## Supplementary information for "Bifunctional probes reveal the rules of intracellular ether lipid transport"

### Table of Contents

**Supplementary Figures S1-S11** ... **3**

*Fig. S1| Localization of pPC(18:1/$Y_{16}$) after 30 min in HCT-116 and U2OS cells with organelle co-labelling.* 3

*Fig. S2| Negative control without lipid loading and/or without UV-irradiation in HCT-116 cells at 4 min.* ... 4

*Fig. S3| Negative control without lipid loading and/or without UV-irradiation in HCT-116 cells at 10 min.* ... 4

*Fig. S4| Negative control without lipid loading and/or without UV-irradiation in HCT-116 cells at 20 min.* ... 5

*Fig. S5| Negative control without lipid loading and/or without UV-irradiation in HCT-116 cells at 30 min.* ... 5

*Fig. S6| Negative control without lipid loading and/or without UV-irradiation in HCT-116 cells at 60 min.* ... 6

*Fig. S7| Negative control without lipid loading and/or without UV-irradiation in HCT-116 cells at 120 min.* . 6

*Fig. S8| Negative control without lipid loading and/or without UV-irradiation in U2OS cells.* ... 7

*Fig. S9| The plasmalogen is more enriched in the Golgi at 120 min in HCT-116 cells.* ... 7

*Fig. S10| The plasmalogen shows a slower transport to the Golgi in U2OS cells* ... 7

*Fig. S11| The lipid loading has no effect on the overall endosome number and size at the 15 min timepoint* ... 8

**Materials** ... **8**

*Liposome preparation* ... 8

*Lipid loading* ... 8

*Cell fixation, click labelling, immunofluorescence and plasma membrane labelling* ... 8

*Primary antibodies* ... 8

*Secondary antibodies* ... 9

**Methods** ... **9**

*Liposome preparation* ... 9

*Cell culture* ... 9

*Lipid loading* ... 9

*UV-irradiation and cell fixation protocol* ... 10

*Immunostaining protocol* ... 10

*Click labelling of photo-crosslinked lipid-protein conjugates and plasma membrane labeling* ... 11

*Fluorescence microscopy* ... 11

**Image analysis** ... **11**

**Mathematical modelling** ... **13**

**Kinetic models and model performance** ... **13**

**Chemical Synthesis** ... **15**

*General synthetic procedures* ... 15

*Synthesis of ePC($Y_{18}$/18:1) (1)* ... 15

*Synthesis of PC($Y_{18}$/18:1) (2)* ... 21

*Synthesis of sn2-bifunctional lipids* ... 22

*Test reactions for the synthesis of the bifunctional long chain alcohol* ... 24

*Optimization of epoxide opening reaction* ... 25

**NMR spectra of new compounds** ... **26**

### Supplementary Figures S1-S11

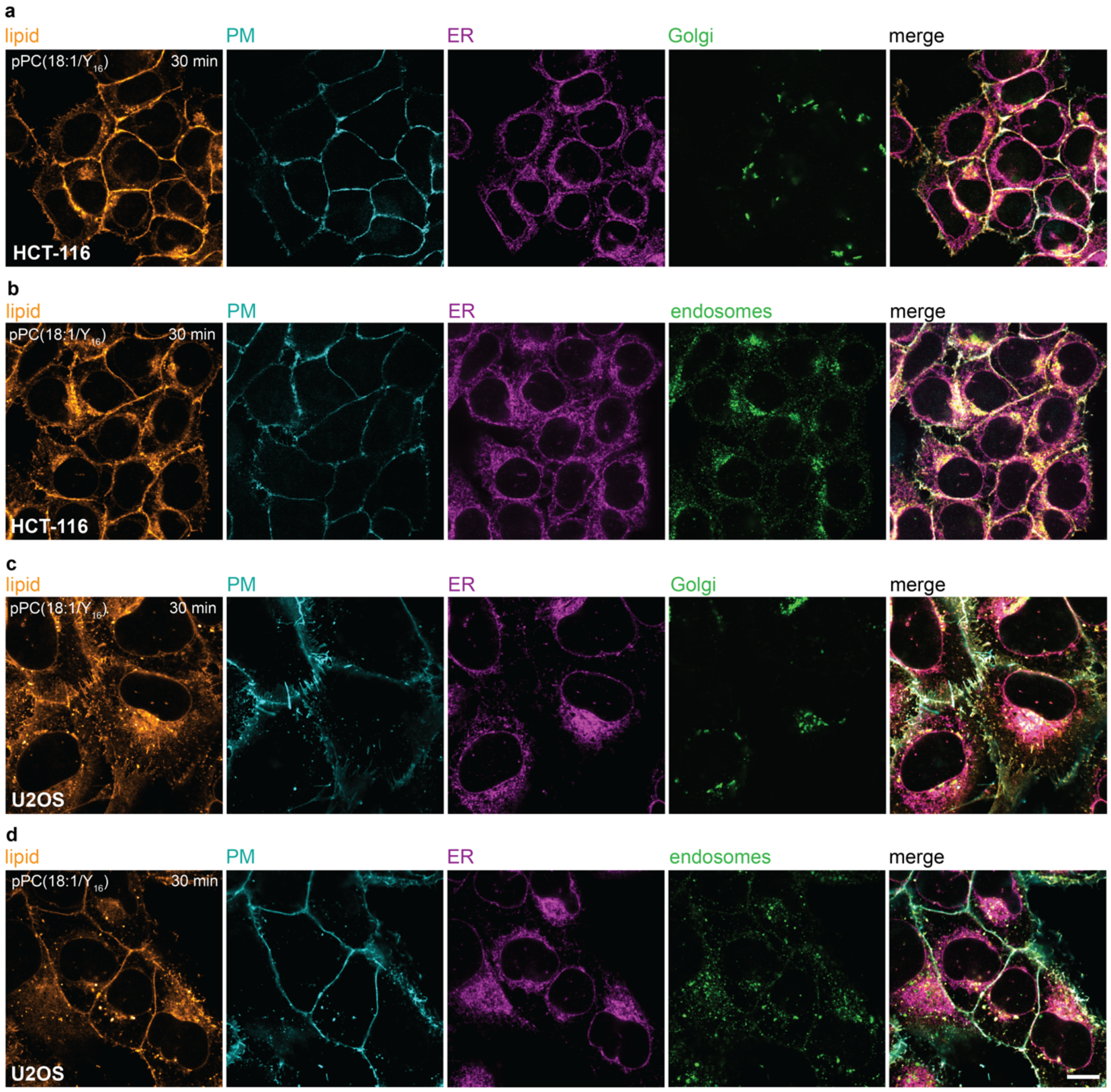


**Fig. S1| Localization of pPC(18:1/$Y_{16}$) after 30 min in HCT-116 and U2OS cells with organelle co-labelling. a,** Cellular localization of pPC(18:1/$Y_{16}$) after 30 min in HCT-116 cells with co-staining of the plasma membrane, ER and the Golgi. **b,** Cellular localization of pPC(18:1/$Y_{16}$) after 30 min in HCT-116 cells with co-staining of the plasma membrane, ER and endosomes. **c,** Cellular localization of pPC(18:1/$Y_{16}$) after 30 min in U2OS cells with co-staining of the plasma membrane, ER and the Golgi. **d,** Cellular localization of pPC(18:1/$Y_{16}$) after 30 min in U2OS cells with co-staining of the plasma membrane, ER and endosomes. Scale bar is 10 µm.

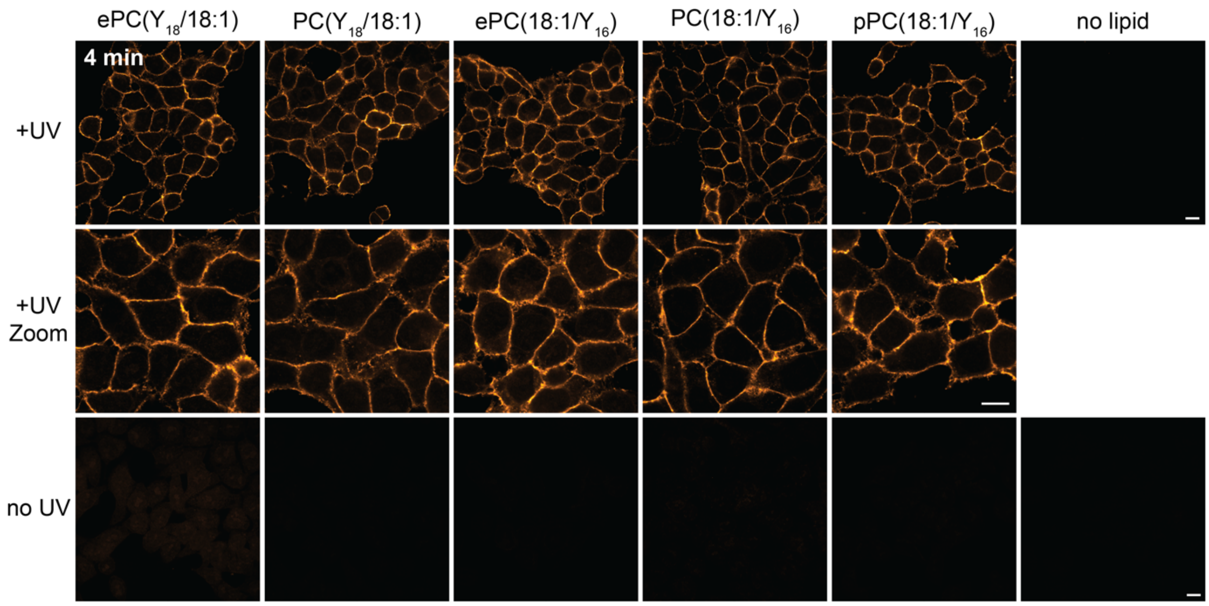


**Fig. S2| Negative control without lipid loading and/or without UV-irradiation in HCT-116 cells at 4 min.** The UV and no UV-images are adjusted to the same intensity individually for each lipid. Scale bar is 10 µm.

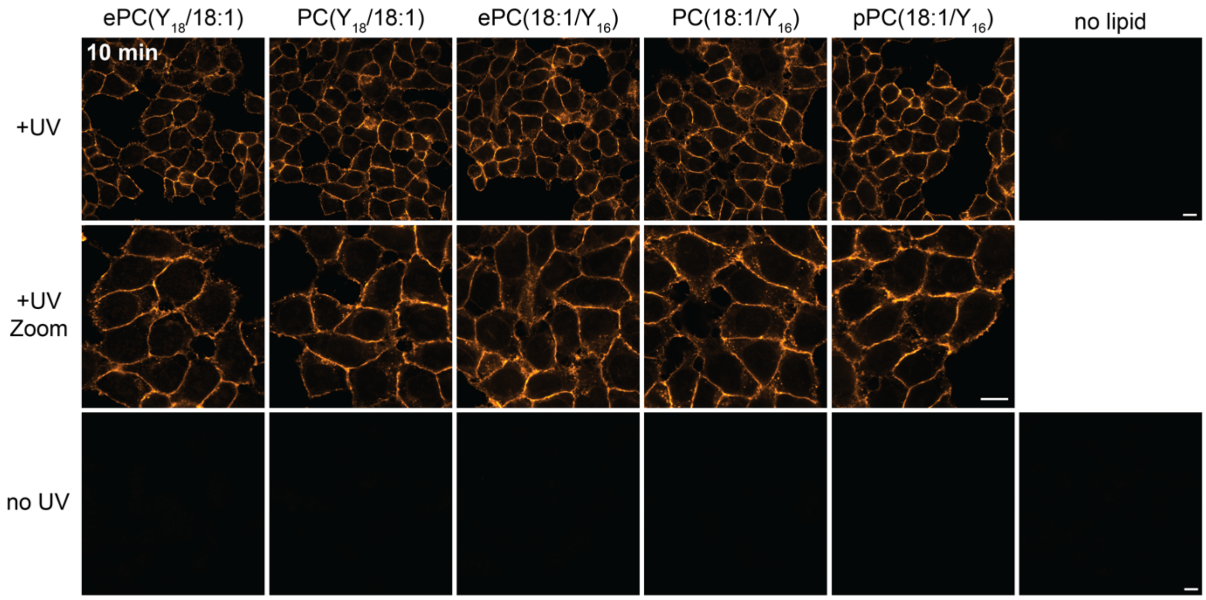


**Fig. S3| Negative control without lipid loading and/or without UV-irradiation in HCT-116 cells at 10 min.** The UV and no UV-images are adjusted to the same intensity individually for each lipid. Scale bar is 10 µm.

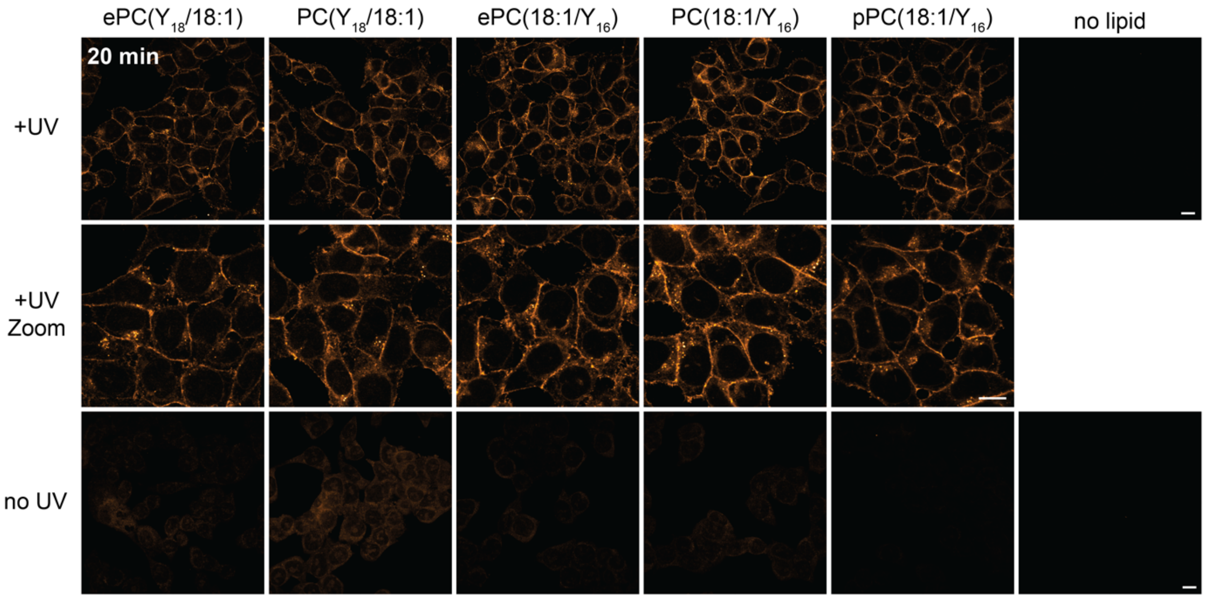


**Fig. S4| Negative control without lipid loading and/or without UV-irradiation in HCT-116 cells at 20 min.** The UV and no UV-images are adjusted to the same intensity individually for each lipid. Scale bar is 10 µm.

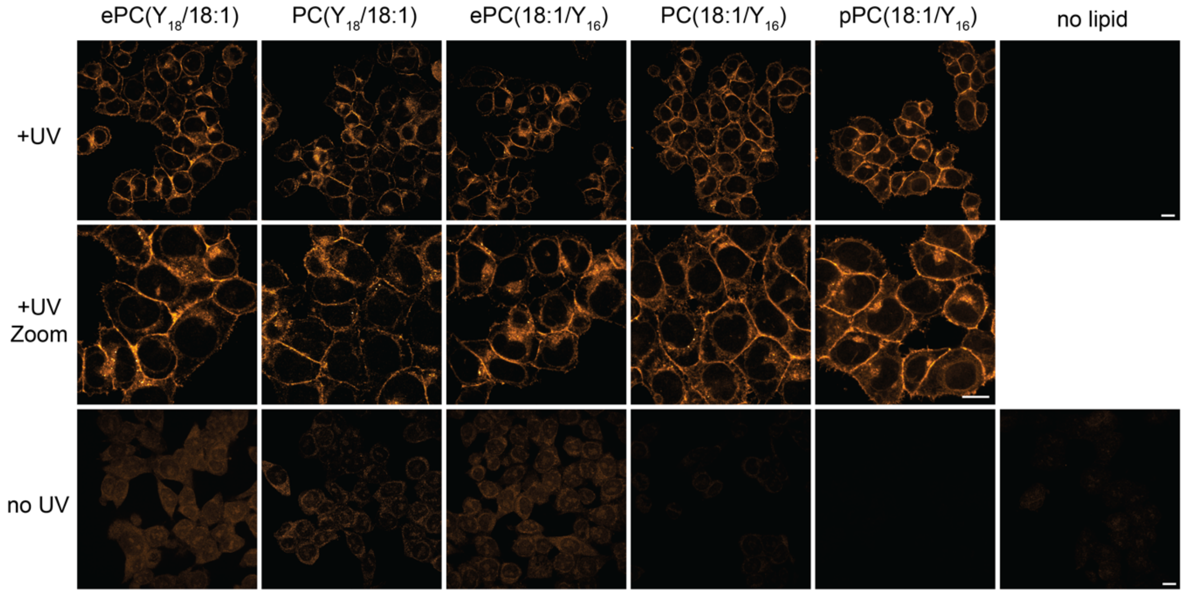


**Fig. S5| Negative control without lipid loading and/or without UV-irradiation in HCT-116 cells at 30 min.** The UV and no UV-images are adjusted to the same intensity individually for each lipid. Scale bar is 10 µm.

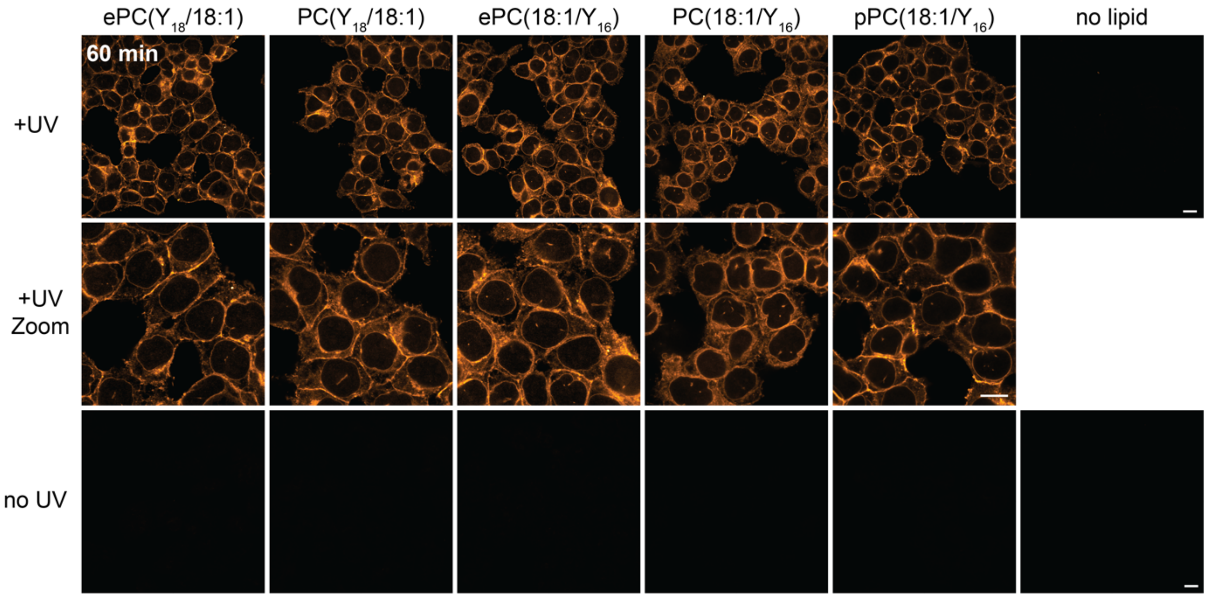


**Fig. S6| Negative control without lipid loading and/or without UV-irradiation in HCT-116 cells at 60 min.** The UV and no UV-images are adjusted to the same intensity individually for each lipid. Scale bar is 10 µm.

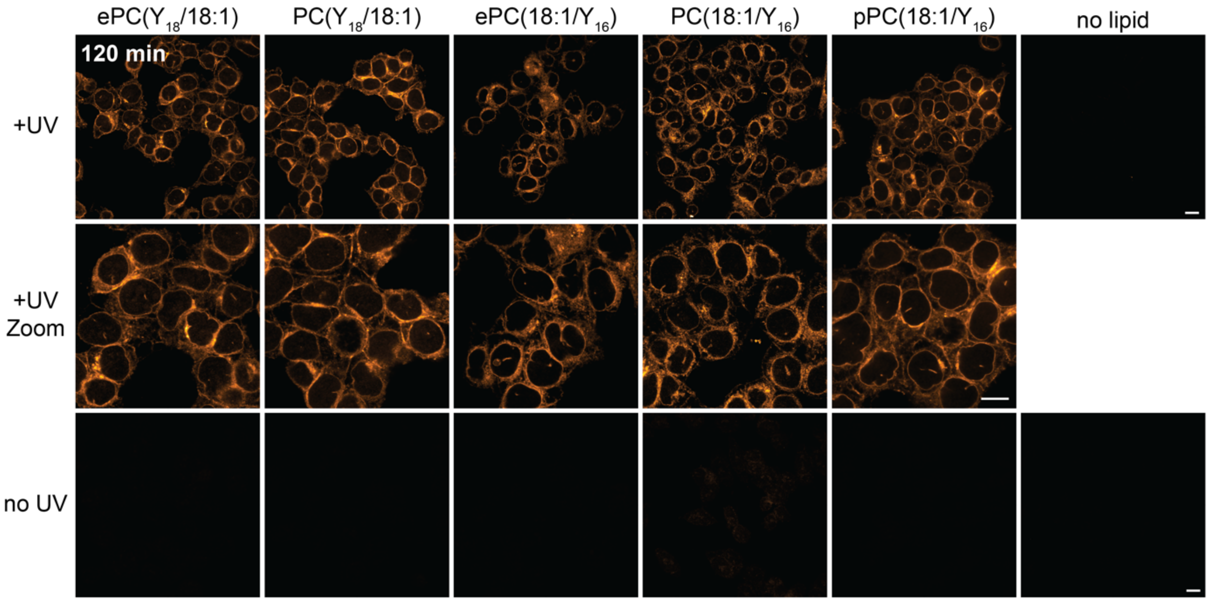


**Fig. S7| Negative control without lipid loading and/or without UV-irradiation in HCT-116 cells at 120 min.** The UV and no UV-images are adjusted to the same intensity individually for each lipid. Scale bar is 10 µm.

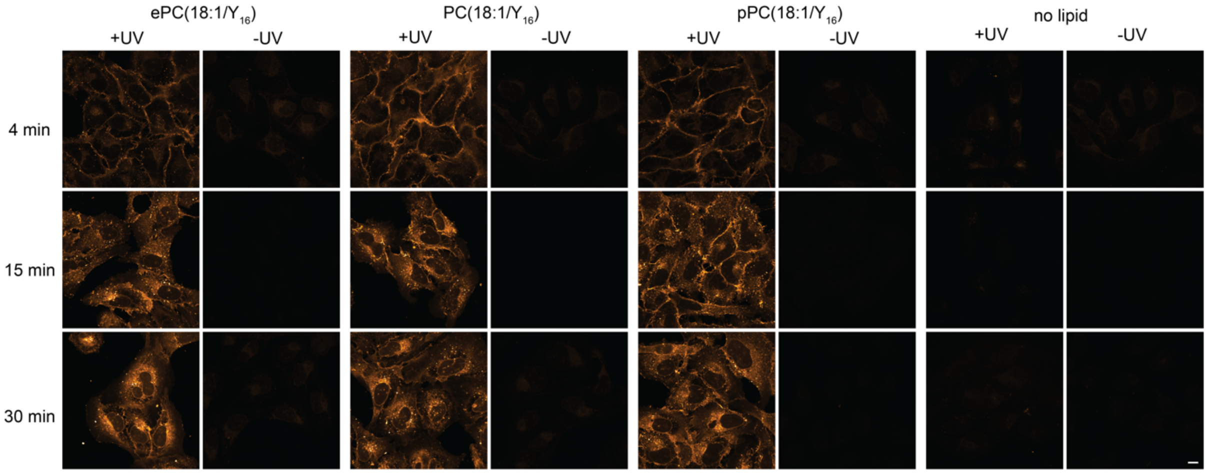


**Fig. S8| Negative control without lipid loading and/or without UV-irradiation in U2OS cells.** All images from one timepoint are adjusted to the same intensity. Scale bar is 10 µm.

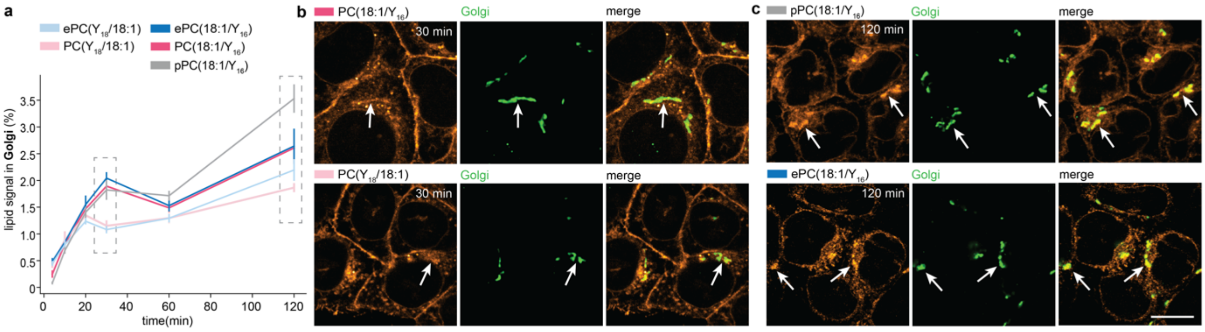


**Fig. S9| The plasmalogen is more enriched in the Golgi at 120 min in HCT-116 cells. a**, Quantification of the relativ lipid signal in the Golgi apparatus in HCT-116 cells. **b**, Cellular Localization of PC(18:1/$Y_{16}$) and PC($Y_{18}$/18:1) at 30min with co-staining of the Golgi. Scale bar is 10 µm. **c**, Cellular localization of pPC(18:1/$Y_{16}$) and ePC(18:1/$Y_{16}$) at 120 min with co-staining of the Golgi. Scale bar is 10 µm.

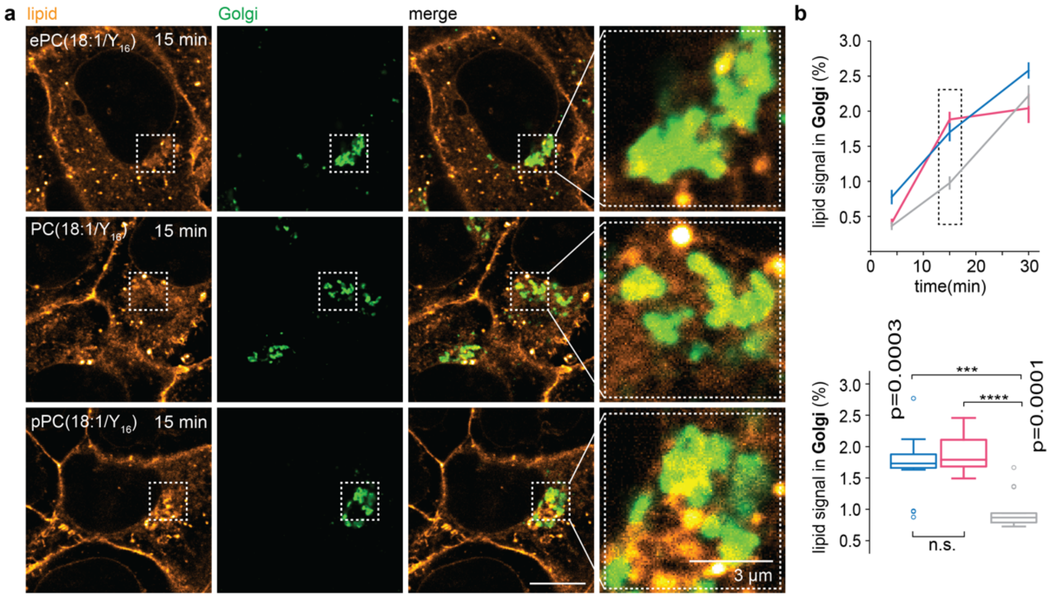


**Fig. S10| The plasmalogen shows a slower transport to the Golgi in U2OS cells. a,** Cellular localization of the *sn2*-bifunctional lipids at 15 min with co-staining of the Golgi. **b**, Quantification of the relative lipid signal in the Golgi. Scale bar is 10 µm.

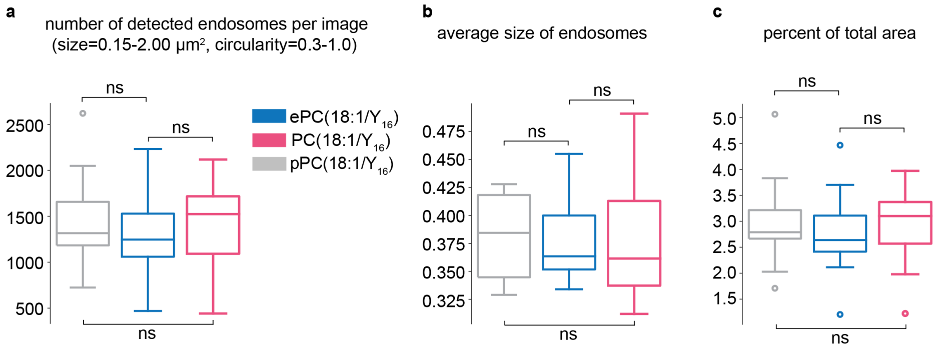


**Fig. S11| The lipid loading has no effect on the overall endosome number and size at the 15 min timepoint. a**, Quantification of the number of endosomes. **b**, Quantification of the average size of endosomes. **c**, Quantification of the percentage of the cell covered area assigned to endosomes. The last frame of the stack (bottom plane) of the endosomal antibody-channel was used for quantification based on a staining with Rab5/rabbit antibody in a dilution of 1:35000 and the Rab7/rabbit-antibody in a dilution of 1:500 to label the endosomes. ImageJ was used to analyze the detected particles. The threshold was set to a minimum of 400. All particles with a size of 0.15-2.00 $\mu m^2$ and a circularity of 0.3-1.00 were analyzed. The data was plotted with Python using the Seaborn library. The data includes two biological replicates with 40 images per lipid.

### Materials

**Liposome preparation**

1-palmitoyl-2-oleoyl-glycero-3-phosphocholine (POPC) (MW: 760.08) was purchased from Avanti. Cholesterol (MW: 386.65) was purchased from Sigma. Chlorofom was purchased from TCI (P0018). LiposoFast Basic was purchased from Avestin. A Nuclepore™ Track-Etch Membrane from Whatsman® (diameter 47 mm, pore size 0.1 µm, polycarbonate) was used for the extrusion to prepare liposomes.

**Lipid loading**

U2-OS and HCT-116 cells were purchased from ATCC. Cellview microplates, 96 well (Item No. 655891) were purchased from Greiner. Methyl-alpha-cyclodextrin (Av. MW: 1127) was purchased from Cyclodextrin-Shop. McCoy's 5A medium (16600082), FBS (A5670701) and Penicillin-Streptomycin (15140122) were purchased from Thermo Fisher Scientific. The FBS was heat inactivated by heating up the serum at 56ºC for 30 min. Hank's Balance Salt Solution (HBSS) was purchased as a salt from Merck (H6136) and prepared according to the manufacturer's instructions.

**Cell fixation, click labelling, immunofluorescence and plasma membrane labelling**

Formaldehyde solution was purchased from SAV Liquid Production GmbH (FN-1000-4-1). Triton X-100 (Av. MW: 625) was purchased from Serva. AF594-Picolyl-Azide (MW: 925.00) was purchased from Jena Bioscience. Glycine (MW: 75.07, 50046)), CuSO4 (MW: 159.61, 45167), L-ascorbic acid (MW: 176.12, A92902) were purchased from Merck. MAXpack Immunostaining Media Kit (including Blocking, Staining and Washing Medium, 15252-15254) was purchase from Active Motif. MemBrite® Fix 405/430 Cell Surface Staining Kit was purchased from biotium (30092-T). EZ-Link NHS-PEG4-Biotin (MW: 588.67, A39259) and Streptavidin, Alexa Fluor 405 conjugate (Av. MW: 57 056, S32351) were purchased from Thermo Fisher Scientific. The NHS-PEG4-Biotin was resuspended using acetonitrile and aliquoted, aliquots were then dried under a vacuum centrifuge and stored at -20 ºC.

**Primary antibodies**

Antibody mixes containing several antibodies were used to homogenously label organelles. Two different antibody mixes were used. Golgi/ER and Endosomes/ER antibody mixes were prepared in MAXbind Staining Solution at the given dilution.

| Organelle | Antibody | Dilution | Supplier | Code |
|---|---|---|---|---|
| **Golgi** | Golgin97 (D8P2K) (rabbit) | 1:3500 | Cell Signaling Technologies | CST-13192S, Lot: 1 |
| | Giantin (rabbit) | 1:3500 | Abcam | ab80864,<br>Lot: GR3209923-1 |
| | GM130 XP (D6B1) (rabbit) | 1:3500 | Cell Signaling Technologies | CST-12480S,<br>Lot: 3 |
| **ER** | Calnexin 1 (mouse) | 1:1000 | Abcam | ab112995,<br>Lot: GR3246794-6 |
| | Lamin A/C (4C11) (mouse) | 1:1000 | Cell Signaling Technologies | CST-4777S, Lot:5 |
| | BAP31 (mouse) | 1:1000 | Enzo Life Sciences | ALX-804-601-C100,<br>Lot: L15093 |
| | Calreticulin (mouse) | 1:1000 | Abcam | ab22683,<br>Lot: GR3361946-5 |
| **Endosomes** | Rab5 (C8B1) (rabbit) | 1:35000 | Cell Signaling Technologies | CST-3547S<br>Lot: 7 |
| | Rab7 (D95F2) XP® (rabbit) | 1:500 | Cell Signaling Technologies | CST-9367S<br>Lot:3 |

**Table S1**: List of used primary antibodies

**Secondary antibodies**

| Antibody | Dilution | Supplier | Code |
|---|---|---|---|
| Goat anti-mouse lgG AFPlus 647 | 1:1000 | Thermo Fisher Scientific | A32728<br>Lot: YB363606 |
| Goat anti-Rabbit lgG AF488 | 1:1000 | Thermo Fisher Scientific | A11008<br>Lot: A11008 |

**Table S2**: List of used secondary antibodies

### Methods

The protocol for visualizing lipid transport was slightly adapted from Iglesias et. al and description of the protocol details is reproduced here with the necessary small alterations for the convenience of the reader.[1]

**Liposome preparation**

Liposomes were prepared by mixing chloroform solutions of POPC, bifunctional lipid and cholesterol in a 1:2:1 ratio to a final lipid concentration of 3 mM (0.75 mM POPC, 1.5 mM bifunctional lipid, 0.75 mM cholesterol). The chloroform was then removed under argon stream to form a lipid film that was subsequently rehydrated with filtered PBS to the required volume. The liposome solutions were vortexed and sonicated repeatedly until no clumps remained. This homogenous emulsion was extruded 21 times with a 100 nm filter and stored at 4 °C.

**Cell culture**

U2OS and HCT-116 cells were cultured using McCoys's 5A+GlutaMAX$^{TM}$ medium supplemented with 10% fetal bovine serum (FBS) and 100 U/ml of Gibco Penicillin-Streptomycin. Cells were cultured in 75 $cm^2$ flasks and splitted every 2-3 days. Cells were kept at 37 °C with 5% $CO_2$ and used up to passage 15. Cells were seeded in 96 well plates with a density of 10.000 cells per well for U2OS cells and 40.000 cells per well for HCT-116. The cells were seeded 1 day prior to lipid transport experiments. For cell counting, a Neubauer counting chamber was used and the cells were mixed with a 0.4% trypan blue stain in a ratio of 1:1.

**Lipid loading**

Bifunctional lipids were loaded onto cells using 0.5 mM lipids in liposomes in the presence of 4 mM α-methylcyclodextrin in HBSS. The mixture was incubated at 37 °C for at least 30 min. Before loading the liposomes, the cells were washed twice with 200 µl of pre-warmed HBSS. The addition of 75 µl of the liposome-cyclodextrin solution marks the starting point for the different periods of time that were examined.

Plasma membrane labelling of U2OS and HCT-116 cells with NHS-PEG4 Biotin and Streptavidin, Alexa Fluor 405 conjugate (*This method was used for the time-course experiments in U2OS cells and to test specificity of fluorescence signal derived from bifunctional lipid probes in HCT-116 cells (Fig. 2b and Fig. S2-S8))*

For the 4 min time point NHS-PEG4-Biotin was added to the liposome/cyclodextrin solution to a final concentration of 5 mM immediately before loading.
For all other timepoints, the cells were incubated for 4 min at 37 °C with the liposome-cyclodextrine solution to load the lipids into the plasma membrane of the cells. After washing with HBSS the cells were left with complete growing medium (McCoys's 5A+GlutaMAX™ medium supplemented with 10% fetal bovine serum (FBS) and 100 U/ml of Gibco Penicillin-Streptomycin). A 5 mM NHS-PEG4-Biotin solution in HBSS was added to the cells 4 minutes before UV irradiation and fixation to obtain a plasma membrane stain. Before UV-crosslinking the cells were washed twice with 200 µl pre-warmed HBSS.

Plasma membrane labelling of HCT-116 cells with MemBrite® Fix 405/430 *(This method was used for the dataset of the time-course experiments to quantify the relative lipid signal in organelles in HCT-116 cells. We used this alternative method, as the above-described protocol repeatedly failed to yield a bright, homogenous plasma membrane stain in HCT-116 cells. In particular the basolateral areas (bottom planes) were found to be less accessible for staining with* NHS-PEG4-Bioti.

For the 4 min time point, cells were first treated with MemBrite® Fix 405/430 to label the plasma membrane before adding the liposome/cyclodextrin solution. In detail, cells were washed twice with 200 µl of pre-warmed HBSS. The cells were incubated with the Pre-Staining Solution (1X in HBSS) for 3 min at 37 °C. Cells were washed once with 100 µl HBSS and incubated with 50 µl of the MemBrite® Fix stain solution (1X in HBSS) for 3 min at 37 °C. After one wash with HBSS, the cells were incubated for 4 min at 37 °C with the liposome-cyclodextrine solution to load the lipids into the plasma membrane of the cells. Before UV-crosslinking the cells were washed twice with 200 µl pre-warmed HBSS.

For all other timepoints, the cells were incubated for 4 min at 37 °C with the liposome-cyclodextrine solution to load the lipids into the plasma membrane of the cells. After washing with HBSS the cells were left with complete growing medium (McCoys's 5A+GlutaMAX™ medium supplemented with 10% fetal bovine serum (FBS) and 100 U/ml of Gibco Penicillin-Streptomycin). Around 6 minutes before UV irradiation, the cells were washed once with HBSS and incubated with 50 µl of the MemBrite® Fix 405/430 Pre-Staining Solution (1X in HBSS) for 3 min at 37 °C. Cells were washed once with 100 µl HBSS and incubated with 50 µl of the MemBrite® Fix stain (1x) in HBSS for 3 min at 37 °C. Before UV-crosslinking the cells were washed twice with 200 µl pre-warmed HBSS.

**UV-irradiation and cell fixation protocol**

After a certain period of time, cells were exposed to UV-light (310 nm) for 10 seconds. A row of 8 LEDs was used to illuminate the wells from beneath the 96-well plate through the glass bottom plate. The LEDs were supplied by Violumas (high power 2x2COB 310 nm LED without lens, #VC2X2C48LX-310), had an average maximum power output of 290 mW and the cells were placed at an approximately distance of 0.5 mm from the light source. Immediately after UV-irradiation, the medium was removed and the cells were fixed with a solution of 4 % paraformaldehyde in PBS for 20 min at RT. Subsequently, the cells were washed with 0.1% Triton in 100 mM glycine solution in PBS and incubated for 1 h at room temperature with the Triton-solution. After washing three times with PBS, the plates were kept at 4 °C.

**Immunostaining protocol**

For organelle staining a MAXpack™ Immunostaining kit was used. PBS was removed and the cells were incubated with 80 µl of MAXblock Blocking Medium for 1 hour at 37 °C. The Blocking solution was removed and the cells were washed with 100 µl of MAXwash washing medium three times with 10 minutes of incubation time each time. The washing medium was removed and 80 µl of primary antibodies (see **Table S1**) diluted in MAXbind were added to the correct wells. The cells were incubated with the antibodies for 1 hour at 37 °C. The cells were washed three times with MAXwash washing medium (10 min incubation) and the secondary antibodies (see **Table S2)** diluted in MAXstain medium were added. The cells were incubated for 1 hour at room temperature in dark environment. The antibody solution was removed and the cells were washed three times with MAXwash (10 min incubation time). The plates were kept with PBS at 4 °C.

**Click labelling of photo-crosslinked lipid-protein conjugates and plasma membrane labeling**
After washing thrice with 100 mM HEPES, 75 µl of the Click-solution (97mM PBS, 0.002 mM AF568-picolyl azide-dye, 5 mM ascorbic acid, 0.5 mM THPTA, 0.1 mM $CuSO_4$) were added to each well and the fixed cells were incubated for 40 min at 37 °C. Samples were washed thrice with PBS and thrice with 100 mM HEPES before 75 µl of freshly prepared Click solution was added. The fixed cells were incubated for additional 20 min. Samples were washed thrice with PBS and kept at 4 °C.

*For samples that were incubated with NHS-PEG4-Biotin during lipid loading:* Before imaging, cells were incubated with 50 µl of 0.1 µM Streptavidin Alexa Fluor 405 conjugate for at least 1 h at room temperature to label the plasma membrane.

**Fluorescence microscopy**
Images were acquired using an Olympus 100x silicone immersion oil objective (UPLSAPO100XS) mounted in an Olympus IX83 microscope. The microscope was fitted with a Yokogawa CSU-W1SoRa unit and images were captured by an ORCA-Fusion camera from Hamamatsu. Four laser lines were used for excitation: 405 nm, 488 nm, 564 nm and 641 nm with their respective filter sets. All images were acquired with identical settings. The plasma membrane was imaged in the 405 channel and the lipid in the 561 channel. The 488 and 647 channels were used for two other organelles. HCT-116 cells were imaged using 25-slice z-stacks and U-2OS cells were imaged using 14-slice z-stacks, both with 0.5 µm distance in z between frames. To ensure that all images were acquired at the same starting plane, the Olympus TruFocus Z-drift compensation system was used. Each frame was exposed for 250 ms with 75% laser power.

### Image analysis

The image analysis pipeline was slightly adapted from Iglesias et. al.[1]

Briefly, fluorescence microscopy z-stack images were analyzed in a High-Performance Computing cluster using Python[2] and Ilastik[3]. The Python code and the Ilastik models used can be found in http://doi.org/10.5281/zenodo.12784327. HCT-116 image stacks consisted of a 25-frame stack and U2OS images stacks of a 14-frame stack, both with 0.5 µm distance between the individual frames. Original images can be downloaded from https://doi.org/10.6019/S-BIAD1293.

Images were selected by visual inspection before automated image analysis. Images that contained large, highly fluorescent clumps indicative of apoptotic and dead cells were excluded from the analysis. Control images were samples not loaded with bifunctional lipids but otherwise treated the same to assess nonspecific background signal. Both +UV and -UV conditions were recorded. Control images were acquired on every 96-well plate.

The processing of lipid images, determination of cell covered area and allocation of lipid signal to individual organelles were identical to the routines in Iglesias et al..[1] The small adaptations in the Ilastik Models and backgorund correction are described below.

#### Ilastik Models
For segmenting the PM, ER, Golgi and endosomes, the Autocontext workflow in Ilastik was used that runs Pixel Classification twice and shows the result of the first stage to the second stage of the process. All pixel features and scales were used. Images were segmented in a High-Performance Computing cluster.

#### Background correction
Background correction for U2OS cells was done according to Iglesias et.al..[1] For HCT-116 cells an additional step was required, as the HCT cells often showed a high intracellular background fluorescence (also in the nucleus) which was observed in some cases for the NoUV/+lipid controls as well. This resulted in too high values for the quantified relative lipid intensity in the ER. The fluorescence in the nuclei was defined as background, and the nuclei were segmented by blurring the ER Mask (obtained from immunofluorescence labelling) with a gaussian filter (sigma=3). The obtained image was inverted and the individual regions were labelled. The cell free area was excluded by only measuring regions with an area from 1000 to 15000 pixels. The mean intensity in the defined regions was subtracted from the image in addition to the background subtraction based on the control images without lipid loading.

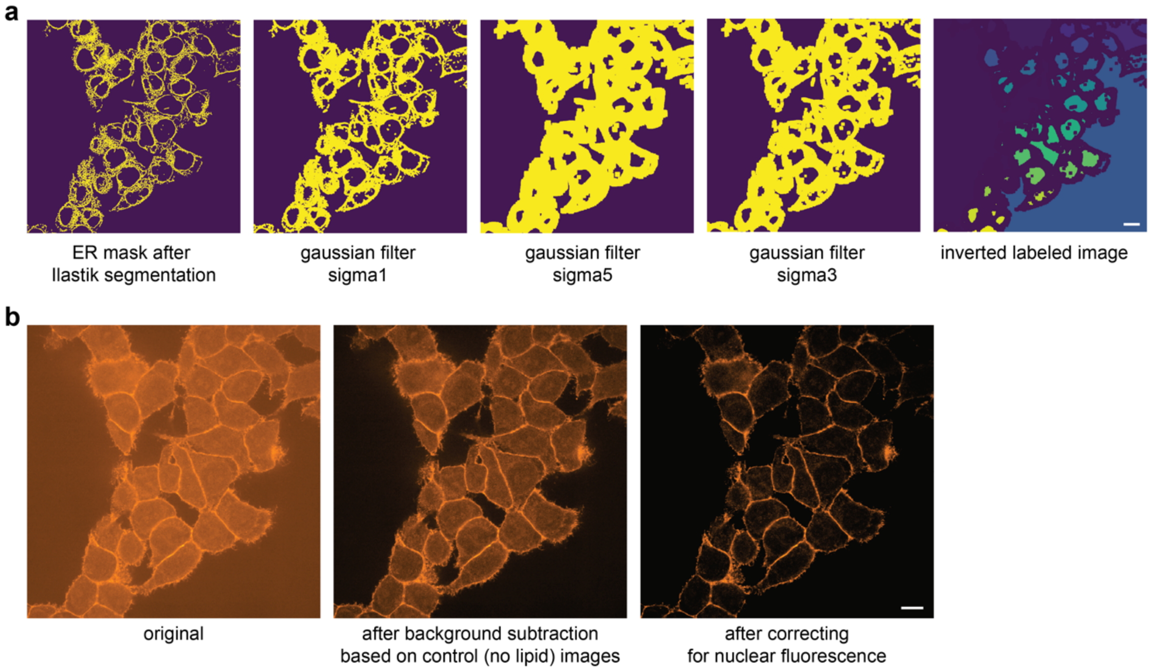


**Fig. S12| Background subtraction for HCT-116 cells. a**, The sigma value of a gaussian filter was optimized to segment nuclei regions based on the ER mask obtained after Ilastik segmentation. Only regions with an area between 1000 and 15000 pixels were selected to exclude the cell free area. **b**, The original image was first background subtracted based on the intensity of the no lipid/+UV images. Afterwards, the mean intensity of the nuclei-regions was subtracted from the image. Scale bar is 10 µm.

#### Calculation of mean intensity of the cell area at 4 min in HCT cells for Figure 2

The overall mean fluorescence was corrected by the fluorescence of the cell free area. For the calculations one frame (frame 20/25) was used and a gaussian filter (from SciPy library, sigma=4) was applied. Three different regions of the cell free area were selected manually and the mean fluorescence intensities were measured with ImageJ. The sum of the mean and four times the standard deviation was subtracted from the blurred image. All negative and zero values were removed and the mean fluorescence intensity was calculated with Python.

### Mathematical modelling

The kinetic model was slightly adapted from Iglesias et. al and description of the modelling process is reproduced here with the necessary small alterations for the convenience of the reader.[1]

The entire modeling process can be reproduced using the Matlabscripts provided separately in http://doi.org/10.5281/zenodo.12784327 ('Optimization Toolbox' is needed for lsqcurvefit).
At t0 = 0 the total quantity of each lipid resides in the extracellular space ("external reservoir"), representing the situation where the loading solution has just been added (Et0 = 100 %) and the plasma membrane (Mt0 = 0 %) is free of lipid. M can be populated by lipids from E with a specific rate ($k_E > 0$ ). Furthermore, lipid from M can be transported to an acceptor membrane ($A(t_0) = 0$ %) with a specific rate ($k_A > 0$). Both processes can be combined to a set of ordinary differential equation (ODE) together with the initial conditions (initial value problem).

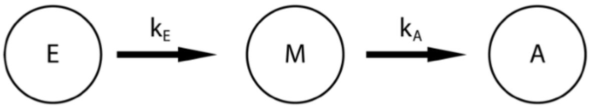


$$\frac{dE}{dt} = -k_E \cdot E(t)$$

$$\frac{dM}{dt} = k_E \cdot E(t) - k_A \cdot M(t)$$

$$\frac{dA}{dt} = k_A \cdot M(t)$$

The differential equations for the individual organelles were constructed successively (see Kinetic Models section and file:"DGLModel_MSNorm.m") To extract lipid transport rate constants, the ODE system - which was solved numerically using ode45 solver in Matlab R2020b - was embedded in a parameter optimization routine based on Matlabs nonlinear least-square solver (lsqcurvefit).

After deriving the transport rates, the model was calculated (solid non-transparent lines in the model plots). The robustness of the model was then tested using a wild Bootstrap / Monte Carlo with normally distributed weights.[4,5] For the resampling, synthetic noise was generated in the same order of magnitude as the residuals of the fit. This was added to the model and the transport rate optimization was restarted (100 MC runs). The new models are shown with transparency (1% opacity) in the corresponding plots. This allowed us to check the reproducibility of the fit and to estimate the variance of the rates (standard deviation of the 100 MC runs).
As the model is a superposition of different transport processes, it is difficult to compare individual sub-processes directly with each other. To estimate the efficiency of vesicular and non-vesicular transport pathways from the PM to the endosomes (Endo) and to the ER, a model for these compartments was calculated that only includes the optimized transport rates (from the 100 MC runs) from the plasma membrane to Endosomes ($k_{PM\_Endo}$) and the ER ($k_{\mathrm{PM\_ER}}$), respectively.

### Kinetic models and model performance

The model is a "transport only" model (see Fig. 3f) that describes the exchange of lipids between cellular membranes and does not account for depletion of bifunctional lipids over time. The fraction of lipid signal (total assigned intensity normalized to 100%) in each organelle derived from imaging data is used as input and the sum does not change over time. The used model is the "minimal" model for the purpose of this study which contains the required minimum number of parameters required to fit the data.
The model captures vesicular retrograde transport along the secretory pathway from the PM via Endosomes and the Golgi to the ER and retrograde non-vesicular lipid transport from the PM to the ER. Anterograde lipid transport is captured by a summary rate $k_{\mathrm{ER\_PM}}\mathrm{ER}(t)$ which encompasses all lipid transport (vesicular and non-vesicular).

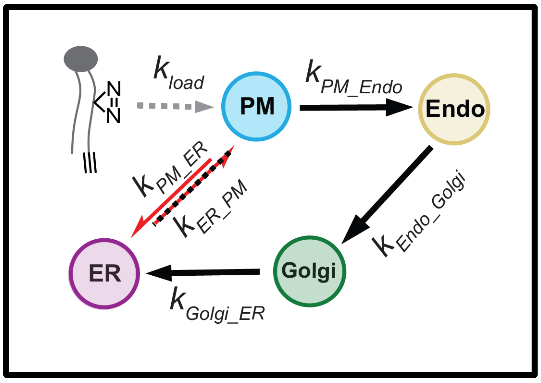


$$\frac{dLipo}{dt} = -k_{Lipo\_PM}Lipo(t)$$

$$\frac{dPM}{dt} = -k_{Lipo\_PM}Lipo(t) + k_{ER\_PM}ER(t) - k_{PM\_Endo}PM(t) - k_{PM\_ER}PM(t)$$

$$\frac{dEndo}{dt} = k_{PM\_Endo}PM(t) - k_{Endo\_Golgi}ER(t)$$

$$\frac{dER}{dt} = k_{PM\_ER}PM(t) + k_{Golgi\_ER}Golgi(t) - k_{ER\_PM}ER(t)$$

$$\frac{dGolgi}{dt} = k_{Endo\_Golgi}Endo(t) - k_{Golgi\_ER}Golgi(t)$$

The rate constant for lipid transfer from the liposomes to the plasma membrane is not accessible for two reasons. First, there are no data points for the incorporation period (<4 minutes) and second, the exact fraction of total liposomal lipid transferred to the cells is unclear due to the sample preparation. Since all of the lipid later detected was rapidly transferred from the liposomes to the plasma membrane during the incorporation process, the same fast rate constant was fixed (1000) for all lipids.

**Supplementary Table 1: Rate constants**

| ***Rates ($min^{-1}$)*** *Mean ± STD* | *ePC($Y_{18}$/18:1)* | *PC($Y_{18}$/18:1)* | *ePC(18:1/$Y_{16}$)* | *PC(18:1/$Y_{16}$)* | *pPC(18:1/$Y_{16}$)* |
|---|---|---|---|---|---|
| **PM→Endo** | 0.0039±0,0007 | 0,0039±0,0008 | 0,0033±0,0006 | 0,0032±0,0005 | 0,0024±0,0003 |
| **PM→ER** | 0.1273±0,0127 | 0,0950±0,0098 | 0,0612±0,0051 | 0,0524±0,0044 | 0,0349±0,0024 |
| **ER→PM** | 0.0964±0,0111 | 0,0728±0,0097 | 0,0341±0,0051 | 0,0269±0,0038 | 0,0203±0,0028 |
| **Endo→Golgi** | 0.2165±0,0485 | 0,1570±0,0390 | 0,1099±0,0220 | 0,1076±0,0253 | 0,0809±0,0146 |
| **Golgi→ER** | 0.0646±0,0159 | 0,0674±0,0156 | 0,0337±0,0089 | 0,0331±0,0085 | 0,0219±0,0052 |

### Chemical Synthesis

**General synthetic procedures**
All chemicals were obtained from commercial sources (Acros, Sigma-Aldrich, TCI chemicals, Avanti Polar Lipids, Alfa Aesar, Roth, Fluka or Merck) and were used without further purification. Solvents for flash chromatography were obtained from VWR and dry solvents from Sigma. Deuterated solvents were obtained from Deutero GmbH, Karlsruhe, Germany. TLC was performed on precoated plates of silica-gel (Merck, 60 F254) using UV-light (254 nm or 365 nm) or staining solution of phosphomolybdic acid in EtOH (3 g phosphomolybdic acid in 100 ml EtOH) for analysis. Preparative column chromatography was performed using silica gel from Merck (silica 60, grain size 0.063-0.200 nm) with a pressure of 1 bar. $^{1}$H-, $^{13}$C- and $^{31}$P-NMR spectra were measured on 400 MHz Advance$^{TM}$ III HD Nanobay Bruker spectrometer. Chemical shift of $^{1}$H- and $^{13}$C-NMR spectra are referenced indirectly to tetramethylsilane. J values are given in Hz and chemical shifts in ppm. Splitting patterns are mentioned as follows: s, singlet; d, doublet; t, triplet; q, quartet; m, multiplet; m$_c$, centered multiplet. $^{13}$C-NMR spectra were broadband hydrogen decoupled. Mass sepctra (ESI) were acquired using a QExactive instrument (Thermo Fisher Scientific) equipped 660 with a robotic nanoflow ion source.

**Synthesis of ePC(Y$_{18}$/18:1) (1)**

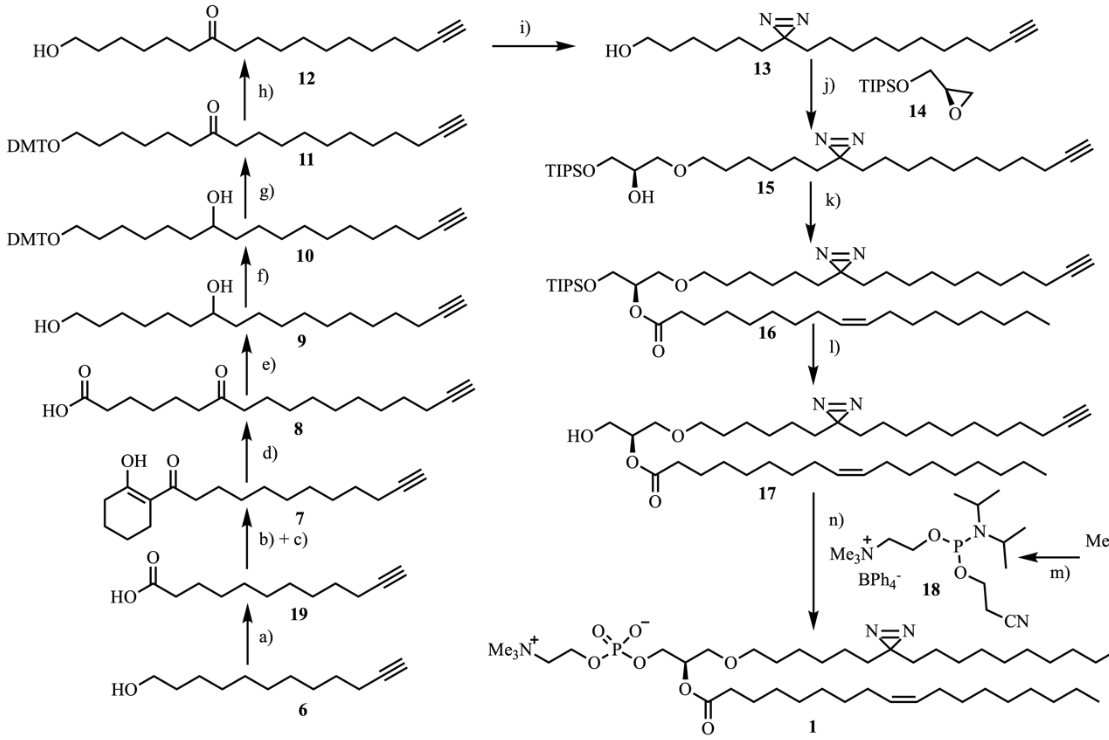


**Scheme A: Overview of synthesis of ePC(Y$_{18}$/18:1) (1). a)** $CrO_3$, $H_2SO_4$, $H_2O$, 0 °C-RT, 1h, 85%; **b)** $(COCl)_2$, DMF, DCM, 1 h, 0 °C-RT; **c)** 1-morpholino-cyclohexene, $NEt_3$, DCM, O.N., RT, 76% over 2 steps; **d)** aq. KOH, 15 min, 100 °C, 94%; **e)** $LiAlH_4$, THF, O.N., 0 °C-RT, 83%; **f)** DMTCl, pyridine, 3.5 h, RT, 95%; **g)** DMP, pyridine, DCM, O.N., RT, 82%; **h)** $FeCl_3$, MeOH, $CHCl_3$, O.N., RT, 90%; **i)** 1. ammonia (solution in MeOH), MeOH, 0 °C-RT, 5.5 h, 2. $H_2NOSO_3H$, MeOH, O.N., 0 °C-RT, 3. $NEt_3$, iodine, MeOH, 8 h, 16%; **j) 14**, 5 mol% $Sc(OTf)_3$, DCM, 2 d, RT, 59%; **k)** oleic acid, DMAP, EDC·HCl, DCM, O.N., 0 °C-RT, 57%; **l)** TBAF, acetic acid, THF, O.N., 0 °-RT, 32%; **m)** 1. $NaBPh_4$, water. 2. 2-Cyano-ethyl-N,N,N',N'-tetraisopropylphosphorodi-amidite, 1*H*-tetrazole, MeCN, O.N., RT;[6] **n)** 1. **18**, 1*H*-tetrazole, MeCN, 3.5 h, RT. 2. $^{t}$BuOOH, MeCN, 1.5 h, 0 °C-RT. 3. $NEt_3$, DCM, 20 h, RT, 47% over 3 steps.

Compounds **14** and **18** were synthesized according to literature protocols. The analytical data was in accordance with those reported.[6,7] All other protocols and analytical data are listed below.

#### *Dodec-11-ynoic acid (19)*

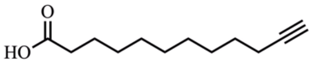


Freshly prepared Jones reagent (3.4 g $CrO_3$ (34 mmol, 1.2 eq.), 10 ml $H_2O$, 7.5 ml $H_2SO_4$) was added dropwise to a solution of 11-dodecyn-1-ol (5.0 g, 27 mmol, 1.0 eq.) in 500 ml acetone at 0 °C. The reaction mixture was stirred for 30 min at 0°C and subsequently 30 min at room temperature. Acetone was removed under reduced pressure and the remaining aqueous solution transferred onto a mixture of EtOAc and $H_2O$. The aqueous layer was extracted with EtOAc (2x). The combined organic layers were dried over $Na_2SO_4$ and the solvent removed under reduced pressure. The obtained light yellowish oil was purified by flash chromatography (CyHex/EtOAc 7:1, 2% acetic acid). Dodec-11-ynoic acid (**19**) was isolated as a white solid.

**$^1$H NMR (400 MHz, $CD_3OD$)** δ = 2.28 (t, *J* = 7.4 Hz, 2H), 2.20 – 2.11 (m, 3H), 1.59 ($m_c$, 2H), 1.55 – 1.46 (m, 2H), 1.46 – 1.24 (m, 10H) ppm.
**$^{13}$C NMR{$^1$H} (101 MHz, $CD_3OD$)** δ = 177.67, 85.05, 69.35, 34.93, 30.46, 30.35, 30.19, 30.13, 29.74, 29.69, 26.06, 19.01 ppm.
**HR-MS (ESI negative)** m/z calculated for $C_{12}H_{20}O_2$: 196.146; found: 195.138 $[M-H]^-$.
**Yield**: 4.6 g (23 mmol, 85 %).

#### *1-(2-Hydroxycyclohex-1-en-1-yl)dodec-11-yn-1-one (7)*

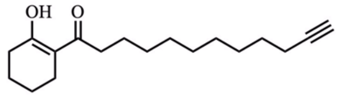


A solution of dodec-11-ynoic-acid (**19**) (4.4 g, 22 mmol, 1.0 eq.) in 25 ml dry DCM was treated with 7 drops of dry DMF. Subsequently, oxalyl chloride (2.2 ml, 33 mmol, 1.4 eq.) was added dropwise at 0 °C. The reaction mixture was stirred for 20 min at 0 °C followed by 40 min at room temperature. A solution of 1-morpholinocyclohexene (7.5 g, 44.8 mmol, 2.0 eq.) in 30 ml dry DCM was heated to 35 °C and then the acyl chloride mixture was added dropwise. Subsequently, the reaction mixture was stirred at room temperature overnight. Aqueous HCl (55 ml, 20 % v/v) and 95 ml chloroform were added and the mixture was stirred at room temperature for 8 h. The phases were separated and the organic phase was washed with water (3x). The combined aqueous layers were reextracted with DCM (3x) and the organic phases were then combined and their solvent removed under reduced pressure. The crude product was purified by flash chromatography (*n*-hexane/EtOAc, 95:5). **Compound 7** was isolated as a light yellowish oil.

**$^1$H NMR (400 MHz, $CD_3OD$)** δ = 2.44 (t, *J* = 7.4 Hz, 2H), 2.39 – 2.26 (m, 4H), 2.21 – 2.11 (m, 3H), 1.75 – 1.64 (m, 4H), 1.59 ($m_c$, 2H), 1.54 – 1.46 (m, 2H), 1.46 – 1.24 (m, 10H) ppm.
**$^{13}$C NMR{$^1$H} (101 MHz, $CD_3OD$)** δ = 203.45, 181.87, 107.89, 85.06, 69.38, 37.99, 31.82, 30.53, 30.51, 30.42, 30.17, 29.77, 29.71, 25.39, 24.86, 23.96, 22.75, 19.03 ppm.
**HR-MS (ESI positive)** m/z calculated for $C_{18}H_{28}O_2$: 276.209; found: 277.220 $[M+H]^+$.
**Yield:** 4.7 g (17.0 mmol, 76 %).

#### *7-Oxooctadec-17-ynoic acid (8)*

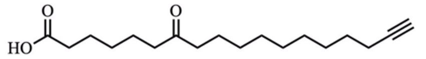


Compound **7** (7.0 g, 25.3 mmol, 1.0 eq.) was heated to 100 °C and treated with aq. KOH (50 ml, 60 % v/v). After stirring for 15 min at 100 °C the reaction mixture was cooled to room temperature. The reaction mixture was diluted with 150 ml water and slowly neutralised by dropwise addition of conc. aqueous HCl (20 ml). Upon neutralization, a colourless precipitate of crude product appeared which was collected by filtration. Further crude product was obtained by extracting the remaining water phase with chloroform (3x), combining the organic layers and removing the solvent under reduced pressure. The crude product was purified by flash chromatography (DCM/MeOH, 97:3). 7-Oxooctadec-17-ynoic acid (**8**) was isolated as a white solid.

**$^1$H NMR (400 MHz, $CD_3OD$)** δ = 2.46 (t, *J* = 7.0 Hz, 2H), 2.45 (t, *J* = 7.0 Hz, 2H), 2.28 (t, *J* = 7.4 Hz, 2H), 2.20 – 2.11 (m, 3H), 1.67 – 1.45 (m, 8H), 1.44 – 1.36 (m, 2H), 1.36 – 1.21 (m, 10H) ppm.*
*signals (triplets) at 2.46 and 2.45 ppm overlap.

**$^{13}$C NMR{$^{1}$H} (101 MHz, $CD_3OD$)** δ = 214.13, 177.52, 85.07, 69.36, 43.48, 43.22, 34.75, 30.47, 30.27, 30.14, 29.74, 29.70, 25.84, 24.88, 24.50, 18.99 ppm.*
*signals between 29.6 and 30.6 ppm overlap.
**HR-MS (ESI negative)** m/z calculated for $C_{18}H_{30}O_3$: 294.219; found: 293.215 $[M-H]^-$.
**Yield:** 7.0 g (23.8 mmol, 94 %).

#### *Octadec-17-yne-1,7-diol (9)*

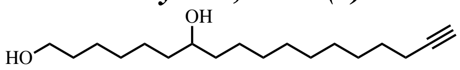


$LiAlH_4$ (640 mg, 17.1 mmol, 2.5 eq.) was suspended in 120 ml dry THF under an argon atmosphere. 2.0 g of 7-oxooctadec-17-ynoic acid (**8**) (6.8 mmol, 1.0 eq.) were dissolved in 40 ml dry THF and added slowly at 0 °C. The reaction mixture was stirred over night at room temperature. The reaction was quenched with a sat. aqueous $NH_4Cl$-solution and all solids were removed by filtration. The reaction mixture was transferred onto a mixture of EtOAc and $H_2O$, the layers were separated and the aqueous layer extracted with EtOAc (2x). The combined organic layers were dried over $Na_2SO_4$ and the solvent removed under reduced pressure. The crude product was purified by flash chromatography (CyHex/EtOAc, 1:1). Octadec-17-yne-1,7-diol (**9**) was isolated as a colourless oil.

**$^{1}$H NMR (400 MHz, $CD_3OD$)** δ = 3.54 (t, $J$ = 6.6 Hz, 3H), 3.51 ($m_c$, 1H), 2.21 – 2.08 (m, 3H), 1.63 – 1.22 (m, 26H) ppm.
Triplet at 3.54 and centered multiplet at 3.51 overlap.
**$^{13}$C NMR{$^{1}$H} (101 MHz, $CD_3OD$)** δ = 85.08, 72.43, 69.33, 62.99, 38.44, 38.38, 33.63, 30.83, 30.68, 30.61, 30.21, 29.78, 29.72, 26.96, 26.80, 19.01 ppm.
**HR-MS (ESI positive)** m/z calculated for $C_{18}H_{34}O_2$: 282.256; found: 300.286 $[M+NH_4]^+$.
**Yield:** 1.6 g (5.7 mmol, 83 %)

#### *1-(Bis(4-methoxyphenyl)(phenyl)methoxy)octadec-17-yn-7-ol (10)*

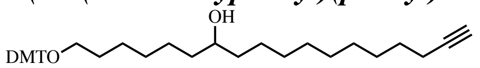


A solution of octadec-17-yne-1,7-diol (**9**) (2.2 g, 7.8 mmol, 1.0 eq.) in 90 ml dry pyridine was treated with DMTCl (2.64 g, 7.80 mmol, 1 eq.). The reaction mixture was stirred at room temperature for 3.5 h. Pyridine was removed under reduced pressure and the crude product was purified by flash chromatography (CyHex/EtOAc, 9:1, 0.2 % $NEt_3$). **Compound 10** was isolated as a colourless oil.

**$^{1}$H NMR (400 MHz, $CD_3OD$)** δ = 7.45 – 7.36 (m, 2H), 7.33 – 7.15 (m, 7H), 6.87 – 6.79 (m, 4H), 3.76 (s, 6H), 3.54 – 3.45 (m, 1H), 3.04 (t, $J$ = 6.5 Hz, 2H), 2.19 – 2.08 (m, 3H), 1.65 – 1.53 (m, 2H), 1.53 – 1.19 (m, 24H) ppm.*
**$^{13}$C NMR{$^{1}$H} (101 MHz, $CD_3OD$)** δ = 159.95, 146.93, 137.89, 131.18, 129.30, 128.63, 127.59, 113.93, 87.02, 85.09, 72.38, 69.36, 64.32, 55.68, 38.39, 38.29, 31.01, 30.83, 30.70, 30.61, 30.53, 30.21, 29.79, 29.72, 27.42, 26.77, 26.69, 19.02 ppm.*
*The spectra show low level of an impurity of DMTOH (grey arrows) which was removed in subsequent steps (see NMR spectra of **11**). The yield was calculated using the ratio of the integrals from the aromatic protons of the product and the impurity as they showed a slightly different chemical shift.
**HR-MS (ESI negative)** m/z calculated for $C_{39}H_{52}O_4$: 584.387; found: 629.383 $[M+HCOO]^-$.
**Yield:** 4.3 g, (7.4 mmol, 95 %).

#### *Synthesis of 1-(Bis(4-methoxyphenyl)(phenyl)methoxy)octadec-17-yn-7-one (11)*

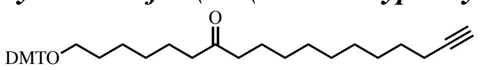


A solution of of **compound 10** (7.3 g, 12.5 mmol, 1.0 eq.) in 80 ml DCM was added dropwise to a solution of Dess-Martin-periodinane (10.6 g, 25.0 mmol, 2.0 eq.) in 60 ml DCM and 60 ml pyridine. The reaction mixture was stirred at room temperature overnight. After removing the solvents under reduced pressure, the crude product was purified by flash chromatography (CyHex/EtOAc, 9:1, 0.2 % $NEt_3$). **Compound 11** was isolated as a colourless oil.

**$^{1}$H NMR (400 MHz, $CD_3OD$)** δ = 7.41 (d, *J* = 7.6 Hz, 2H), 7.34 – 7.22 (m, *J* = 15.5, 8.2 Hz, 6H), 7.17 (t, *J* = 7.2 Hz, 1H), 6.82 (d, *J* = 8.7 Hz, 4H), 3.76 (s, 6H), 3.03 (t, *J* = 6.3 Hz, 2H), 2.48 – 2.31 (m, 4H), 2.21 – 2.03 (m, 3H), 1.62 – 1.43 (m, *J* = 13.6, 6.6 Hz, 8H), 1.43 – 1.08 (m, 16H) ppm.
**$^{13}$C NMR{$^{1}$H} (101 MHz, $CD_3OD$)** δ = 214.25, 159.96, 146.92, 137.87, 131.19, 129.30, 128.64, 127.61, 113.95, 87.04, 85.09, 69.38, 64.22, 55.69, 43.50, 43.38, 30.88, 30.47, 30.26, 30.14, 30.00, 29.75, 29.70, 27.20, 24.90, 24.80, 19.01 ppm.
**HR-MS (ESI negative)** m/z calculated for $C_{39}H_{50}O_4$: 582.371; found: 627.368 $[M+HCOO]^-$.
**Yield:** 6.0 g (10 mmol, 82 %).

#### *1-Hydroxyoctadec-17-yn-7-one (12)*

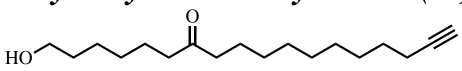


**Compound 11** (3.0 g, 5.2 mmol, 1.0 eq.) was placed in a 500 ml flask. A solution of anhydrous $FeCl_3$ (1.1 g, 6.8 mmol, 1.3 eq.) in 200 ml MeOH and 200 ml $CHCl_3$ was added. The reaction mixture was stirred over night at room temperature. The solvents were removed under reduced pressure and the crude product was purified by flash chromatography with a gradient (CyHex/EtOAc, 8:2 to 6:4). 1-Hydroxyoctadec-17-yn-7-one (**12**) was isolated as a colorless solid.

**$^{1}$H NMR (400 MHz, $CD_3OD$)** δ = 3.54 (t, *J* = 6.6 Hz, 2H), 2.46 (t, *J* = 7.3 Hz, 2H), 2.45 (t, *J* = 7.3 Hz, 2H), 2.23 – 2.09 (m, 3H), 1.63 – 1.45 (m, 8H), 1.45 – 1.18 (m, 14H) ppm.*
* Signals (triplets) at 2.46 and 2.45 ppm overlap.
**$^{13}$C NMR{$^{1}$H} (101 MHz, $CD_3OD$)** δ = 214.08, 85.05, 69.39, 62.86, 43.47, 43.39, 33.46, 30.47, 30.27, 30.14, 30.10, 29.74, 29.69, 26.73, 24.88, 24.84, 19.02 ppm.
**HR-MS (ESI positive)** m/z calculated for $C_{18}H_{32}O_2$: 280.240; found: 281.245 $[M+H]^+$.
**Yield:** 1.3 g (4.6 mmol, 90 %).

#### *6-(3-(Undec-10-yn-1-yl)-3H-diazirin-3-yl)hexan-1-ol (13)*

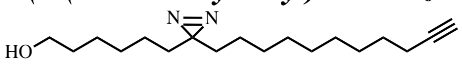


Molecular sieves (3Å, powder) were dried with heat under high vacuum. Ammonia (16 ml, 7N, solution in methanol) was then added and the solution was cooled to 0 ºC. **Compound 12** (920 mg, 3.40 mmol, 1 eq.) was dissolved in 12 ml dry MeOH and the solution added in dropwise fashion to the reaction mixture. After stirring for 1 h at 0 ºC, the ice bath was removed and the reaction mixture was stirred for an additional 4.5 h. Subsequently, $H_2NOSO_3H$ (860 mg, 7.6 mmol, 2.3 eq.) was dissolved in 10 ml dry MeOH and added dropwise at 0 ºC. The reaction mixture was stirred over night at room temperature. The molecular sieves were removed by filtration and the filtrate was evaporated *in vacuo.* The residue was redissolved in 35 ml MeOH, cooled to 0 ºC and 5 ml of $NEt_3$ were added dropwise. The reaction mixture was then allowed to reach room temperature. After 1 h, iodine was added in small batches at 0 ºC until the reaction mixture remained brownish. The reaction was quenched with sat. aqueous $Na_2S_2O_3$-solution. The reaction mixture was extracted with DCM (3x), the combined organic layers dired over $Na_2SO_4$, and the solvent removed under reduced pressure. The crude product was purified by flash chromatography (CyHex/EtOAc 8:2). The bifunctional alcohol **13** was isolated as a light yellowish oil.

**$^{1}$H NMR (400 MHz, $CD_3OD$)** δ = 3.53 (t, *J* = 6.6 Hz, 2H), 2.20 – 2.10 (m, 3H), 1.57 – 1.45 (m, 4H), 1.45 – 1.20 (m, 18H), 1.17 – 1.02 (m, 4H) ppm.
**$^{13}$C NMR{$^{1}$H} (101 MHz, $CD_3OD$)** δ = 85.05, 69.36, 62.87, 33.86, 33.83, 33.48, 30.49, 30.44, 30.27, 30.13, 29.75, 29.70, 29.54, 26.72, 24.90, 19.01 ppm.
**HR-MS (ESI positive)** calculated for $C_{18}H_{32}N_2O$: 292.252; found: 310.282 $[M+NH_4]^+$.
**Yield:** 160 mg (547 µmol, 16 %).

#### *(R)-1-((Triisopropylsilyl)oxy)-3-((6-(3-(undec-10-yn-1-yl)-3H-diazirin-3-yl)hexyl)oxy)propan-2-ol (15)*

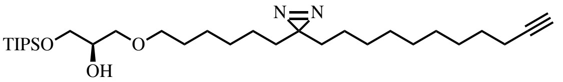


A solution of bifunctional alcohol **13** (96.0 mg, 328 µmol, 1.0 eq.) in 2 ml dry DCM was treated with TIPS-(S)-glycidol (**14**) (265 mg, 1.15 mmol, 3.5 eq.) and $Sc(OTf)_3$ (8.0 mg, 16 µmol, 0.05 eq.) under an argon atmosphere. The reaction mixture was stirred overnight at room temperature. After adding more $Sc(OTf)_3$ (8.0 mg, 16.3 µmol, 0.05 eq.) the reaction mixture was stirred for one more day. The solvent was removed under reduced pressure and the crude product was purified by flash chromatography (CyHex/EtOAc 95:5). **Compound 15** was isolated as a colourless oil.

**$^1$H NMR (400 MHz, $CDCl_3$)** δ = 3.85 – 3.77 (m, 1H), 3.77 – 3.66 (m, 2H), 3.50 – 3.38 (m, 4H), 2.16 (td, *J* = 7.1, 2.7 Hz, 2H), 1.92 (t, *J* = 5.4 Hz, 1H), 1.60 – 1.43 (m, 4H), 1.41 – 1.15 (m, 18H), 1.15 – 1.00 (m, 25H) ppm.
**$^{13}$C NMR{$^1$H} (101 MHz, $CDCl_3$)** δ = 84.87, 71.58, 71.54, 70.86, 68.19, 64.40, 33.01, 32.98, 29.62, 29.46, 29.44, 29.32, 29.18, 29.15, 28.97, 28.83, 28.58, 26.06, 23.97, 23.94, 18.51, 18.06, 12.00 ppm.
**HR-MS (ESI positive)** calculated for $C_{30}H_{58}N_2O_3Si$: 522.422; found: 540.456 $[M+NH_4]^+$.
**Yield:** 101 mg (193 µmol, 59 %).*
* The bifunctional alcohol can be reisolated and reused after flash chromatography.

#### *(R)-1-((Triisopropylsilyl)oxy)-3-((6-(3-(undec-10-yn-1-yl)-3H-diazirin-3-yl)hexyl)oxy)propan-2-yl oleate (16)*

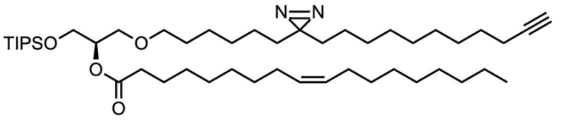


A solution of oleic acid (65.0 mg, 230 µmol, 1.2 eq.) in 1 ml dry DCM was treated with EDC·HCl (90 mg, 0.47 mmol, 2.4 eq.) and DMAP (12.0 mg, 98.2 µmol, 0.5 eq.) and stirred for 15 min. A solution of **compound 15** (101 mg, 193 µmol, 1.0 eq.) in 1 ml dry DCM was treated with the activated oleic acid solution in a dropwise fashion at 0 ºC. The reaction mixture was stirred over night at room temperature. The solvent was removed under reduced pressure and the crude product was purified by flash chromatography (CyHex/EtOAc, 98:2). **Compound 16** was isolated as a yellowish oil.

**$^1$H NMR (400 MHz, $CDCl_3$)** δ = 5.32 ($m_c$, 2H), 5.01 ($m_c$, 1H), 3.80 ($m_c$, 2H), 3.56 ($m_c$, 2H), 3.49 – 3.32 (m, 2H), 2.29 (t, *J* = 7.6 Hz, 2H), 2.16 (td, *J* = 7.1, 2.6 Hz, 2H), 2.06 – 1.93 (m, 4H), 1.93 – 1.88 (m, 1H), 1.68 – 1.56 (m, 2H), 1.56 – 1.42 (m, 4H), 1.42 – 1.14 (m, 38H), 1.11 – 0.99 (m, 25H), 0.86 (t, *J* = 6.7 Hz, 3H) ppm.
**$^{13}$C NMR{$^1$H} (101 MHz, $CDCl_3$)** δ = 173.42, 130.07, 129.85, 84.79, 73.14, 71.47, 69.03, 68.19, 62.13, 34.55, 33.00, 32.03, 29.89, 29.84, 29.65, 29.59, 29.45, 29.32, 29.24, 29.22, 29.17, 29.15, 28.90, 28.82, 28.57, 27.33, 27.30, 26.02, 25.09, 23.96, 23.95, 22.81, 18.50, 18.21, 18.02, 14.24, 12.00 ppm.
**HR-MS (ESI positive)** calculated for $C_{48}H_{90}N_2O_4Si$: 786.667; found: 804.699 $[M+NH_4]^+$.
**Yield:** 87.6 mg (111 µmol, 57 %).*
* The starting material can be reisolated and reused after flash chromatography.

#### *(S)-1-Hydroxy-3-((6-(3-(undec-10-yn-1-yl)-3H-diazirin-3-yl)hexyl)oxy)propan-2-yl oleate (17)*

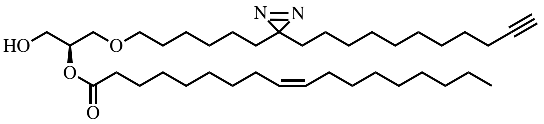


A solution of **Compound 16** (85.0 mg, 108 µmol, 1.0 eq.) in 3 ml dry THF was treated with 60 µl of acetic acid (59.9 µmol, 9.7 eq.). The mixture was cooled to 0 ºC and TBAF (solution in dry THF, 100 µl, 346 µmol, 3.2 eq.) were added dropwise. The reaction mixture was stirred over night at room temperature. The solvent was removed and the crude product was purified by flash chromatography (CyHex/EtOAc, 85:15). **Compound 17** was isolated as a colourless oil.

**$^{1}$H NMR (400 MHz, $CDCl_3$)** δ = 5.34 ($m_c$, 2H), 4.99 ($m_c$, 1H), 3.89 – 3.71 (m, 2H), 3.60 ($m_c$ 2H), 3.43 ($m_c$, 2H), 2.35 (t, *J* = 7.6 Hz, 2H), 2.17 (td, *J* = 7.1, 2.6 Hz, 2H), 2.07 – 1.95 (m, 4H), 1.94 ($m_c$, 1H), 1.68 – 1.46 (m, 6H), 1.44 – 1.13 (m, 38H), 1.13 – 0.99 (m, 4H), 0.88 (t, *J* = 6.7 Hz, 3H) ppm.
**$^{13}$C NMR{$^{1}$H} (101 MHz, $CDCl_3$)** δ = 173.82, 130.16, 129.87, 84.91, 72.91, 71.85, 70.12, 68.21, 63.13, 34.51, 33.03, 32.98, 32.05, 29.91, 29.84, 29.67, 29.53, 29.47, 29.33, 29.25, 29.22, 29.17, 29.14, 28.98, 28.85, 28.60, 27.37, 27.31, 25.99, 25.11, 23.99, 23.95, 22.83, 18.53, 14.27 ppm.
**HR-MS (ESI positive)** calculated for $C_{39}H_{70}N_2O_4$: 630.533; found: 648.563 $[M+NH_4]^+$.
**Yield:** 22.0 mg (34.9 µmol, 32 %).*
* The starting material can be reisolated and reused after flash chromatography.

#### *ePC(Y18/18:1) (1)*

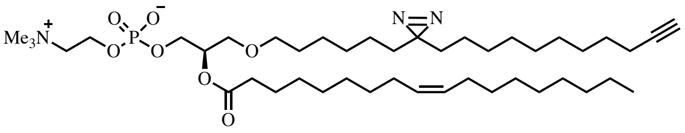


In a 25 ml flask molecular sieves (3Å, powder) were activated with heat under vacuum. A solution of **compound 17** (22.0 mg, 34.9 µmol, 1 eq.) in 3 ml dry MeCN was added. The phosphoramidite **18** (37.0 mg, 59.3 µmol, 1.7 eq.) was dissolved in 2 ml dry MeCN and added. 0.09 ml of a tetrazole solution (0.45 M, 40.4 µmol, 1.2 eq.) were added dropwise at room temperature. The reaction mixture was stirred for 3.5 h at room temperature and subsequently cooled to 0 °C. 3 drops of $^{t}$BuOOH (5.5M in decane, ca. 2 eq.) were added and the reaction mixture was stirred for 1.5 h and allowed to warm up to room temperature. After removing all solids by filtration, the filtrate was concentrated under reduced pressure by co-evaporation with toluene. The remaining solid was dissolved in 2 ml dry DCM. $NEt_3$ (25 µl, 180 µmol, 5.1 eq.) was added and the reaction mixture was stirred for 20 h at room temperature. 10 ml of DCM were added and the organic phase was washed with 1N HCl (2x), sat. $NaHCO_3$-solution (1x) and brine (1x). The combined aqueous phases were reextracted with DCM once. The combined organic phases were concentrated under reduced pressure. The crude product was purified by flash chromatography ($CHCl_3$/MeOH/$H_2O$, 70:30:4). ePC($Y_{18}$/18:1) (**1**) was isolated as a yellowish oil.

**$^{1}$H NMR (400 MHz, $CD_3OD$)** δ = 5.35 ($m_c$, 2H), 5.16 ($m_c$, 1H), 4.34 – 4.20 (m, 2H), 4.07 – 3.91 (m, 2H), 3.69 – 3.53 (m, 4H), 3.53 – 3.37 (m, 2H), 3.23 (s, 9H), 2.35 (t, *J* = 7.4 Hz, 2H), 2.21 – 2.10 (m, 3H), 2.10 – 1.97 (m, 4H), 1.69 – 1.45 (m, 6H), 1.44 – 1.16 (m, 38H), 1.15 – 1.01 (m, 4H), 0.90 (t, *J* = 6.6 Hz, 3H) ppm.
**$^{13}$C NMR{$^{1}$H} (101 MHz, $CD_3OD$)** δ = 174.86, 130.91, 130.79, 85.04, 73.01, 72.41, 70.34, 69.41, 67.43, 65.36, 60.48, 54.72, 54.68, 54.65, 35.23, 33.88, 33.85, 33.08, 30.86, 30.63, 30.59, 30.50, 30.47, 30.37, 30.28, 30.26, 30.18, 30.15, 30.13, 29.76, 29.71, 29.55, 28.16, 27.03, 26.11, 24.95, 24.93, 23.76, 19.02, 14.49 ppm.
**$^{31}$P NMR{$^{1}$H} (162 MHz, $CD_3OD$)** δ = -0.55 ppm.
**HR-MS (ESI positive)** calculated for $C_{44}H_{82}N_3O_7P$: 795.589; found: 796.592 $[M+H]^+$.
**Yield:** 13.1 mg (16.5 µmol, 47 %).

#### Synthesis of PC($Y_{18}$/18:1) (2)

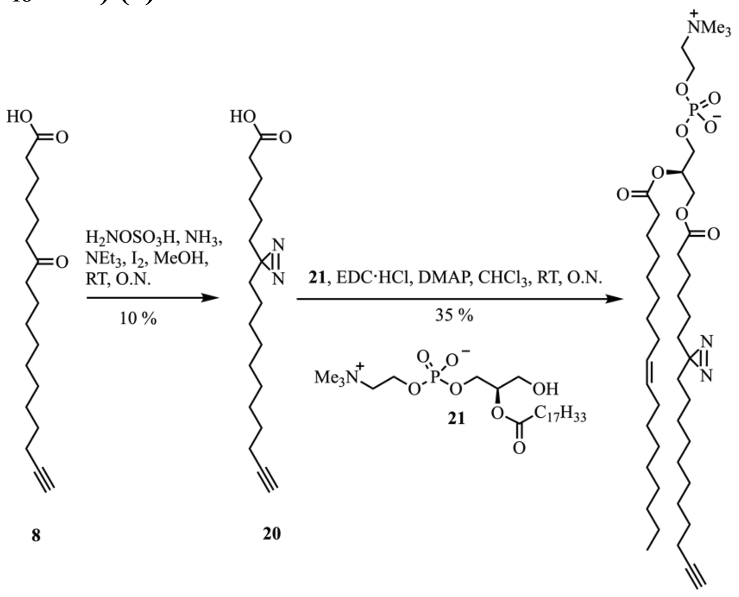


**Scheme B**: Synthesis of PC($Y_{18}$/18:1) (**2**). Lyso PC **21** was synthesized according to a recently published protocol.[1] The analytical data was in accordance with the ones published. Detailed experimental procedures for coumpounds **20** and **1** are described below.

##### *6-(3-Uundec-10-yn-1-yl)-3H-diazirin-3-yl)hexanoic acid (20)*

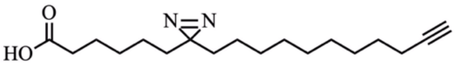


In a 250 ml flask molecular sieves (3Å, powder) were activated with heat under high vacuum. 18 ml of ammonia (7N in MeOH) were added. **Compound 8** (1.0 g, 3.4 mmol, 1.0 eq.) was dissolved in 13 ml dry MeOH and added dropwise. The reaction mixture was stirred at rt. for 5.5 h. 847 mg of hydroxylamine-O-sulfonic acid (7.5 mmol, 2.2 eq.) (dried under high vacuum for 3 h) was dissolved in 13 ml dry MeOH and the solution was added dropwise at 0 °C to the reaction mixture. The reaction mixture was stirred at room temperature overnight. Molecular sieves were removed by filtration. The filtrate was concentrated under reduced pressure and the residue was dissolved in 35 ml MeOH. 5 ml $NEt_3$ were added dropwise at 0 °C and the mixture was stirred at room temperature for 1 h. Subsequently, iodine was added in small batches at 0 °C until the mixture remained brownish. Sat. aqueous $Na_2S_2O_3$-solution was added to remove excess iodine. After extraction with DCM (3x), the crude product was purified by flash chromatography (DCM/MeOH, 95:5). The product containing fraction was purified further by HPLC using a Macherey Nagel VP 250/21 Nucleodur 100-5 C8ec column at 8 ml/min eluting with a gradient The solvent system used was A (75 % MeOH, 25 %water, + 4 % AcOH) and B (50 % MeCN, 40 % i-propanol, 10 % MeOH, + 4 % AcOH). Gradient: 0-2 min: 0-45 % B, 2-8 min: 45-75 % B, 8-11 min: 75-100 % B, 11-27 min: 100-0% B, 27-30 min: 0 % B. The retention time of the product was found to be 22.1 min. Bifunctional fatty acid **20** was isolated as a light yellowish oil.

**$^1$H NMR (400 MHz, MeOD)** δ = 2.27 (t, *J* = 7.4 Hz, 2H), 2.19 – 2.11 (m, 3H), 1.62 – 1.45 (m, 4H), 1.45 – 1.18 (m, 16H), 1.18 – 1.02 (m, 4H) ppm.
**$^{13}$C NMR{$^1$H} (101 MHz, MeOD)** δ = 177.38, 85.03, 69.39, 34.74, 33.80, 33.74, 30.48, 30.44, 30.26, 30.13, 29.76, 29.75, 29.68, 29.47, 25.80, 24.90, 24.63, 19.03 ppm.
**HR-MS (ESI negative)** m/z calculated for $C_{18}H_{30}N_2O_2$: 306.231; found: 305.223 $[M-H]^-$.
**Yield:** 99.0 mg (0.32 mmol, 9 %).

#### *PC(Y₁₈/18:1) (2)*

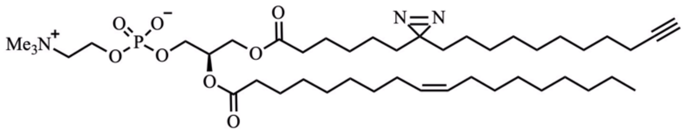


A solution of the bifunctional fatty acid **20** (40.0 mg, 130.5 µmol, 1.7 eq.) in 0.5 ml $CDCl_3$ was treated with EDC·HCl (47.0 mg, 245.2 µmol, 3.2 eq.) and DMAP (5.0 mg, 40.9 µmol, 0.5 eq.) for 15 minutes at room temperature. The solution of activated bifunctional fatty acid **20** was then added to a solution of lyso-PC **21** (40.0 mg, 76.7 µmol, 1.0 eq.) in 1 ml $CDCl_3$. The reaction mixture was stirred overnight at room temperature. After removing the solvents under reduced pressure, the crude product was purified by flash chromatography ($CHCl_3$/MeOH/$H_2O$, 70:30:2). PC($Y_{18}$/18:1) (**2**) was isolated as a colourless oil.

**$^1$H NMR (400 MHz, MeOD)** δ = 5.35 ($m_c$, 2H), 5.24 ($m_c$, 1H), 4.44 (dd, *J* = 12.0, 3.0 Hz, 1H), 4.28 ($m_c$, 2H), 4.17 (dd, *J* = 12.0, 6.9 Hz, 1H), 4.00 (t, *J* = 6.0 Hz, 2H), 3.69 – 3.59 (m, 2H), 3.23 (s, 9H), 2.35 (t, *J* = 7.6 Hz, 2H), 2.31 (t, *J* = 7.4 Hz, 2H), 2.20 – 2.10 (m, 3H), 2.10 – 1.97 (m, 4H), 1.70 – 1.44 (m, 6H), 1.44 – 1.20 (m, 36H), 1.16 – 1.02 (m, *J* = 14.8, 7.6 Hz, 4H), 0.91 (t, *J* = 6.6 Hz, 3H) ppm.*[1]
*Signals (triplets) at 2.35 and 2.31 ppm overlap.
**$^{13}$C NMR{$^1$H} (101 MHz, MeOD)** δ = 174.70, 174.55, 130.93, 130.78, 85.05, 71.81, 69.42, 67.49, 64.90, 63.71, 60.49, 54.68, 35.08, 34.72, 33.86, 33.75, 33.09, 30.87, 30.64, 30.52, 30.48, 30.38, 30.30, 30.27, 30.19, 30.17, 29.77, 29.72, 29.51, 28.18, 26.04, 25.74, 24.94, 24.66, 23.76, 19.03, 14.50 ppm.*[2,3]
*[2] Signals between 29.7 and 30.9 ppm overlap, not all signals are resolved individually.
*[3] Signals at 71.8, 67.5, 64.9, 60.5 and 54.7 ppm appear as multiplets likely due to $^{13}$C-$^{31}$P coupling.
**$^{31}$P NMR{$^1$H} (162 MHz, MeOD)** δ = -0.57 ppm.
**HR-MS (ESI positive)** m/z calculated for $C_{44}H_{80}N_3O_8P$: 809.568; found: 810.588 $[M+H]^+$.
**Yield:** 21.7 mg (26.8 µmol, 35 %).

#### Synthesis of *sn2*-bifunctional lipids

PC(18:1/$Y_{16}$) (**4**) and 6-(3-(non-8-yn-1-yl)-3*H*-diazirin-3-yl)hexanoic acid were synthesized according to a published protocol.[1] The analytical data was in accordance with those reported. Detailed experimental procedures for the synthesis of ePC(18:1/$Y_{16}$) and pPC(18:1/$Y_{16}$) are described below.

#### ePC(18:1/Y₁₆) (3)

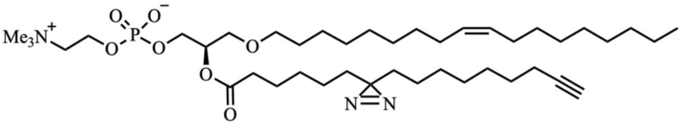


A solution of 6-(3-(non-8-yn-1-yl)-3*H*-diazirin-3-yl)hexanoic acid (70.0 mg, 251 µmol, 2.0 eq.) in 1 ml $CDCl_3$ was treated with EDC·HCl (67.0 mg, 349 µmol, 2.7 eq.) and 8.0 mg DMAP (8.0 mg, 66 µmol, 0.5 eq.) for 10 minutes at room temperature. The solution of activated bifunctional fatty acid was then added to a solution of C18:1 lyso-PC (65.0 mg, 128 µmol, 1.0 eq.) in 1 ml $CDCl_3$. The reaction mixture was stirred overnight at room temperature. After removing the solvents under reduced pressure, the crude product was purified twice by flash chromatography ($CHCl_3$/MeOH/$H_2O$, 70:30:3). ePC(18:1/Y) (**3**) was isolated as a colourless oil.

**$^1$H NMR (400 MHz, MeOD)** δ = 5.44 – 5.28 (m, 2H), 5.21 – 5.08 (m, 1H), 4.33 – 4.18 (b, 2H), 4.08 – 3.89 (m, 2H), 3.68 – 3.57 (m, 4H), 3.55 – 3.36 (m, 2H), 3.23 (s, 9H), 2.34 (t, *J* = 7.3 Hz, 2H), 2.21 – 2.11 (m, 3H), 2.09 – 1.96 (m, 4H), 1.66 – 1.44 (m, 6H), 1.44 – 1.21 (m, 34H), 1.12 (dd, *J* = 14.1, 6.8 Hz, 4H), 0.90 (t, *J* = 6.9 Hz, 3H) ppm.
**$^{13}$C NMR{$^1$H} (101 MHz, MeOD)** δ = 174.75, 130.89, 130.84, 85.03, 73.16, 73.08, 72.58, 70.33, 69.43, 65.33, 60.45, 60.40, 54.74, 54.71, 54.67, 35.04, 33.83, 33.74, 33.07, 30.88, 30.84, 30.76, 30.67, 30.61, 30.46, 30.35, 30.19, 29.96, 29.69, 29.63, 29.62, 29.52, 28.14, 27.23, 25.83, 24.86, 24.66, 23.75, 18.99, 14.47, 1.47 ppm.
**$^{31}$P NMR{$^1$H} (162 MHz, MeOD)** δ = -0.51 ppm.
**HR-MS (ESI positive):** calculated m/z for $C_{42}H_{78}N_3O_7P$: 767.557, found: 768.575 $[M+H]^+$.
**Yield:** 50.7 mg (66.0 µmol, 52 %).

#### pPC(18:1/$Y_{16}$) (5)

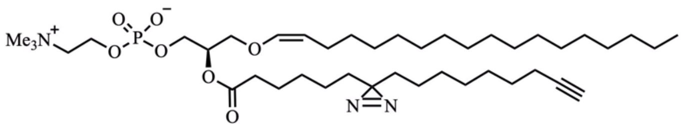

A solution of 6-(3-(non-8-yn-1-yl)-3*H*-diazirin-3-yl)hexanoic acid (10.0 mg, 35.9 µmol, 1.2 eq.) in 0.6 ml $CDCl_3$ was treated with 10.3 mg EDC·HCl (53.7 µmol, 1.8 eq.) and 2.2 mg of DMAP (18.0 µmol, 0.6 eq.) and stirred for 15 min. The activated fatty acid was then added to a solution of C18(Plasm)LPC (15.0 mg, 29.5 µmol, 1.0 eq.) in1 ml $CDCl_3$ under argon and the reaction mixture was stirred overnight at room temperature. The solvents were removed under reduced pressure and the crude product was purified twice by flash chromatography ($CHCl_3$/MeOH/$H_2O$, 70:30:3). pPC(18:1/$Y_{16}$) (**5**) was isolated as a colourless oil.

**$^1$H NMR (400 MHz, MeOD)** δ = 5.98 (d, *J* = 6.2 Hz, 1H), 5.24 – 5.12 (m, 1H), 4.40 – 4.30 (m, 1H), 4.30 – 4.19 (b, 2H), 4.08 – 3.95 (m, 2H), 3.93 – 3.82 (m, 2H), 3.69 – 3.55 (m, 2H), 3.23 (s, 9H), 2.34 (t, *J* = 7.4 Hz, 2H), 2.24 – 2.11 (m, 3H), 2.09 – 1.93 (m, *J* = 6.6 Hz, 2H), 1.67 – 1.44 (m, 4H), 1.44 – 1.18 (m, 40H), 1.18 – 1.00 (m, 4H), 0.90 (t, *J* = 6.7 Hz, 3H) ppm.
**$^{13}$C NMR{$^1$H} (101 MHz, MeOD)** δ = 174.52, 146.25, 108.33, 85.01, 73.10, 73.02, 71.48, 69.42, 64.93, 64.87, 60.48, 60.43, 54.74, 54.71, 54.67, 35.04, 33.85, 33.74, 33.09, 30.90, 30.83, 30.81, 30.77, 30.71, 30.49, 30.46, 30.20, 29.98, 29.73, 29.64, 29.51, 25.81, 24.94, 24.88, 24.68, 23.74, 19.00, 14.45 ppm.
**$^{31}$P NMR{$^1$H} (162 MHz, MeOD)** δ = -0.56 ppm.
**HR-MS (ESI positive):** calculated m/z for $C_{42}H_{78}N_3O_7P$: 767.557, found: 768.572 $[M+H]^+$.
**Yield:** 7.4 mg (9.6 µmol, 32 %)

#### Test reactions for the synthesis of the bifunctional long chain alcohol

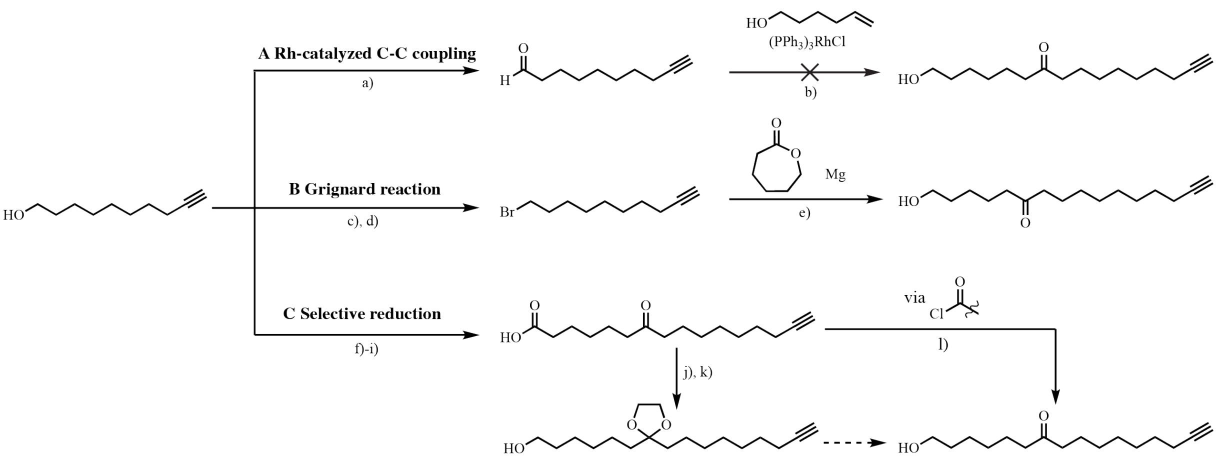


**Scheme C: Overview of tested reactions to obtain 1-hydroxyhexadec-15-yn-7-one or 1-Hydroxyhexadec-15-yn-6-one from 9-Decynol. a)** 1.1 eq. DMP, 1.1 eq. $H_2O$, DCM, 2.5 h, 84 %. **b)** 3 eq. 5-hexen-1-ol, 0.06 eq. $(PPh_3)_3RhCl$, 0.4 eq. 2-amino-3-picoline, 110 °C, O.N, no isol. . **c)** 1.2 eq. MsCl, 2.1 eq. $NEt_3$, dry DCM, 0 °C to r.t, 4.5 h, quant. . **d)** 1.3 eq. NaBr, dry MeCN, 90 °C, 21 h, 86 %. **e)** 7.3 eq. 10-bromodec-1-yne, 1 eq. ε-caporlactone, 1.2 eq. HN(OMe)Me·HCl, 0.3 eq. NaOMe, 8.3 eq. Mg, dry THF, -78 °C, 1 h, 8 %. **f)-i)** accoridng to Iglesias et al.[1] **j)** 1.6 eq. ethyleneglycol, 0.1 eq. TsOH·HCl, toluene, reflux, O.N. **k)** 1.8 eq. $LiAlH_4$, dry THF, r.t., 2.5 h, <14% over 2 steps. **l)** see **Supplementary Table 2**.

We first tried a Rh-catalyzed CC-coupling which did not yield the desired productin our hands.[8] A Grignard coupling only gave poor yields (<7 %) in test reactions.[9] Instead, we synthesized a long chain fatty acid according to a recently published protocol by Iglesias et. al.[1] and attempted to selectively reduce the carboxylic acid in presence of the ketone yielding the final product. We first attempted to activate the carboxylic acid with oxalyl chloride and subsequently used a mild reducing reagent (**Supplementary Table 2**). Treatment with $NaB(OAc)_3$ did not result in product formation (Entry 1+2). Switching to $NBu_4BH_4$ allowed us to isolate the product in acceptable yields (Entry 4-7). However, this reaction gave much worse yields in bigger batches (Entry 9) We also attempted a route via ketone acetalization with ethylene glycol and TsOH. After purification, we always isolated the product together with the hemiketal. The mix was used for $LiAlH_4$ reduction of the carboxylic acid but resulted in an overall very low yield (< 14% over 2 steps). Acetalization with ethylene glycol and TMOF (according to Kirar et al. [10]) did not result in product formation.

**Supplementary Table 2: Reaction conditions for the selective reduction of 7-oxohexadec-15-ynoic acid**

| Entry | batch size | reaction conditions | yield (12) |
|---|---|---|---|
| **1** | 101 mg | 1.6 eq. $(COCl)_2$, cat. DMF, DCM, 0 °C, 20 min<br>2 eq. **$NaB(OAc)_3H$**, DCM, 0 °C, O.N. | 0 % |
| **2** | 101 mg | 1.2 eq. $(COCl)_2$, cat. DMF, DCM, 0 °C, 20 min<br>1.5 eq. **freshly opened $NaB(OAc)_3H$**, DCM, 0 °C, O.N. | 0 % |
| **3** | 100 mg | 1.5 eq. $(COCl)_2$, cat. DMF, 0 °C, 20 min,<br>1.0 eq. **$NBu_4BH_4$**, 10 min | 0 % |
| **4** | 201 mg | 1.4 eq. $(COCl)_2$, cat. DMF, 0 °C, 15 min, r.t., 20 min, **molecular sieves**<br>1.0 eq. $NBu_4BH_4$, 0 °C, 10 min | 30 % |
| **5** | 100 mg | 1.6 eq. $(COCl)_2$, cat. DMF, 0 °C, 10 min, r.t., 20 min, molecular sieves<br>1.0 eq. $NBu_4BH_4$, 0 °C, **3 min** | 33 % |
| **6** | 302 mg | 1.3 eq. $(COCl)_2$, cat. DMF, 0 °C, 15 min, r.t., **45 min**, molecular sieves<br>1.0 eq. $NBu_4BH_4$, 0 °C, 3 min | 39 % |
| **7** | 200 m | 1.5 eq. $(COCl)_2$, cat. DMF, 0 °C, 15 min, r.t., 45 min, molecular sieves<br>1.0 eq. $NBu_4BH_4$, 0 °C, **1 min** | 32 % |
| **8** | 100 mg | 1.5 eq. $(COCl)_2$, cat. DMF, 0 °C, 15 min, r.t., 45 min, molecular sieves<br>1.0 eq. $NBu_4BH_4$, **-10 °C**, 1 min | 0 % |
| **9** | **2.0 g** | 1.4 eq. $(COCl)_2$, cat. DMF, 0 °C, 10 min, r.t., 30 min, molecular sieves<br>1.0 eq. $NBu_4BH_4$, 0 °C, 3 min | 6 % |

#### Optimization of epoxide opening reaction

The reaction conditions and yields of the test reactions for the epoxide opening step can be found in **Supplementary Table 3**.

Many different reagents are described in the literature for high regio- and stereoselective ring opening reaction of epoxides (e.g. Review [11]).
This reaction has also been reported as a key step in alkyl ether PC synthesis, using sodium hydride as the base.[12] In our hands, using NaH did not lead to product formation as the starting material was re-isolated (Entry 1-3). We then tested $BF_3{\cdot}OEt_2$, which was also reported to give high yields for long chain alcohols, but did not work either (Entry 4,5).[13][14] The use of $FeCl_3$ (solid or in solution) gave low yields (Entry 6-12). Overall, catalytic amounts of $Sc(OTf)_3$ gave the best results. We observed lower yields for the C16 bifunctional alcohol compared to 9-Decynol with the same reaction conditions (Entry 16, 17). The yield could be increased significantly by a longer reaction time and additional adding of the catalyst $Sc(OTf)_3$ after 1 day of stirring (Entry 18). We would like to note, that the use of catalytic $Al(OTf)_3$ as well as $InCl_3$ also resulted in product formation with slightly lower yields in contrast to $Sc(OTf)_3$ based on TLC analysis. After Entry 13 we changed the protecting group of (S)-glycidol from TBDMS to TIPS and used molecule D instead of C. This was due to the fact that the TBDMS group was partially cleaved in the following step (esterification of the *sn2*-alcohol with DMAP, EDC·HCl).

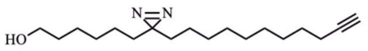

A

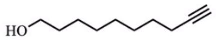

B

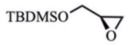

C

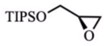

D

**Supplementary Table 3: Reaction conditions for the optimization of the epoxide opening reaction**

| **Entry** | **batch size** | **reaction conditions** | **yield** |
|---|---|---|---|
| **1** | 15 mg A | 1.3 eq. **C**, 3 eq. **NaH**, dry THF, 70 °C, O.N. | 0%, reisolated SM |
| **2** | 60 mg A | 1.3 eq. **C**, 2.7 eq. NaH, **dry THF**, **dry DMF**, 80 °C, O.N. | 0%, reisolated SM |
| **3** | 13 mg A | 1.6 eq. **C**, 2.5 eq. NaH, **dry DMF**, 50 °C, O.N. | 0 %, reisolated.SM |
| **4** | 10 mg A | 1.7 eq. **C**, 1 eq. **$BF_3{\cdot}OEt_2$**, dry DCM, r.t., O.N. | 0 %, reisolated SM |
| **5** | 42 mg B | 1 eq. **C**, **2.9 eq. $BF_3{\cdot}OEt_2$**, dry DCM, r.t., O.N. | < 2% |
| **6** | 42 mg B | 1 eq. C, **0.1 eq. $FeCl_3$ (s),** dry DCM, r.t., 1.5 h | 7 % |
| **7** | 50 mg B | 1 eq. C, **0.2 eq. $FeCl_3$ (s)**, dry DCM, r.t., **5 h** | 23 % |
| **8** | 40 mg | 1 eq. C, **0.1 eq. $FeCl_3$ (s)**, **dry chloroform**, r.t., 5 h | 21 % |
| **9** | 50 mg B | 1 eq. C, **1 eq. $FeCl_3$ (s)**, dry DCM, r.t., 27 h | 0 % |
| **10** | 55 mg B | 1 eq. C, 0.2 eq. $FeCl_3$ (s), dry DCM, **molecular sieves**, r.t. O.N. | 0 % |
| **11** | 54 mg B | 1 eq. C, **0.05 eq. $FeCl_3$ (0.2M in 2-methyl THF)**, dry DCM, r.t., O.N. | < 20% (TLC) |
| **12** | 54 mg B | 1 eq. C, 0.05 eq. $FeCl_3$ (0.2M in 2-methyl THF), **dry chloroform**, r.t., O.N. | < 20% (TLC) |
| **13** | 50 mg B | **5 eq. C**, **0.03 eq. $Sc(OTf)_3$**, dry DCM, r.t., O.N. | 68 % |
| **14** | **102 mg** B | 3.7 eq. **D**, 0.05 eq. $Sc(OTf)_3$, dry DCM, r.t., O.N. | 62 % |
| **15** | 102 mg B | 4 eq. D, **0.2 eq. $InCl_3$**, dry DCM, r.t., O.N. | < 60% (TLC) |
| **16** | **200 mg B** | 3.7 eq. **D**, 0.05 eq. $Sc(OTf)_3$, dry DCM, r.t., O.N. | 59 % |
| **17** | **109 mg A** | 3.7 eq. D, 0.05 eq. $Sc(OTf)_3$, dry DCM, r.t., O.N. | 29 % |
| **18** | 101 mg A | 3.4 eq. D, **0.05 eq. + 0.05 eq. $Sc(OTf)_3$**, dry DCM, r.t., **2 days** | 59 % |

### NMR spectra of new compounds

**Dodec-11-ynoic acid (19)**

*$^{1}H$ spectrum of 19*

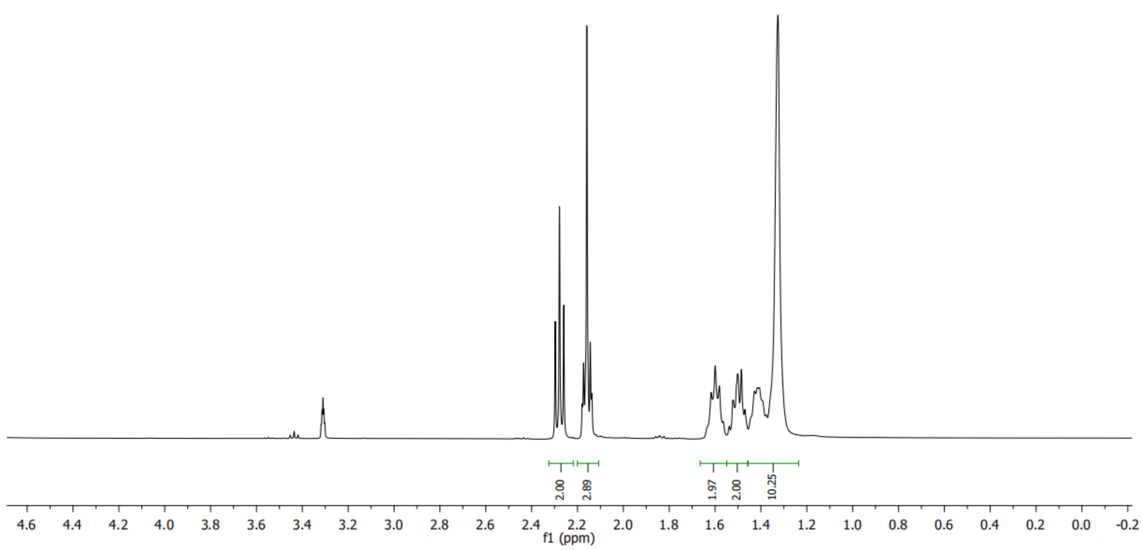

*$^{13}C$ spectrum of 19*

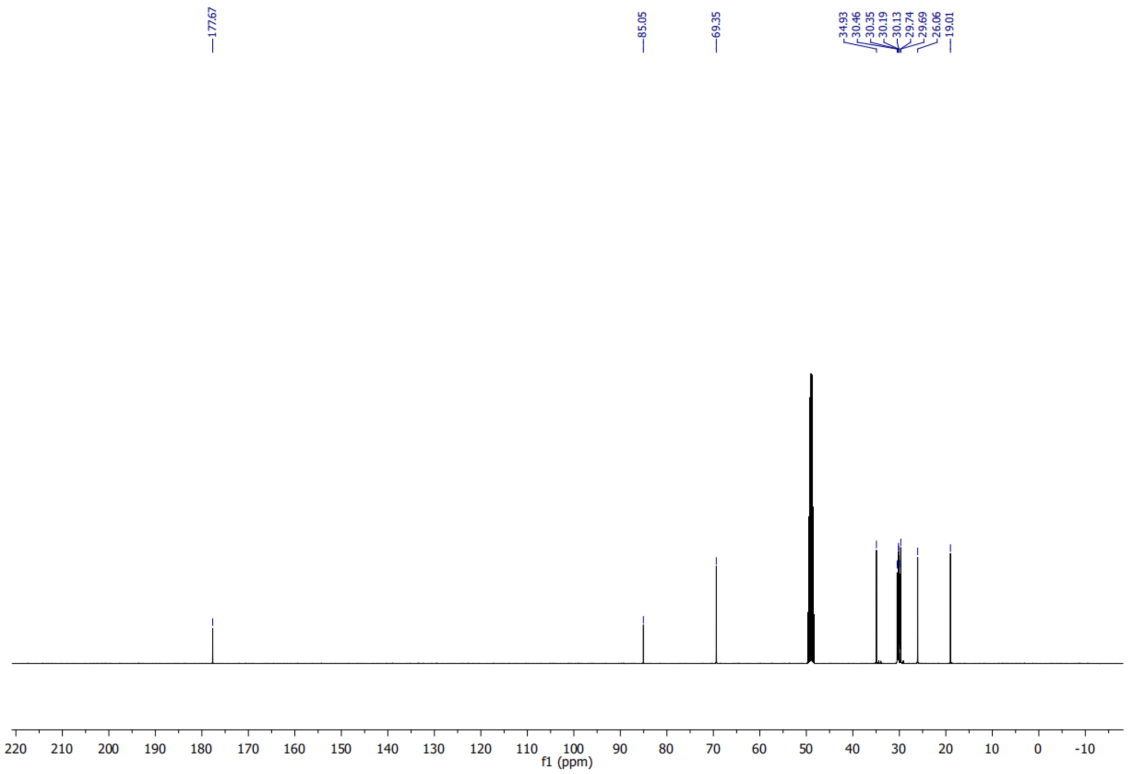

**1-(2-hydroxycyclohex-1-en-1-yl)dodec-11-yn-1-one (7)**

***[1]H spectrum of 7***

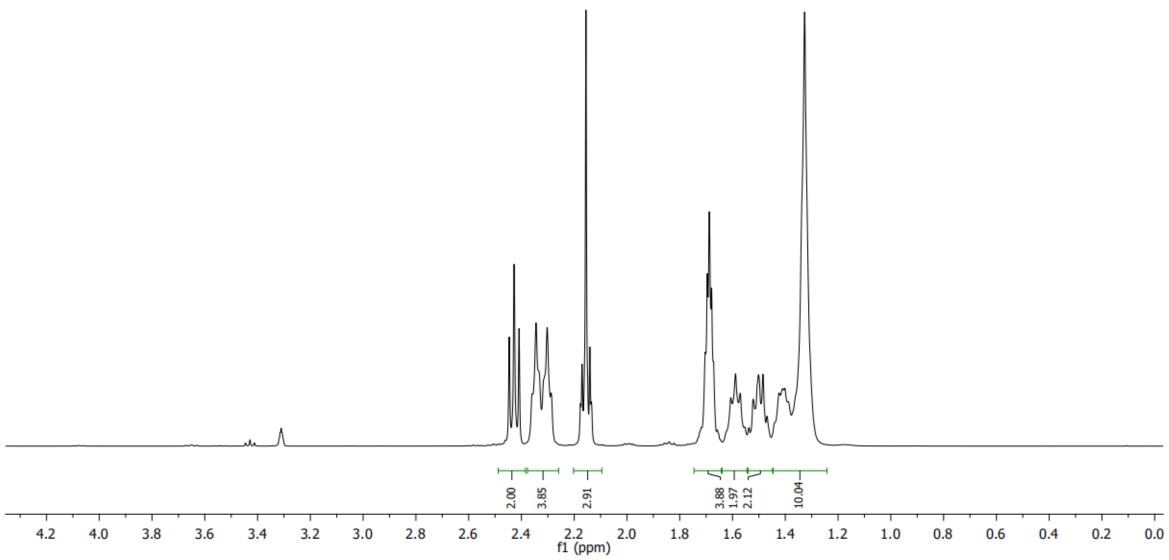

***[13]C spectrum of 7***

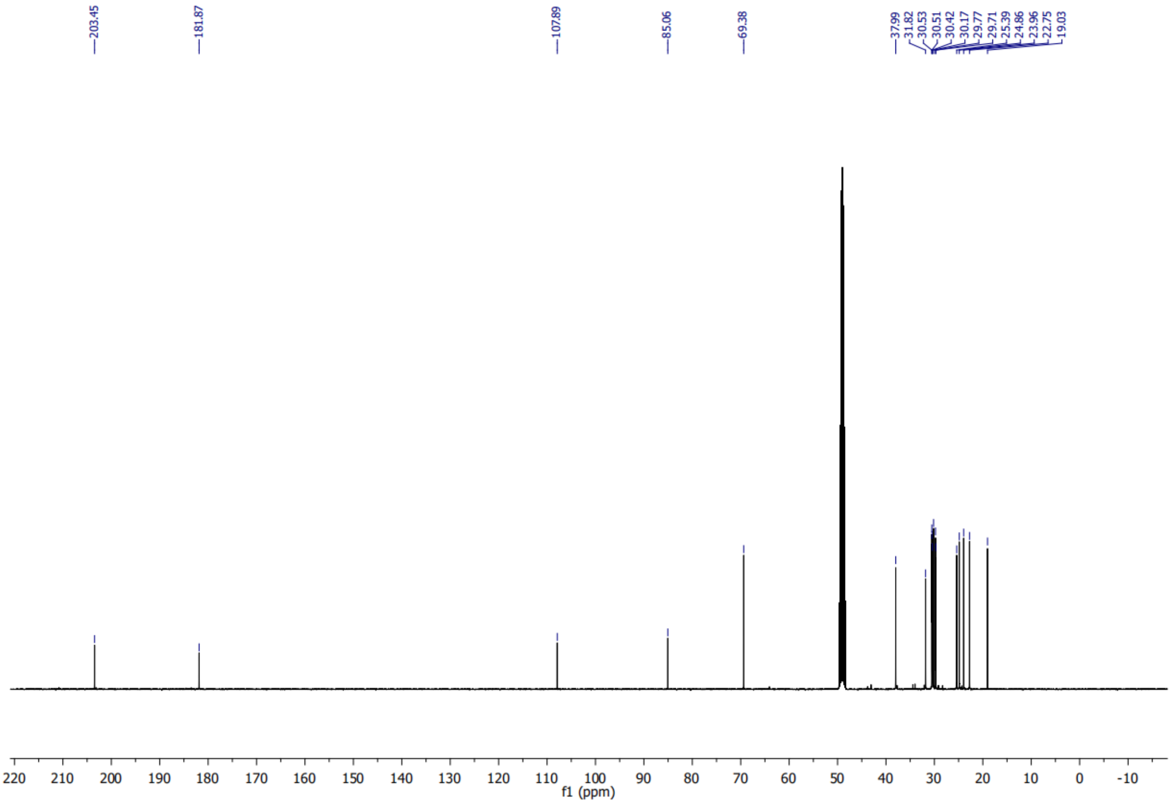

#### 7-oxooctadec-17-ynoic acid (8)

*[1]H spectrum of 8*

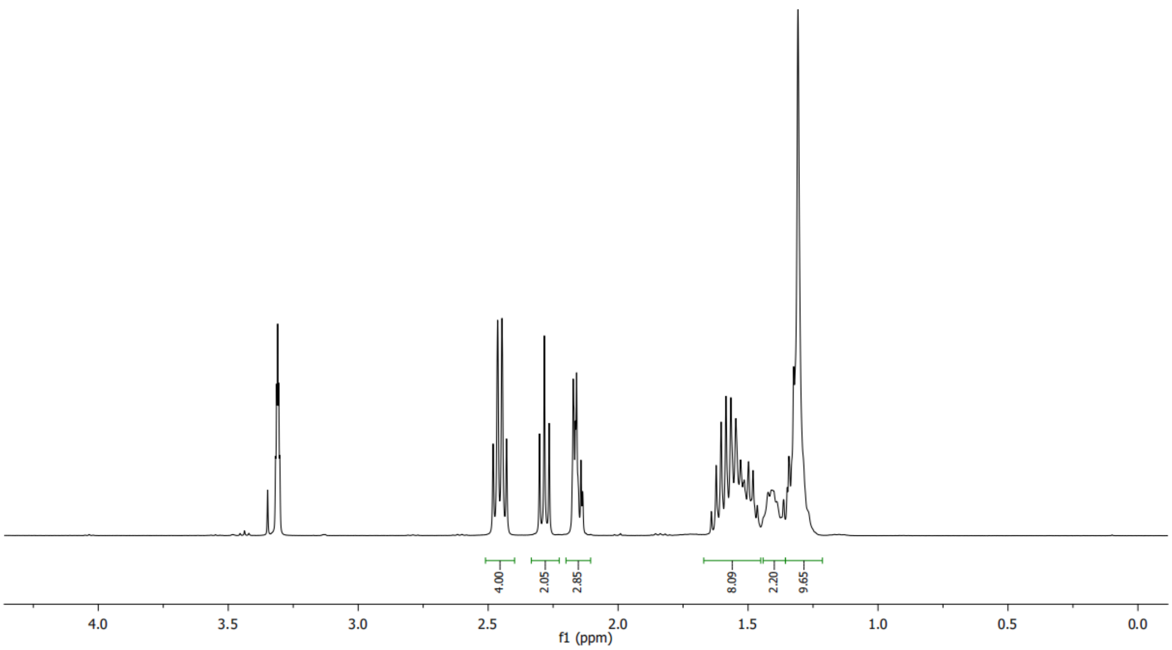


*[13]C spectrum of 8*

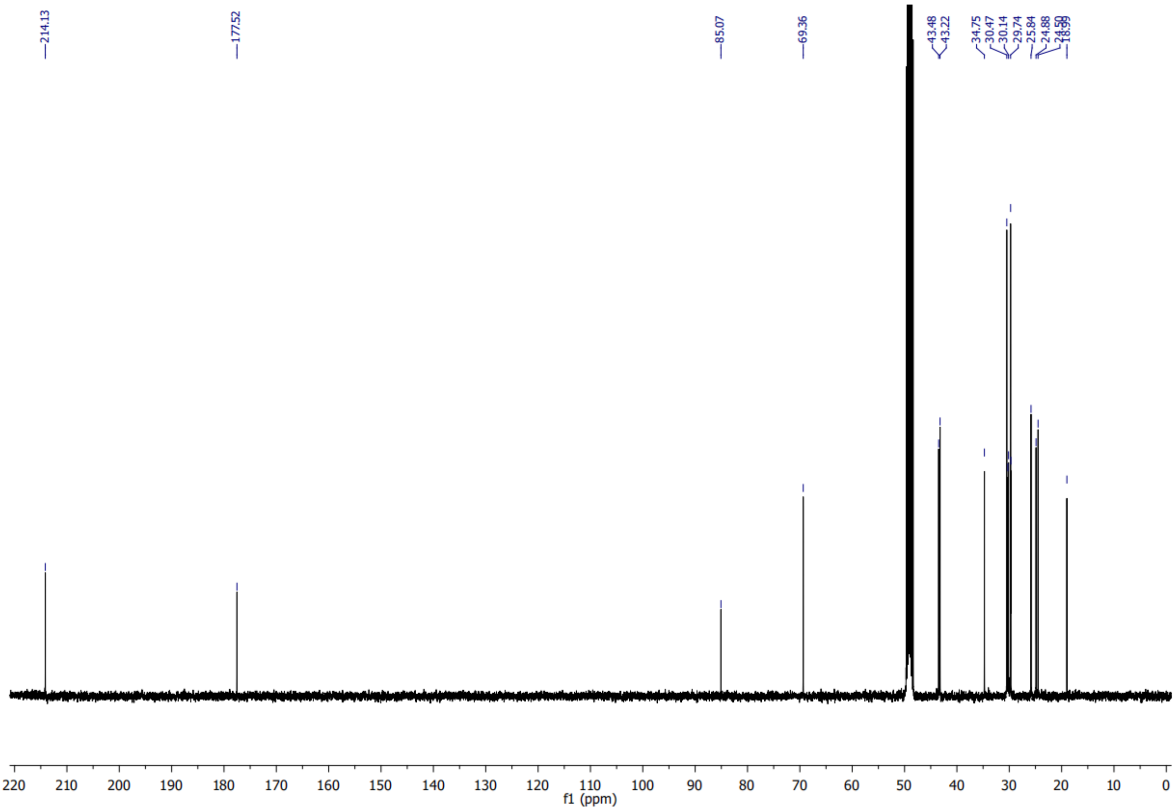

**Octadec-17-yne-1,7-diol (9)**

***$^{1}$H spectrum of 9***

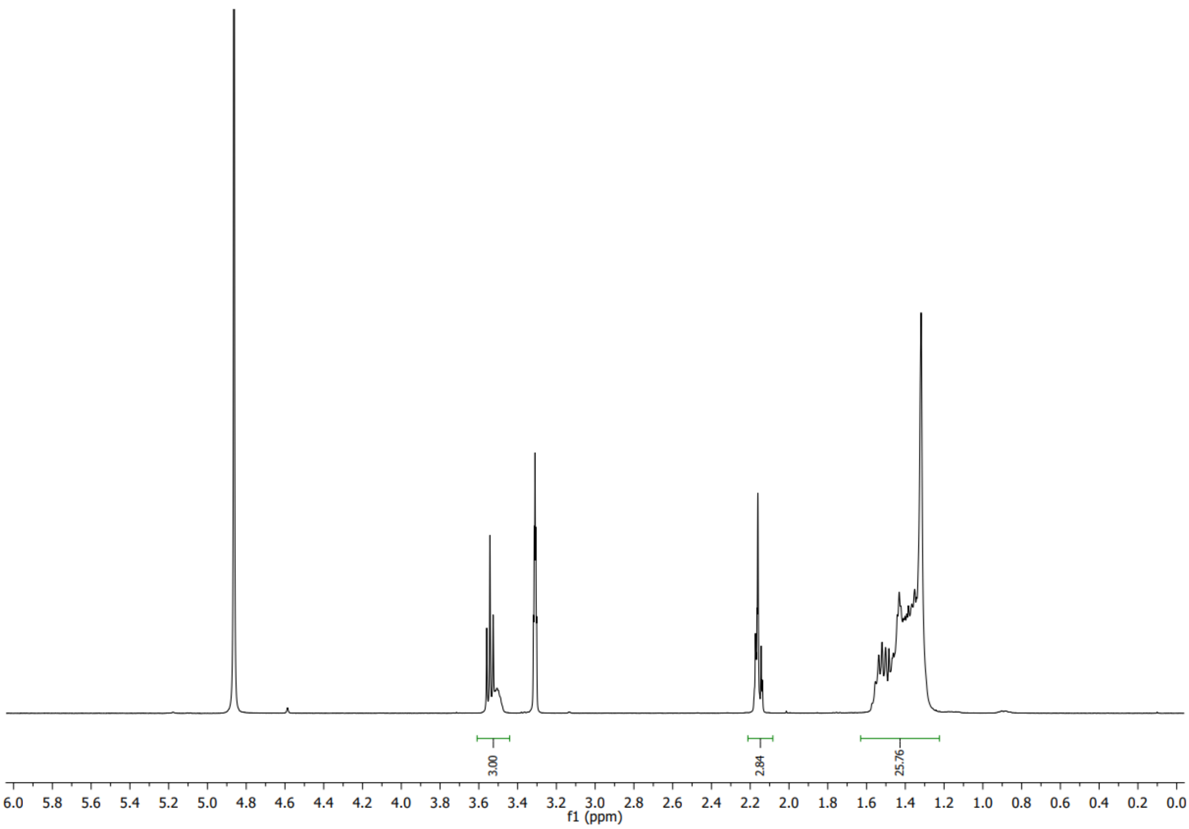


***$^{13}$C spectrum of 9***

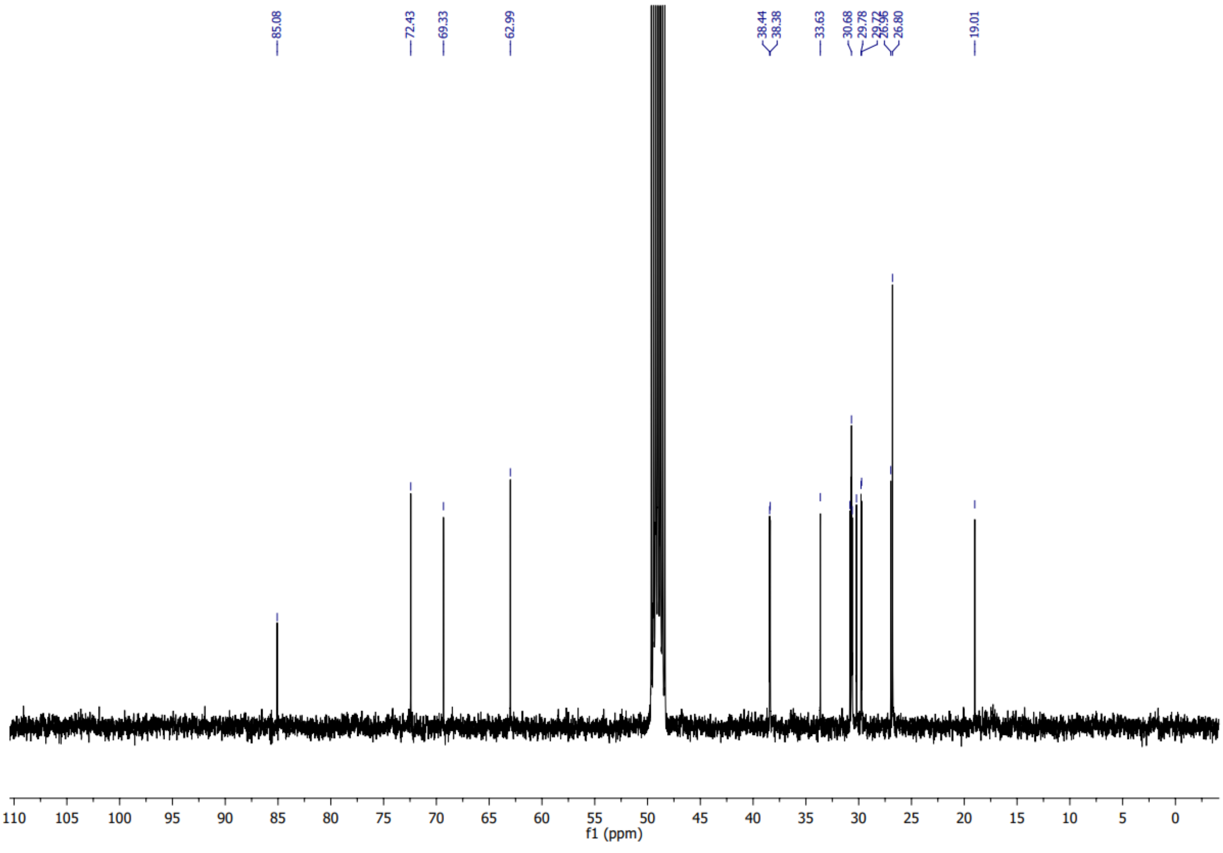

**1-(bis(4-methoxyphenyl)(phenyl)methoxy)octadec-17-yn-7-ol (10)**

##### *$^{1}$H spectrum of 10*

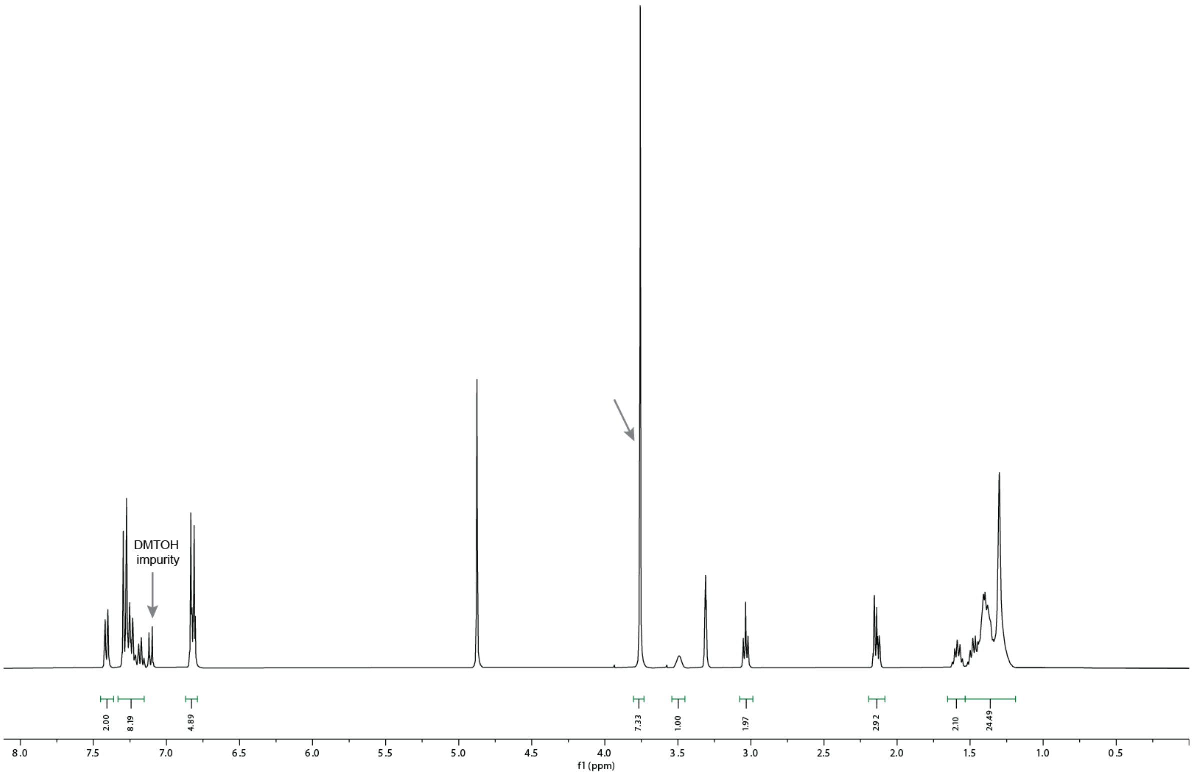


##### *$^{13}$C spectrum of 10*

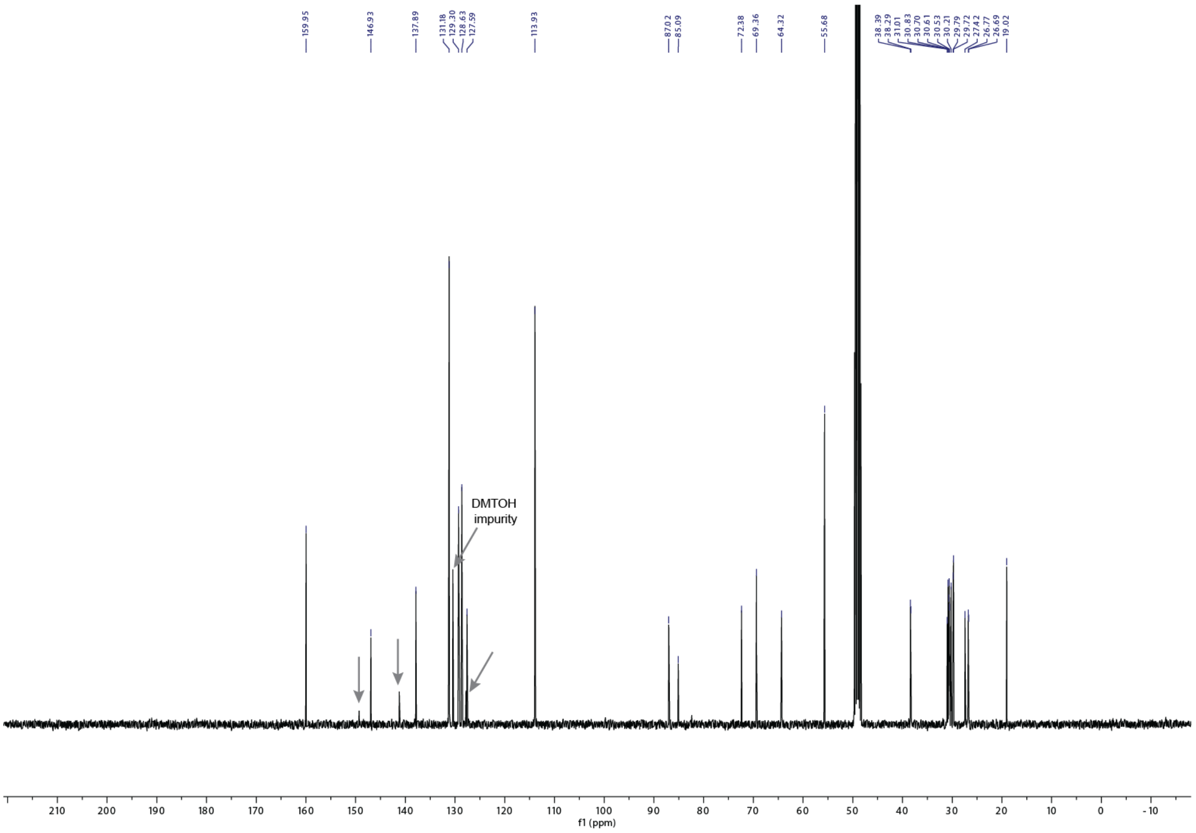

#### 1-(bis(4-methoxyphenyl)(phenyl)methoxy)octadec-17-yn-7-one (11)

##### *[1]H spectrum of 11*

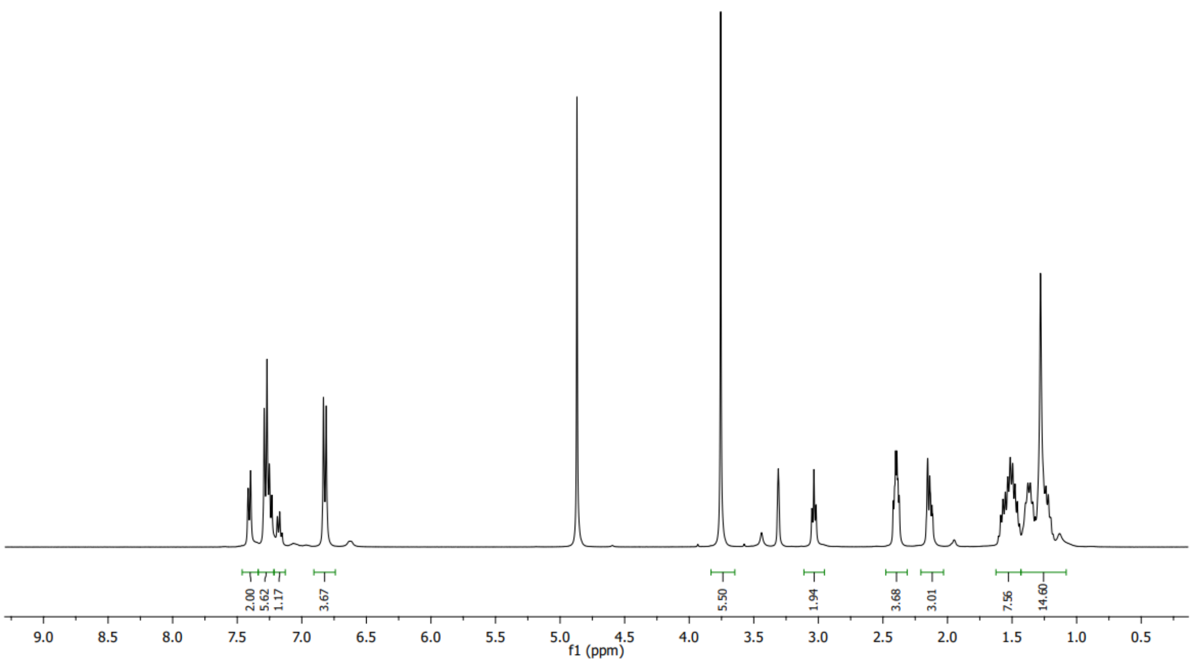


##### *[13]C spectrum of 11*

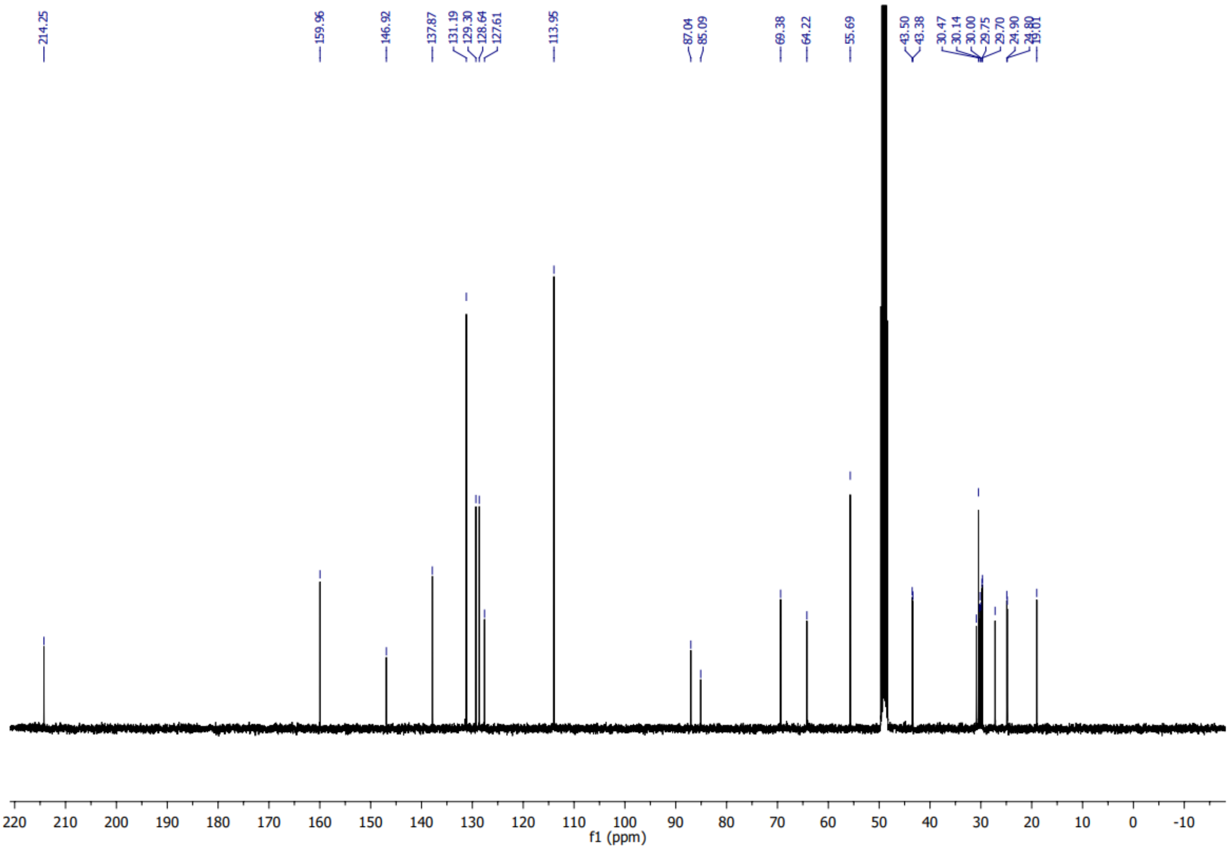

### 1-hydroxyoctadec-17-yn-7-one (12)

##### *[1]H spectrum of 12*

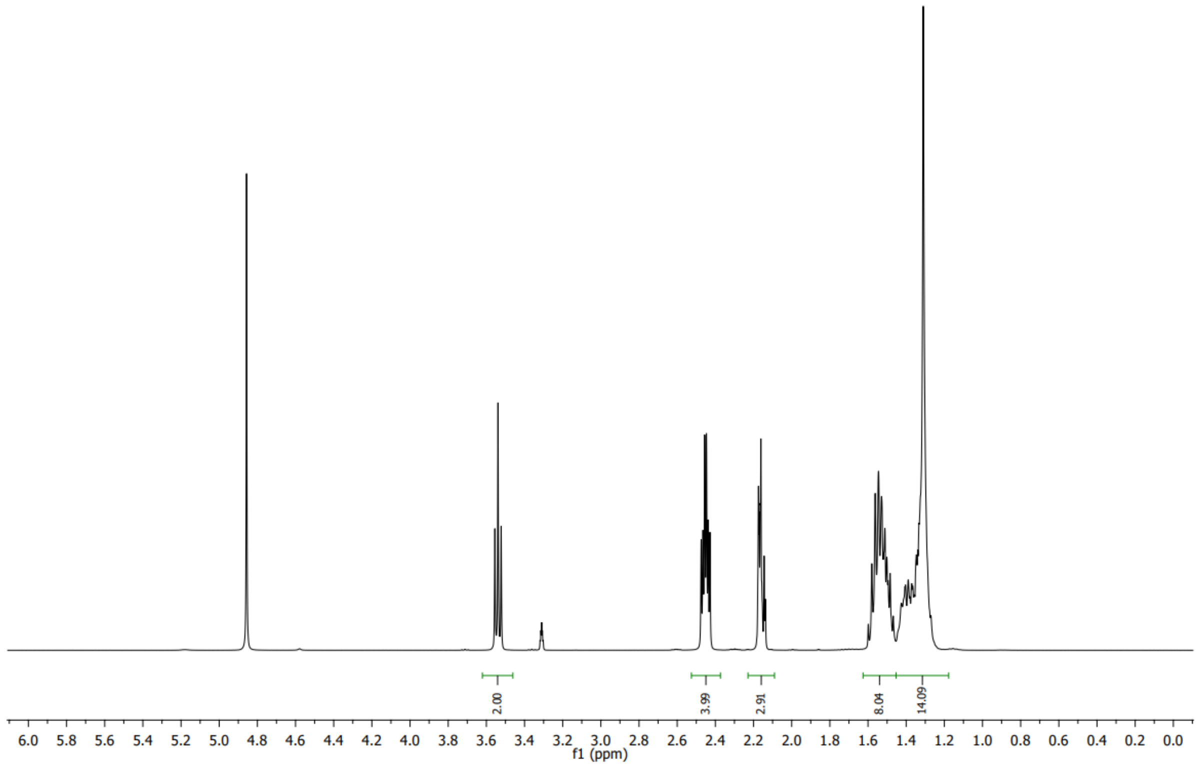


##### *[13]C spectrum of 12*

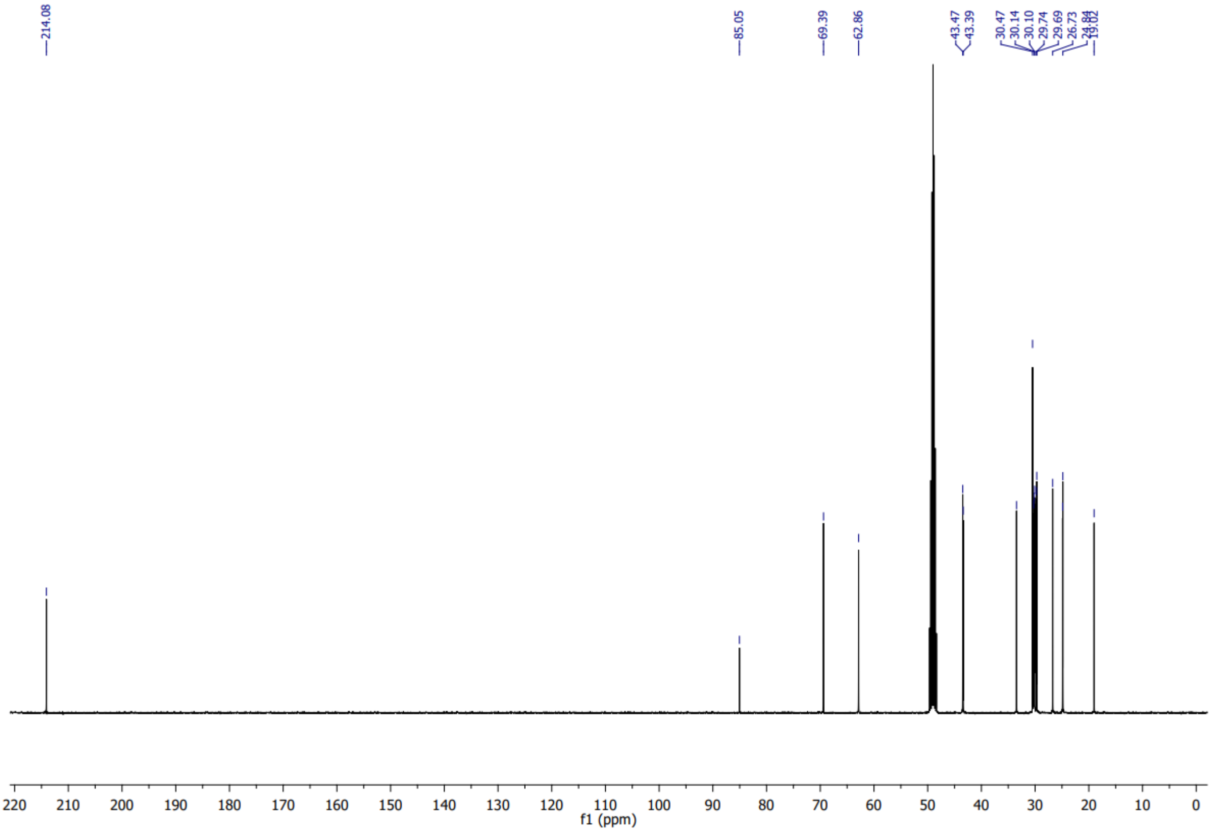

### 6-(3-(undec-10-yn-1-yl)-3*H*-diazirin-3-yl)hexan-1-ol (13)

#### *$^{1}$H spectrum of 13*

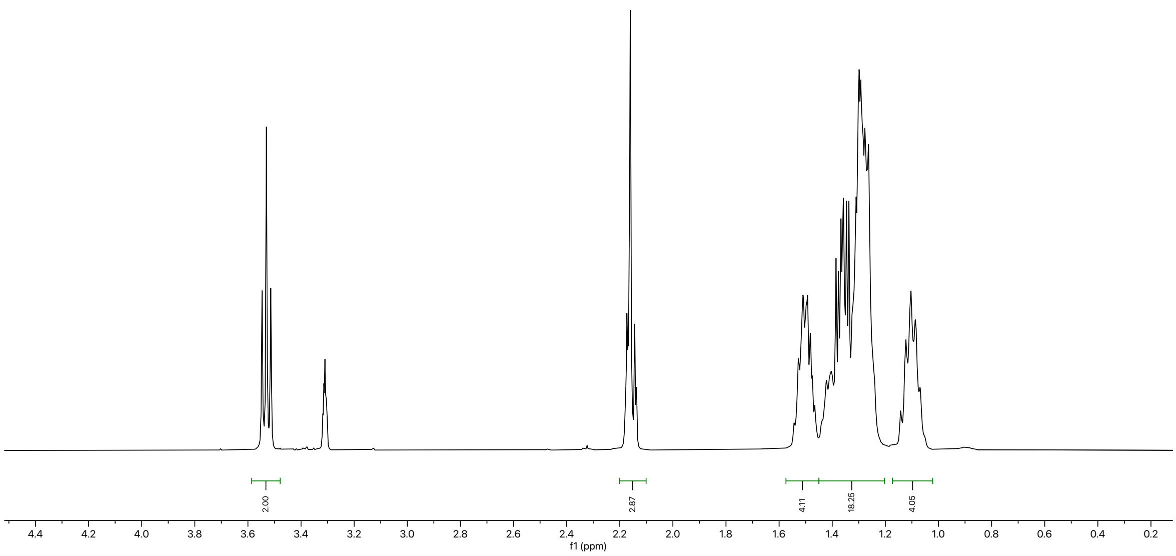


#### *$^{13}$C spectrum of 13*

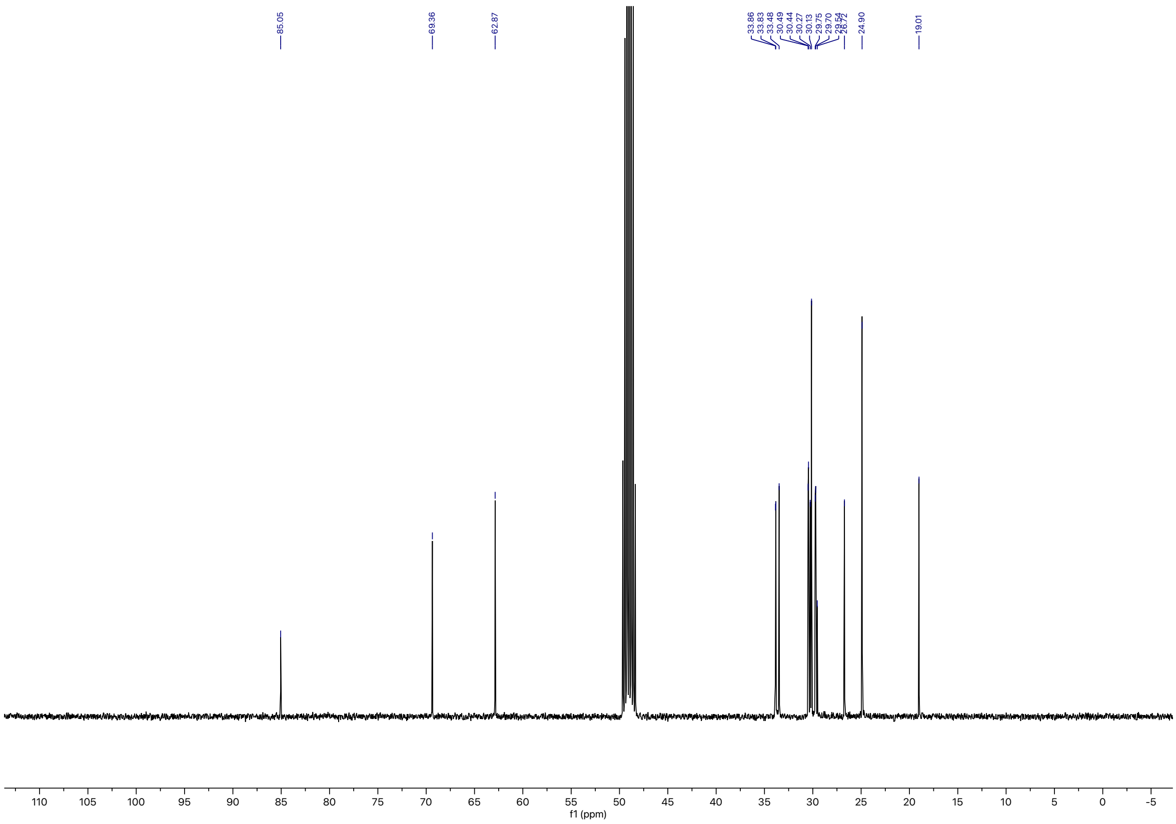

#### (*R*)-1-((triisopropylsilyl)oxy)-3-((6-(3-(undec-10-yn-1-yl)-3*H*-diazirin-3-yl)hexyl)oxy)propan -2-ol (15)

##### *[1]H spectrum of 15*

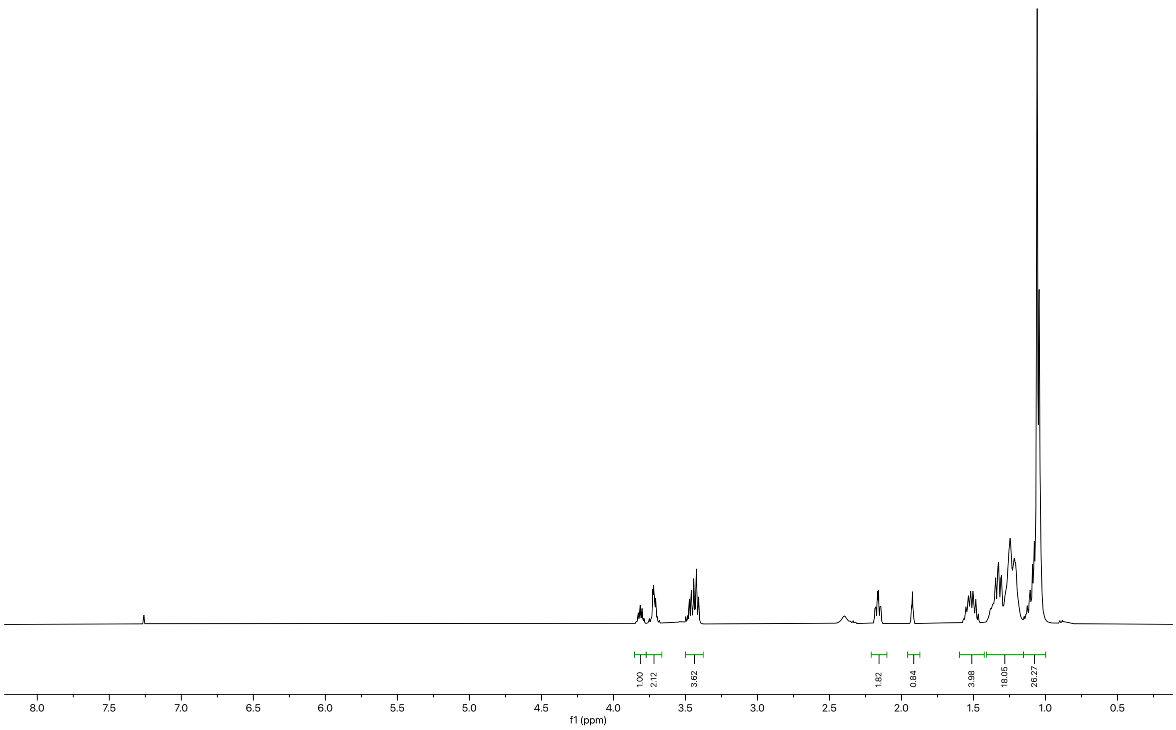

##### *[13]C spectrum of 15*

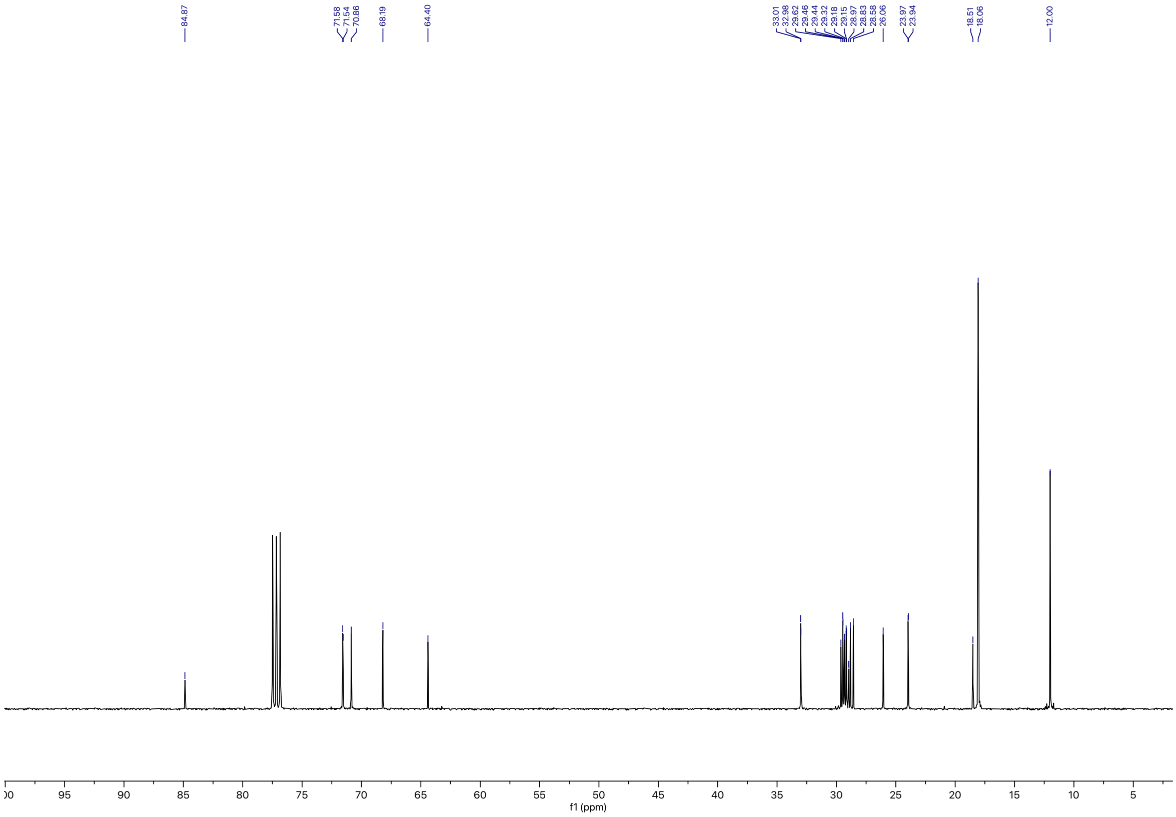

### (*R*)-1-((triisopropylsilyl)oxy)-3-((6-(3-(undec-10-yn-1-yl)-3*H*-diazirin-3-yl)hexyl)oxy)propan-2-yl oleate (16)

#### *$^{1}$H spectrum of 16*

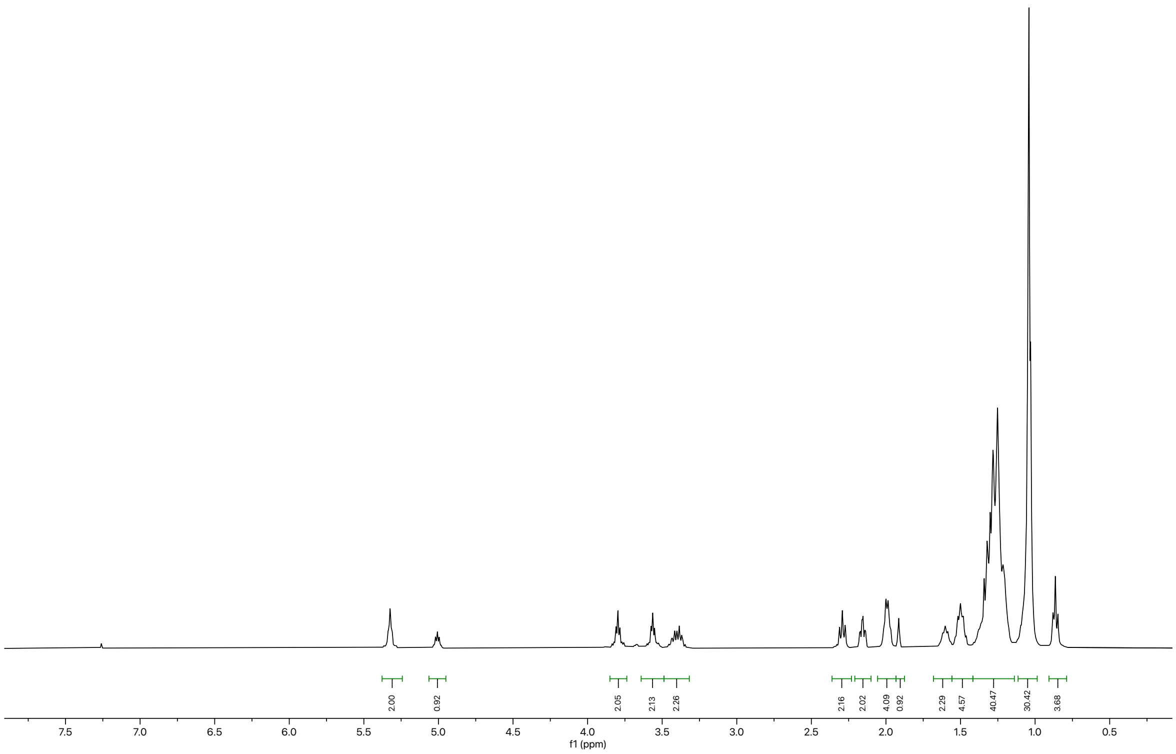

#### *$^{13}$C spectrum of 16*

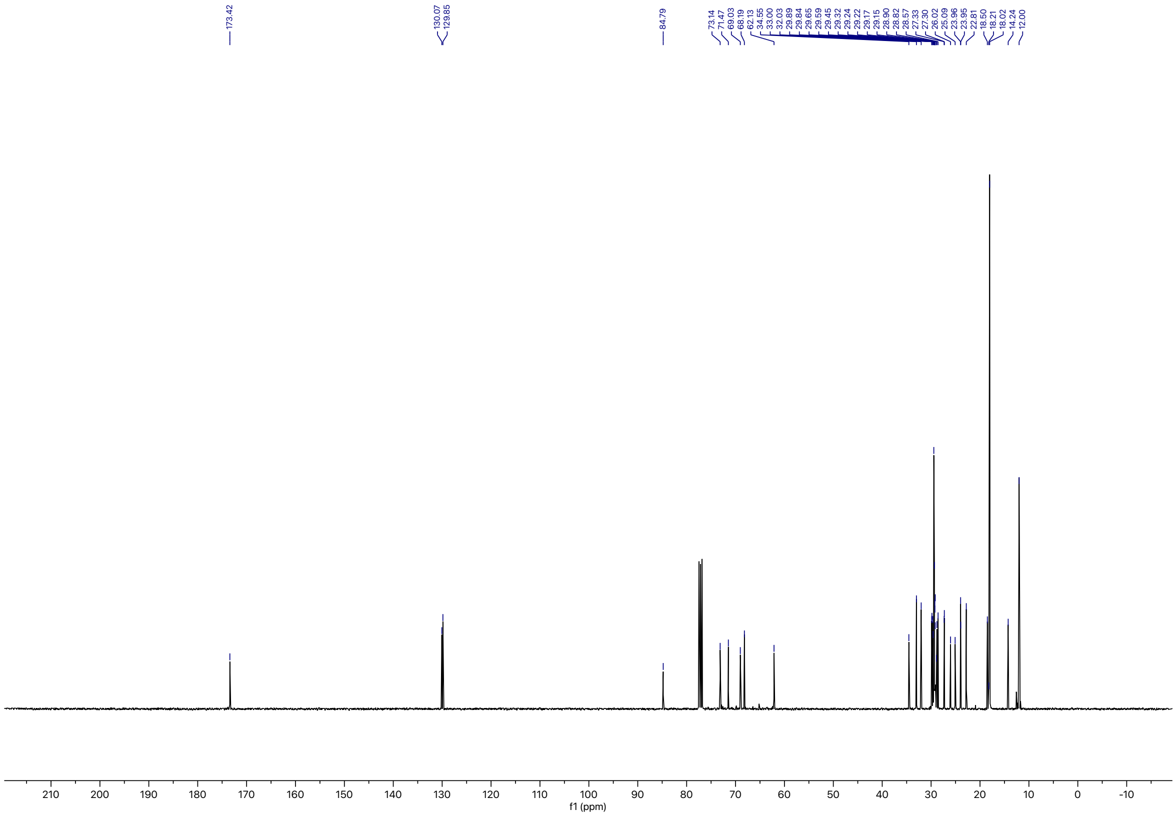

**(*S*)-1-hydroxy-3-((6-(3-(undec-10-yn-1-yl)-3*H*-diazirin-3-yl)hexyl)oxy)propan-2-yl oleate (17)**

***[1]H spectrum of 17***

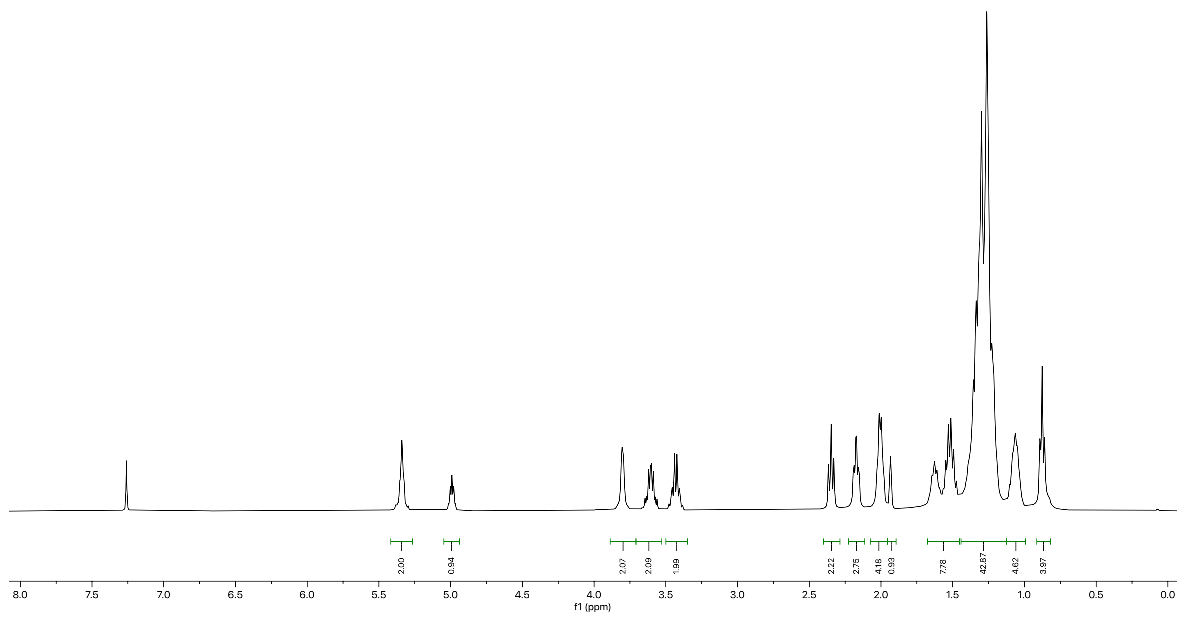

***[13]C spectrum of 17***

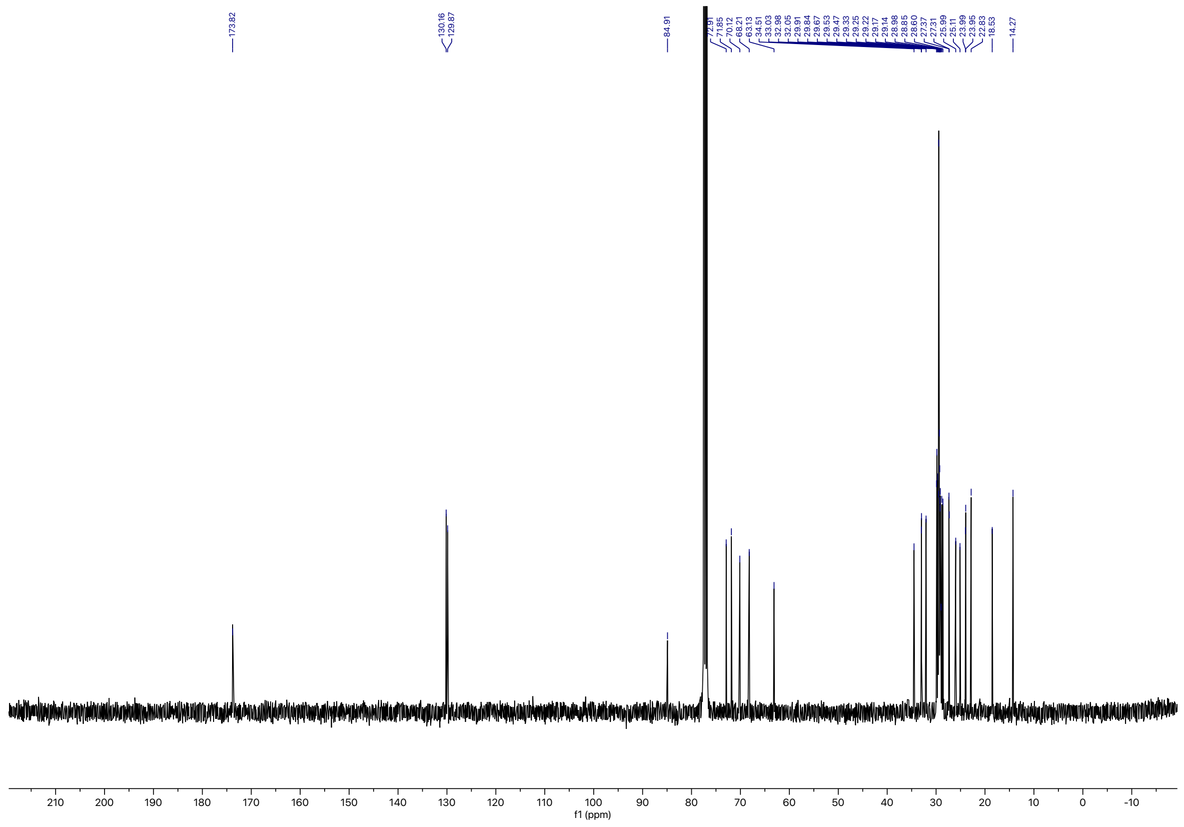

### ePC($Y_{18}$/18:1) (1)

#### *[1]H spectrum of 1*

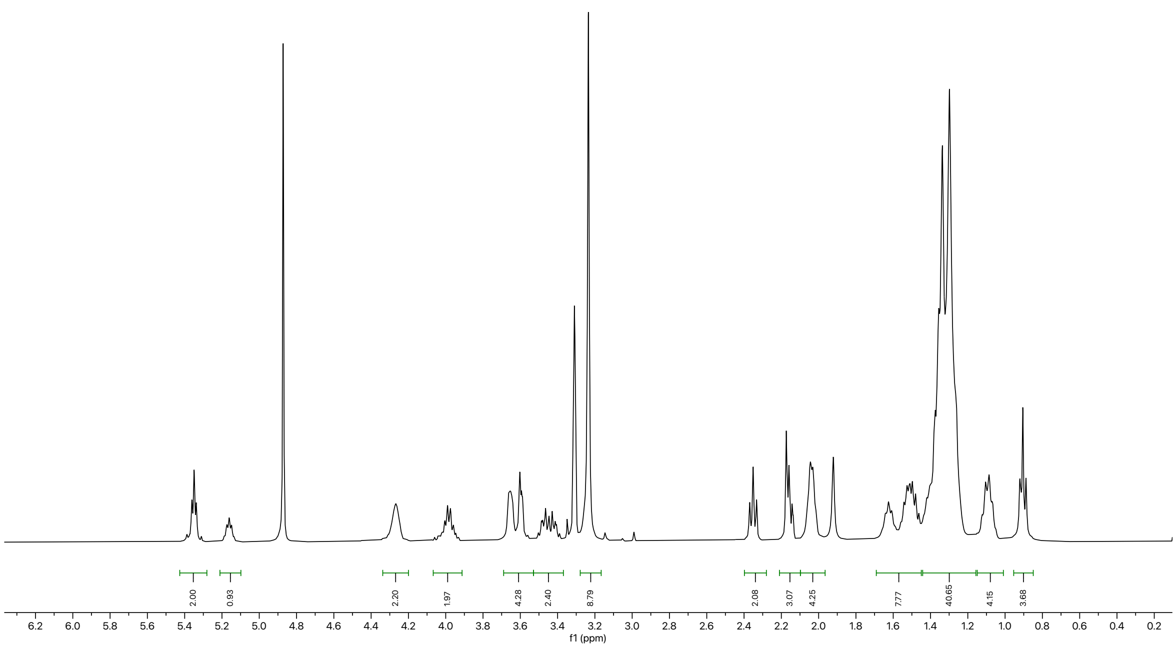

#### *[13]C spectrum of 1*

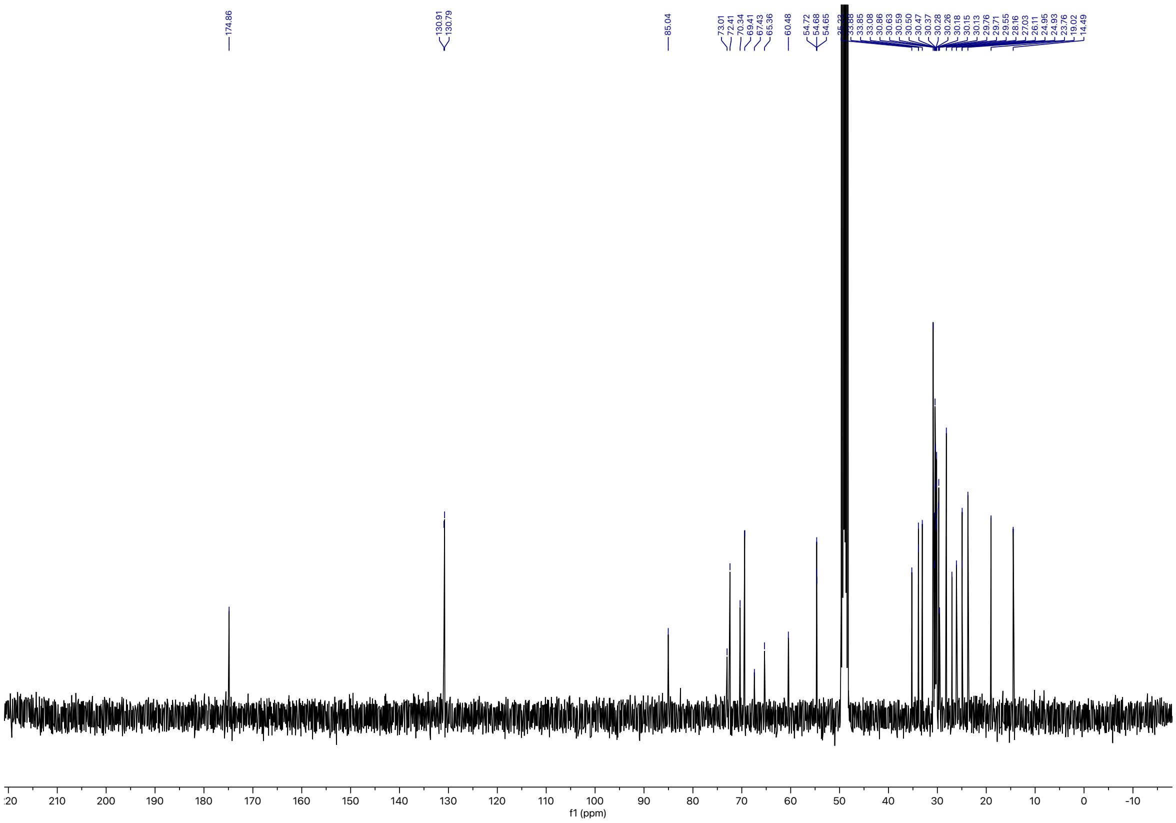

### $^{31}P$ spectrum of 1

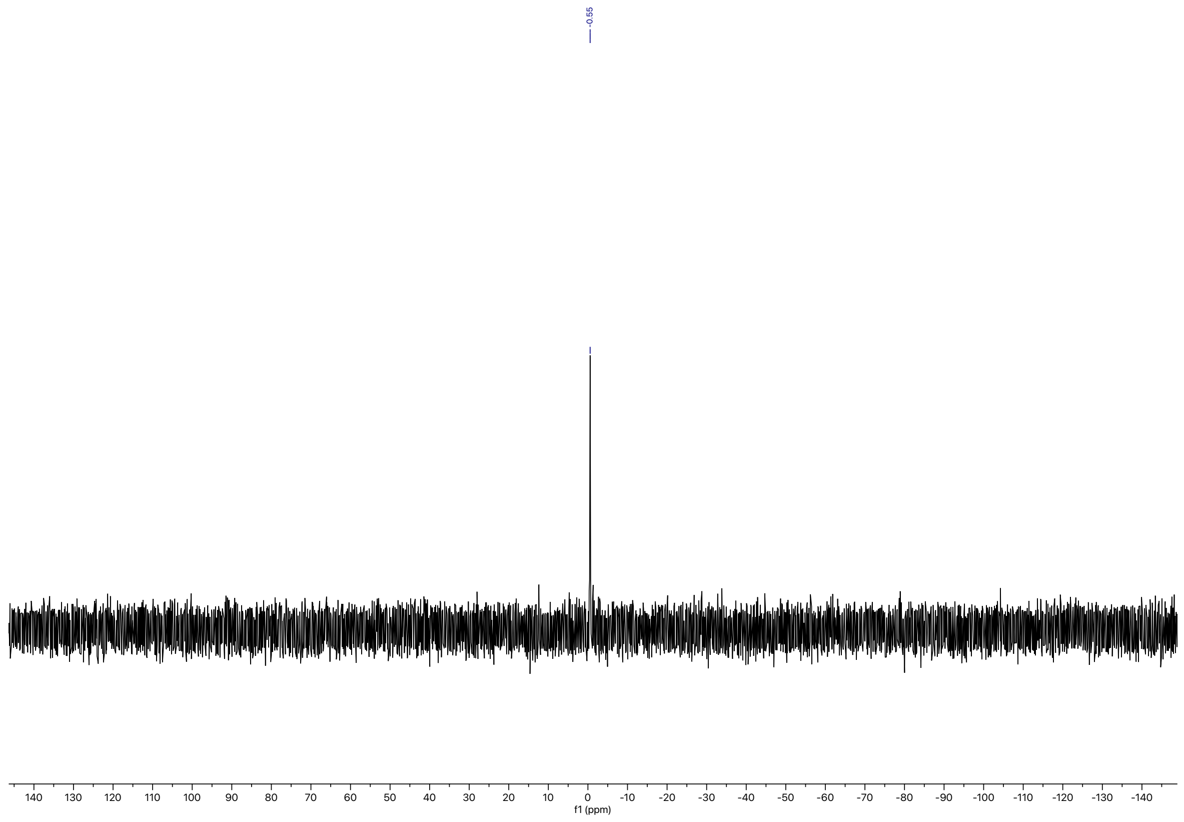

**6-(3-(undec-10-yn-1-yl)-3H-diazirin-3-yl)hexanoic acid (20)**

*[1]H spectrum of 20*

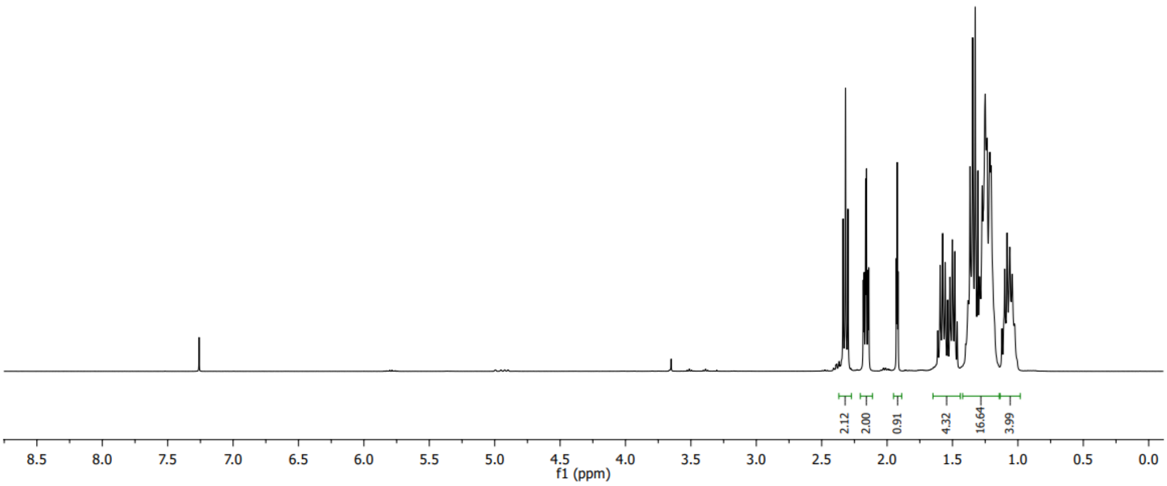

*[13]C spectrum of 20*

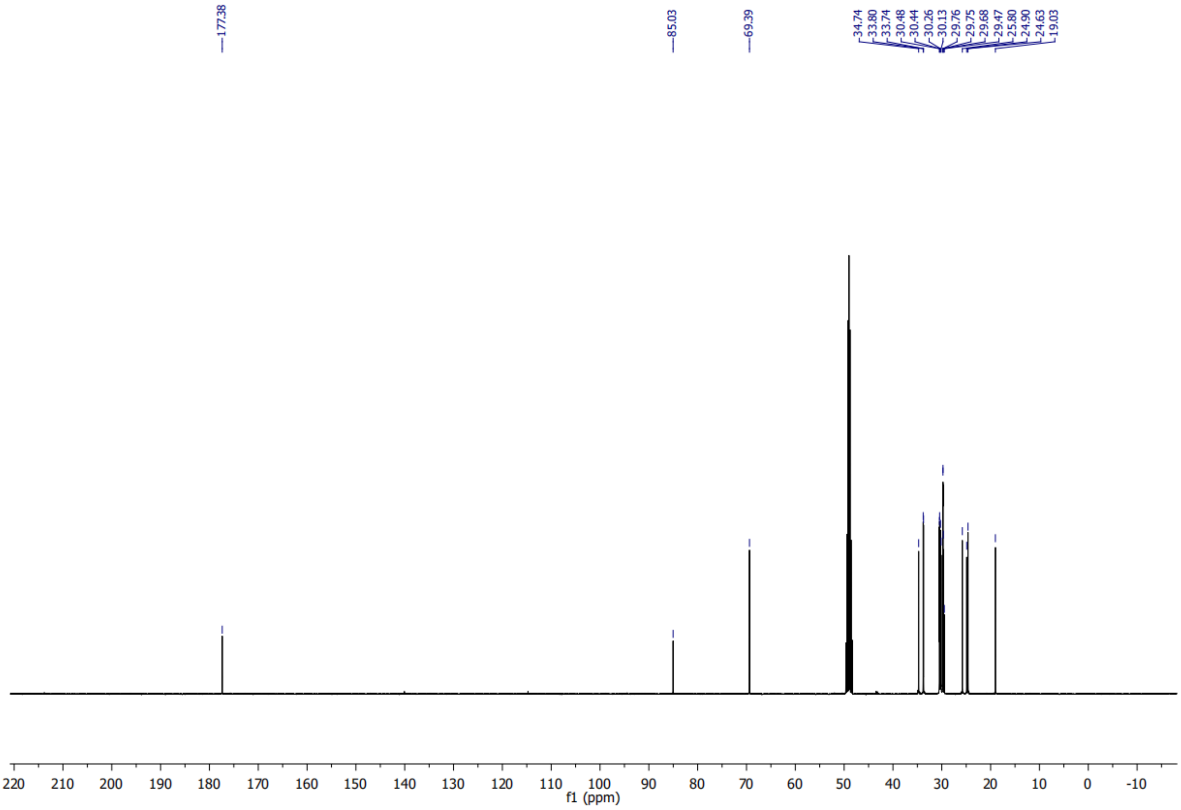

# PC($Y_{18}$/18:1) (2)

*[1]H spectrum of 2*

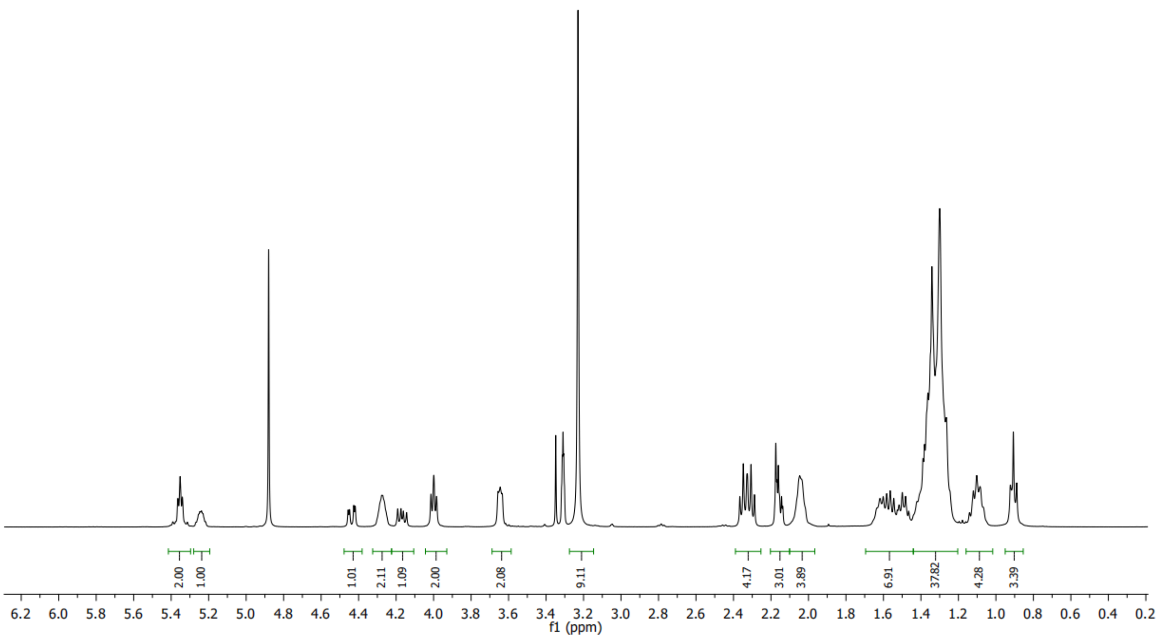


*[13]C spectrum of 2*

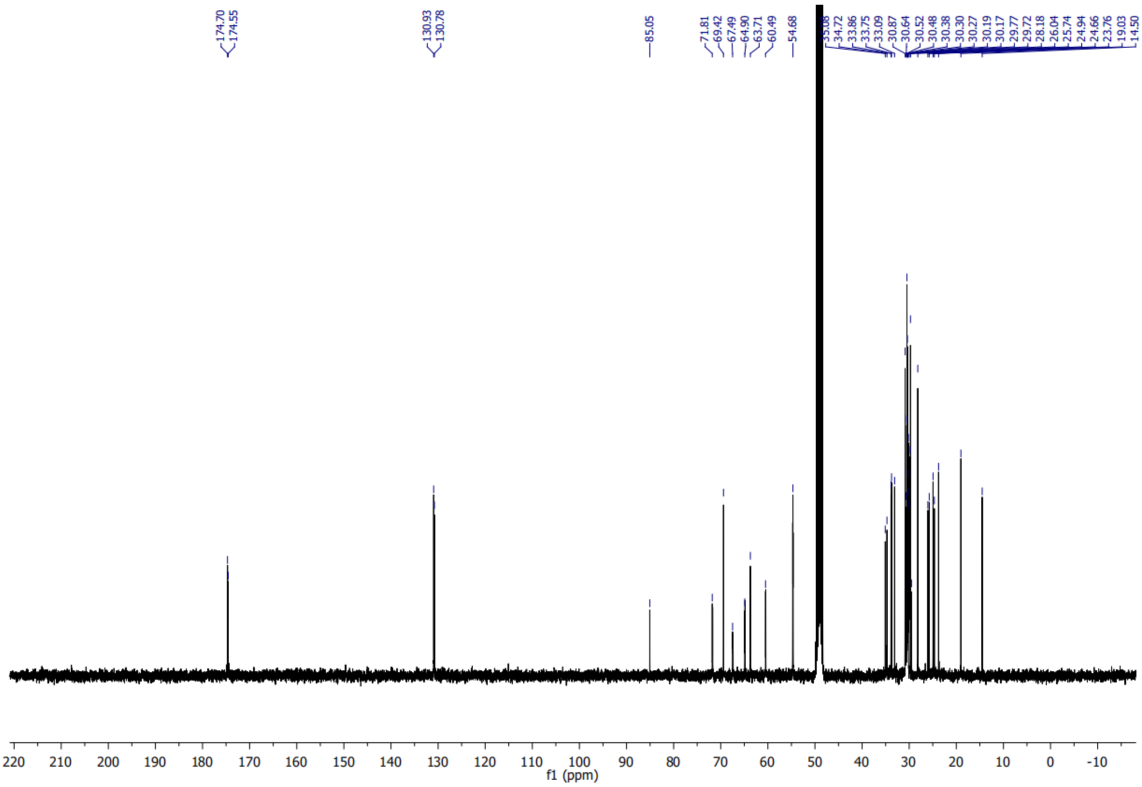

***$^{31}$P spectrum of 2***

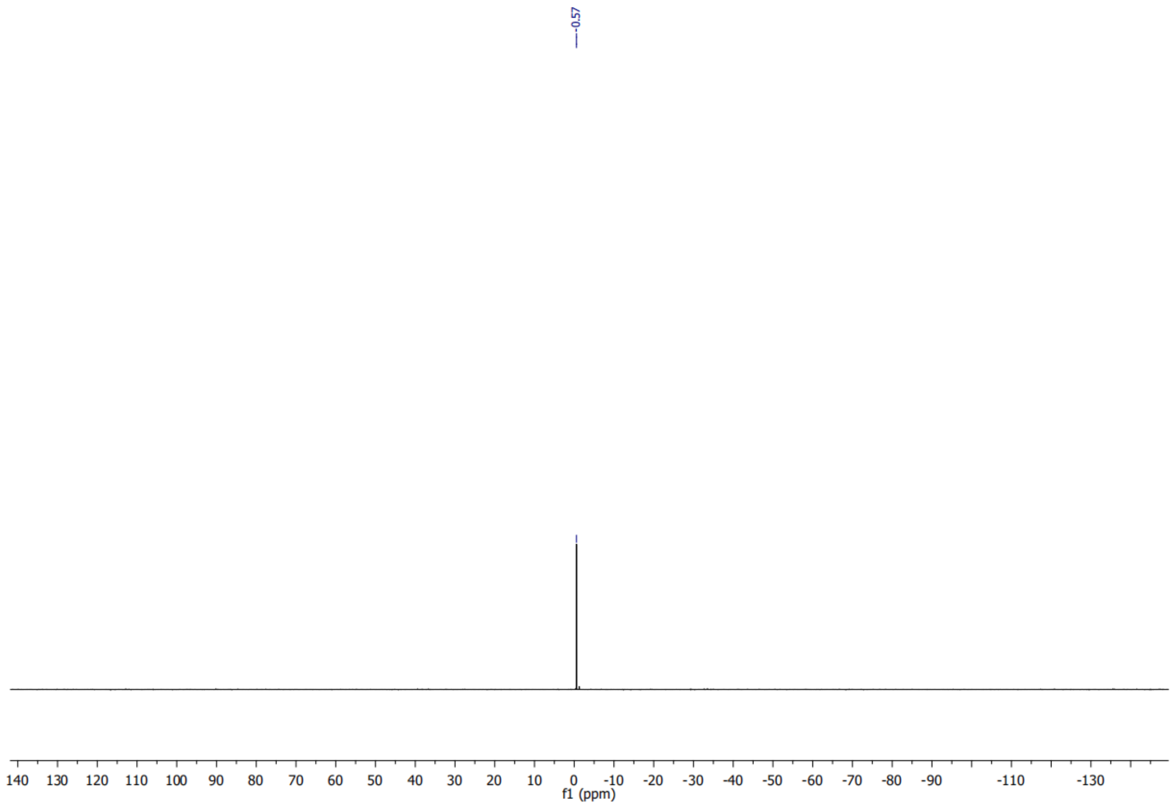

**ePC(18:1/Y$_{16}$) (3)**

***$^{1}$H spectrum of 3***

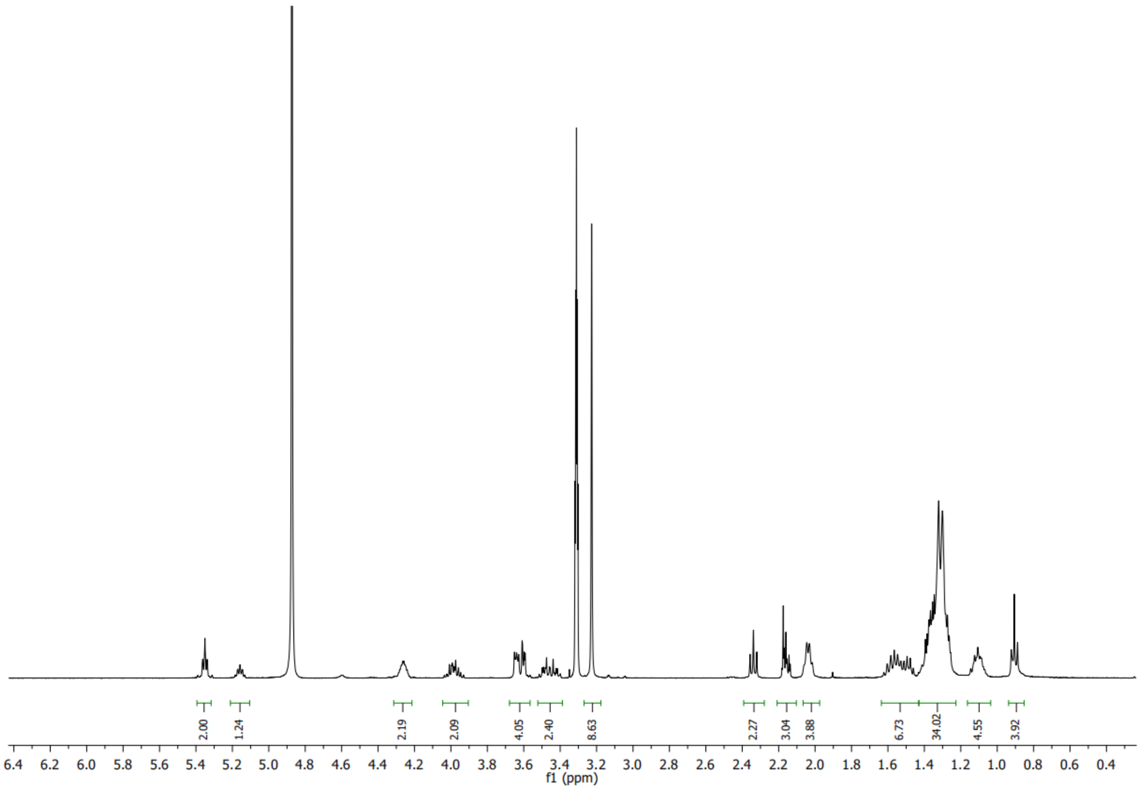

***$^{13}$C spectrum of 3***

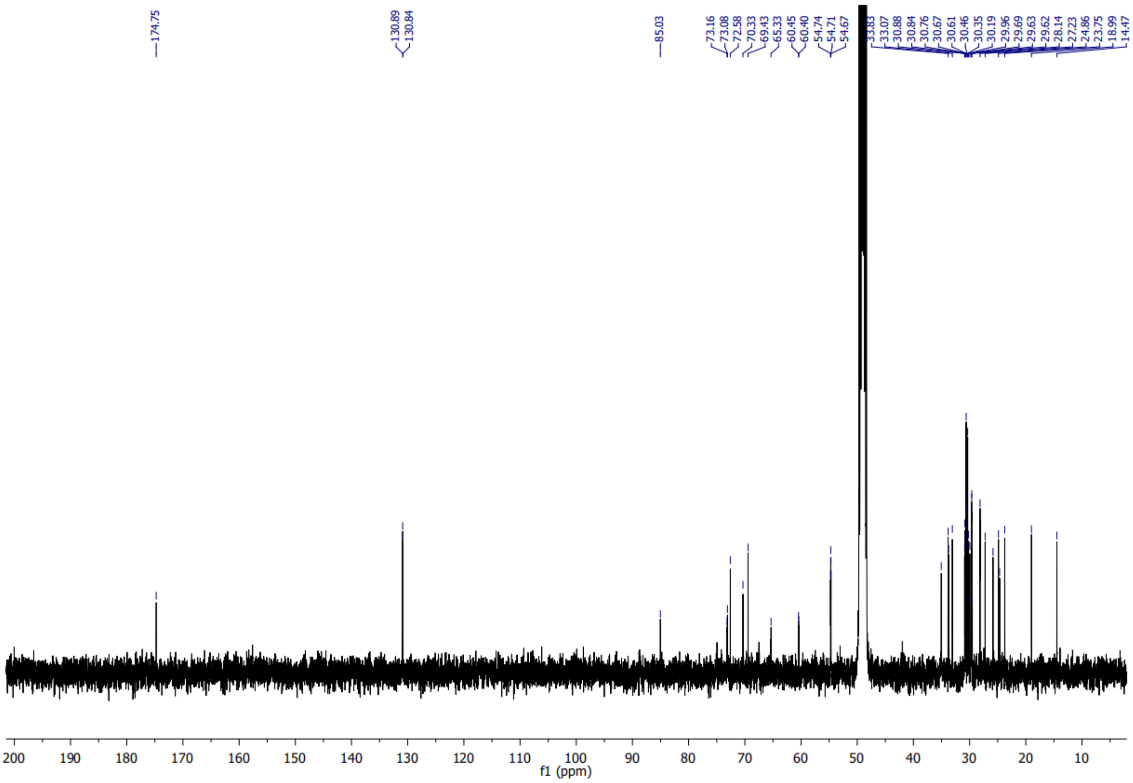

*$^{31}$P spectrum of 3*

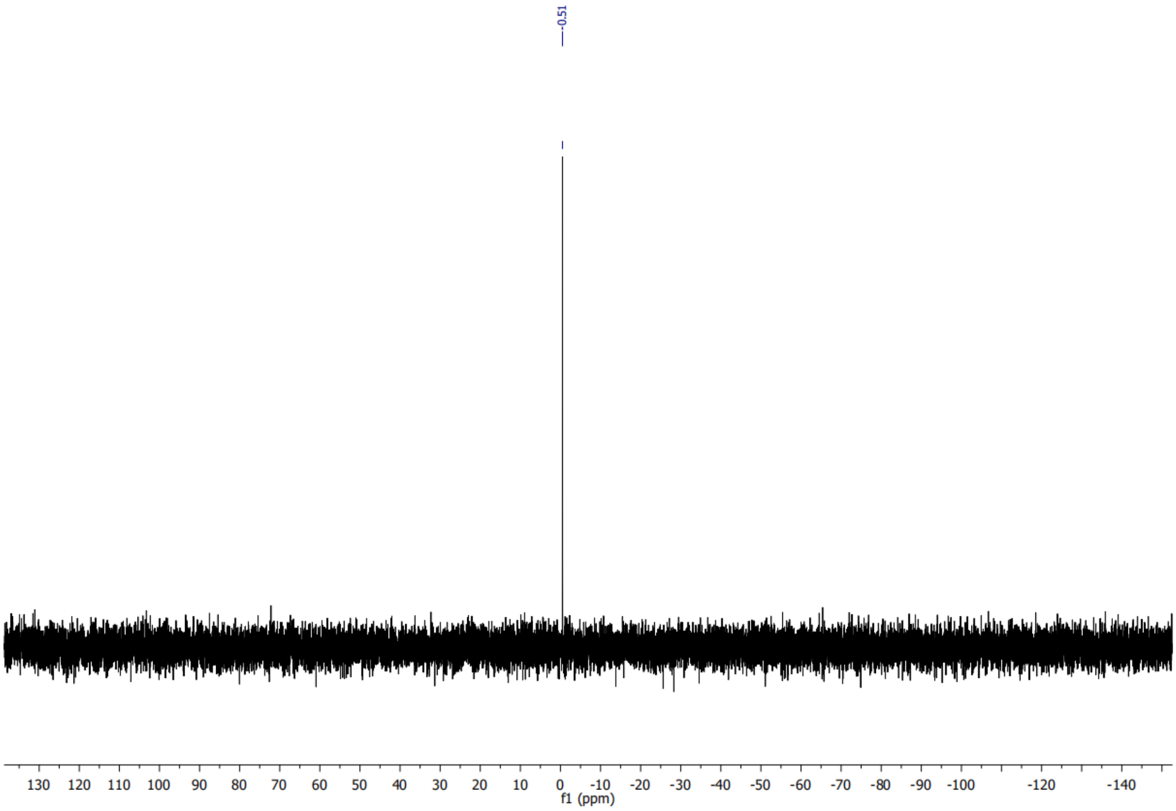

**pPC(18:1/Y$_{16}$) (5)**

***$^{1}$H spectrum of 5***

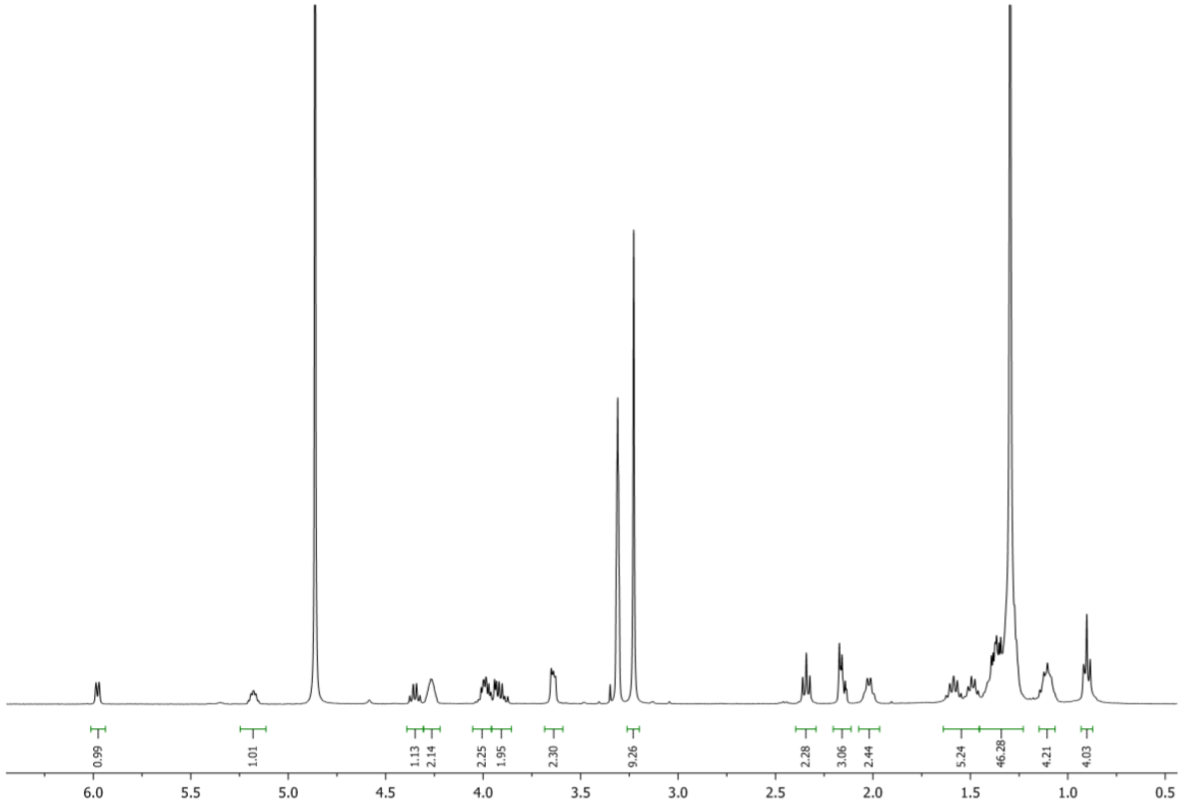


***$^{13}$C spectrum of 5***

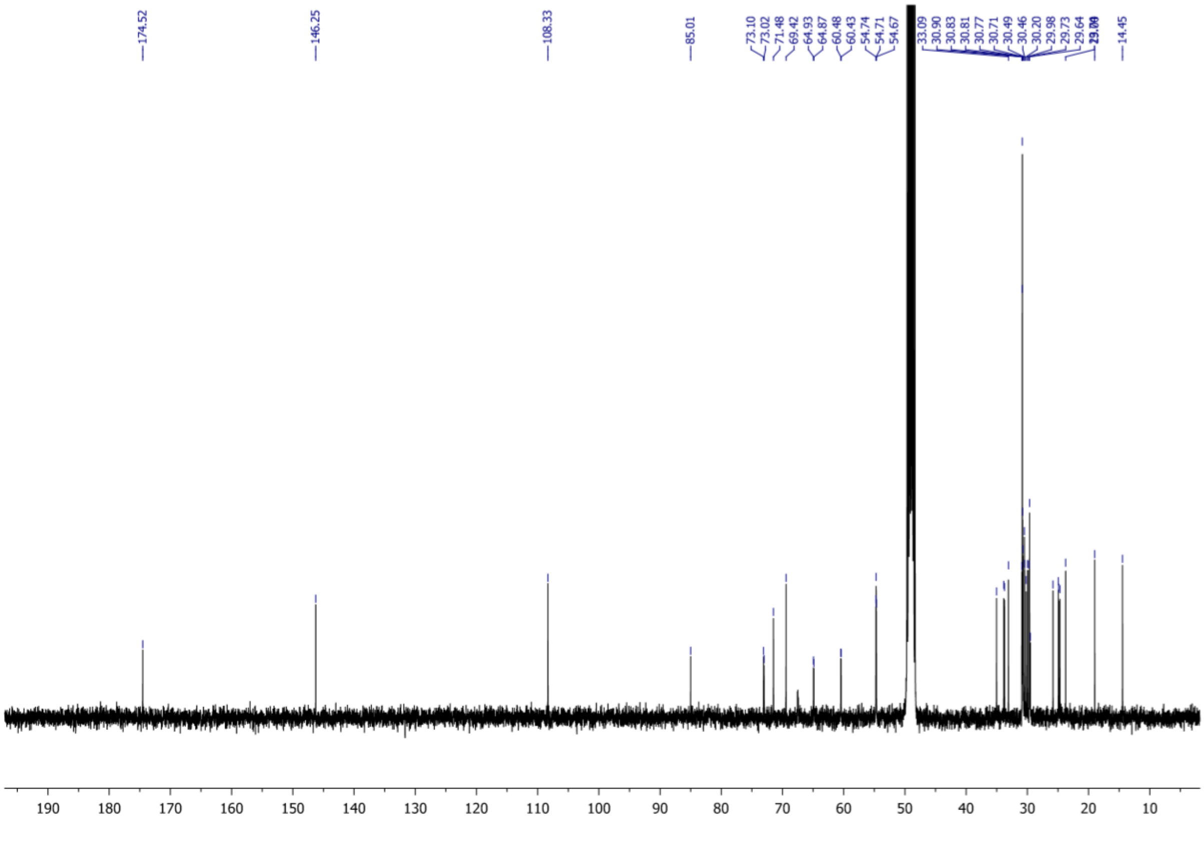

***$^{31}$P spectrum of 5***

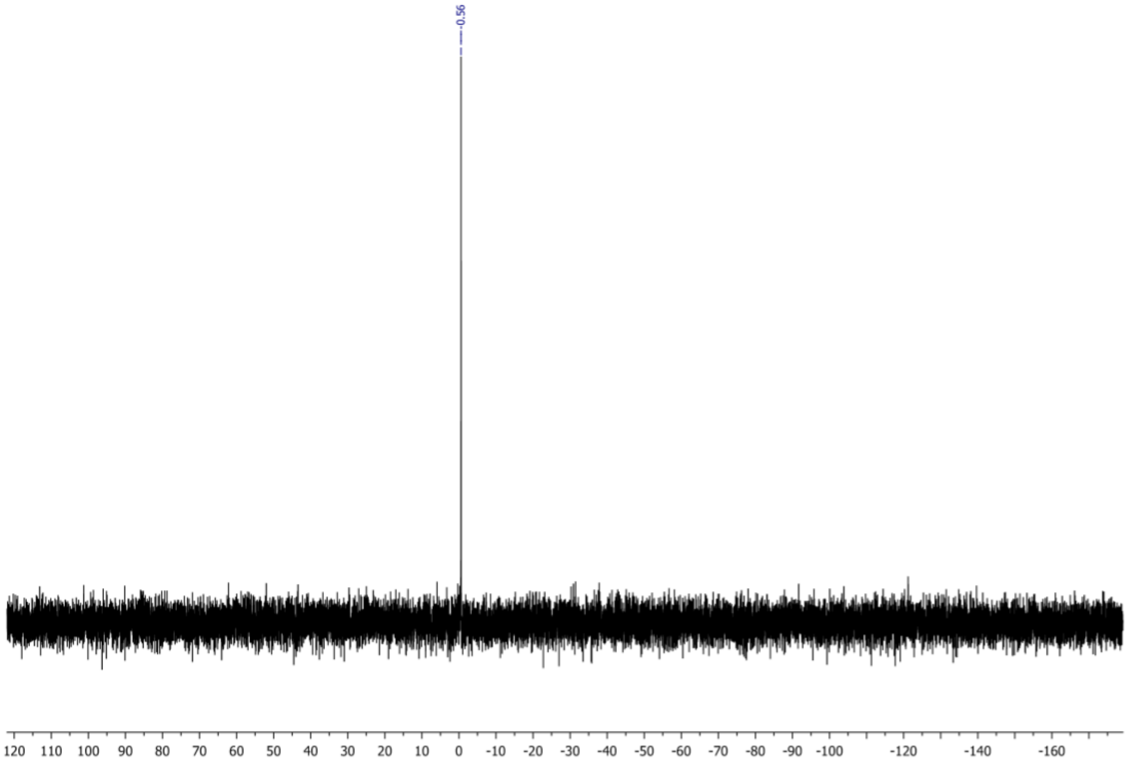
